# Conserved mechanism of nucleoporin regulation of the *Kcnq1ot1* imprinted domain with divergence in embryonic and trophoblast stem cells

**DOI:** 10.1101/430694

**Authors:** Saqib S. Sachani, William A. MacDonald, Ashley M. Foulkrod, Carlee R. White, Liyue Zhang, Mellissa R. W. Mann

## Abstract

Genomic imprinting is an epigenetic phenomenon, whereby dual chromatin states lead to expression of one, and silencing of the other parental allele. Recently, we identified a nucleoporin-mediated mechanism of *Kcnq1ot1* imprinted domain regulation in extraembryonic endoderm stem cells by nucleoporins NUP107, NUP62 and NUP153. Here, we investigate their role in *Kcnq1ot1* imprinted domain regulation in embryonic and trophoblast stem cells. Nucleoporin depletion in both lineages reduced *Kcnq1ot1* noncoding RNA expression and volume, reduced *Kcnq1ot1* paternal domain positioning at the nuclear periphery, and altered histone modifications along with histone modifier enrichment at the imprinting control region. However, while CTCF and cohesin were enriched at nucleoporin binding sites in the imprinting control region in embryonic stem cells, with reduction upon nucleoporin depletion, neither CTCF or cohesin occupied these sites in trophoblast stem cells. Finally, different subsets of silent paternal alleles were reactivated via altered histone modification upon nucleoporin depletion in embryonic and trophoblast stem cells. These results demonstrate a conserved mechanism with divergent regulation of the *Kcnq1ot1* imprinted domain by NUP107, NUP62 and NUP153 in embryonic and extraembryonic lineages.

**Summary Statement:** Investigation of nucleoporins, NUP107, NUP62, and NUP153, revealed a conserved nucleoporin-dependent mechanism that mediates *Kcnq1ot1* imprinted domain regulation in ES and TS cells, although notable lineage-specific divergence was also observed.

## INTRODUCTION

Genomic imprinting is an epigenetic process that regulates parental-specific transcription. Dual chromatin states are generated at master control switches, called imprinting control regions (ICRs) during oogenesis or spermatogenesis. These gametic imprinting marks at ICRs, which includes differential DNA methylation, govern the expression of neighboring imprinted genes in a domain during embryonic development such that one parental allele is expressed while the other parental allele is silenced (Barlow and Bartolomei, 2014). However, it is still not clear when and how differential parental-allelic expression is established from the ICR to genes across an imprinted domain. What is known is that disruptions in imprinted gene regulation at imprinted domains cause aberrant growth and development phenotypes, as well as imprinting disorders. For example, Beckwith-Wiedemann Syndrome (BWS) is an imprinting disorder caused by epigenetic defects at the *KCNQ1OT1* domain, which leads to pre- and post-natal overgrowth (Weksberg et al., 2010).

The *Kcnq1ot1/KCNQ1OT1 (Kcnq1 opposite transcript 1)* imprinting domain harbors one paternally expressed *Kcnq1ot1* noncoding RNA (ncRNA), as well as sixteen protein-coding genes, of which nine are maternally-expressed, while the other genes escape imprint regulation (Barlow and Bartolomei, 2014). On the maternal allele, the *Kcnq1ot1* ICR is methylated, which silences the embedded *Kcnq1ot1* ncRNA promoter, thereby permitting maternal allelic expression of flanking imprinted genes (Golding et al., 2011; Jones et al., 2011; Lewis et al., 2006; Pandey et al., 2008; Redrup et al., 2009). On the paternal allele, the *Kcnq1ot1* ICR is unmethylated, and the *Kcnq1ot1* ncRNA is transcribed, which results in paternal allelic repression of imprinted genes in the domain.

The *Kcnq1ot1* domain exhibits both tissue-specific and developmental stage-specific imprinted gene regulation. This has led to imprinted genes in the domain being classified by their expression during mid-gestation development. Genes that reside closer to the *Kcnq1ot1* ICR exhibit paternal allelic silencing in embryonic and placental tissues, and are designated as inner/ubiquitously imprinted genes, whereas genes located more distal from the *Kcnq1ot1* ICR, which possess paternal allelic silencing in the placenta, are classified as outer/placental-specific imprinted genes (Lewis et al. 2004; Umlauf et al. 2004). Currently, it is unclear when embryonic and extraembryonic lineages acquire differential regulation across an imprinted domain during early stages of development.

During preimplantation development, three distinct cell lineages emerge, such that at the early blastocyst stage, there are epiblast precursor, trophectoderm and primitive endoderm cells. These cells will give rise to the fetus, placenta and yolk sac, respectively (Fig. S1). Pluripotent stem cells can be derived from these three lineages to produce embryonic (ES), trophoblast (TS) and extraembryonic endoderm (XEN) stem cells. At this early developmental time point, ES, TS and XEN stem cells have distinct imprinted expression patterns for genes in the *Kcnq1ot1* imprinted domain (Golding et al., 2011; Lewis et al., 2004; Lewis et al., 2006; Terranova et al., 2008; Umlauf et al., 2004), suggesting that the *Kcnq1ot1* imprinted domain may be differentially regulated even at early stages of lineage differentiation. However, the mechanisms responsible for this differential imprinted expression between the three lineages are still not clear.

In a previous study, we investigated the role of several nucleoporins in *Kcnq1ot1* imprinted domain regulation (Sachani et al., 2018). Nucleoporins comprise the nuclear pore complex, which are large aqueous transport channels embedded within the nuclear membrane. One key function of nuclear pore complexes is transport of cellular material between the cytoplasm and nucleus (Hoelz et al., 2011). However, in recent years, it has become apparent that nuclear transport is not the sole function of the nuclear pore complex/nucleoporins. Previous studies have identified roles for nucleoporins in gene regulation in yeast, *Drosophila*, and mammalian cells, although the mechanisms by which nucleoporins govern this regulation is not fully understood (Capelson et al., 2010; Jacinto et al., 2015; Kalverda et al., 2010; Light et al., 2013; Mendjan et al., 2006; Vaquerizas et al., 2010). Our investigation of nucleoporins, NUP107, NUP62, and NUP153, revealed a nucleoporin-dependent imprinting regulatory mechanism that mediates *Kcnq1ot1* imprinted domain regulation in XEN cells, without impairment of nuclear-cytoplasmic transport function (Sachani et al., 2018). More specifically, our data showed that depletion of *Nup107, Nup62* and *Nup153* in mouse XEN cells led to a reduction in *Kcnq1ot1* noncoding RNA expression and its volume at the paternal *Kcnq1ot1* domain; a shift in positioning of paternal *Kcnq1ot1* domain away from the nuclear periphery; and reactivation of a subset of normally silent paternal alleles of protein-coding genes in the domain. While DNA methylation at the *Kcnq1ot1* ICR was not altered, *Nup107, Nup62* and *Nup153* depletion altered active and repressive histone modifications and reduced cohesin interactions at the *Kcnq1ot1* ICR. These alterations were not observed upon *Nup98/96* depletion. Furthermore, they were not related to nuclear-cytoplasmic transport.

Here, we hypothesized that NUP107, NUP62 and NUP153, but not NUP98, play a regulatory role at the *Kcnq1ot1* imprinted domain in ES and TS cells, and may contribute to the differential regulation of the *Kcnq1ot1* imprinted domain between ES, TS and XEN cells. In the current study, we identified a conserved mechanism of nucleoporin regulation of the *Kcnq1ot1* imprinted domain in ES and TS cells, as previously identified in XEN cells (Sachani et al., 2018). NUP107, NUP62 and NUP153 bind to the *Kcnq1ot1* ICR on each side of the *Kcnq1ot1* ncRNA promoter, and are required for paternal *Kcnq1ot1* ncRNA expression, positioning of the paternal *Kcnq1ot1* domain at the nuclear periphery, as well as recruitment of histone modifiers at the paternal *Kcnq1ot1* ICR in both ES and TS cells. Divergence in NUP107, NUP62 and NUP153 regulation at the *Kcnq1ot1* imprinted domain was also found between the three cell lineages. Firstly, nucleoporins bind promoters of imprinted genes in a lineage-specific manner, leading to paternal allelic silencing of different subsets of imprinted, protein-coding genes in the *Kcnq1ot1* imprinted domain. Secondly, while NUP107, NUP62 and NUP153 are required for CTCF and cohesin localization at the *Kcnq1ot1* ICR in ES cells, TS cells lack CTCF and cohesin occupancy at the *Kcnq1ot1* ICR. Consistent with our hypothesis, NUP98 had a limited role at the *Kcnq1ot1* imprinted domain in all three lineages. Overall, our results demonstrated a conserved mechanism with divergent regulation of the *Kcnq1ot1* imprinted domain by NUP107, NUP62 and NUP153 in embryonic and extraembryonic lineages as early as the blastocyst stage.

## RESULTS

### *Nup107, Nup62 and Nup153* depletion alter *Kcnq1ot1* ncRNA expression in ES and TS cells

Wild type C57BL/6J X *Mus musculus castaneus* ES and TS cells were transfected with siRNAs targeting *Nup107, Nup62, Nup98/96* or *Nup153* to produce RNA and protein depletion (Fig. S2, S3). Nucleoporin-depleted cells were then assessed for total *Kcnq1ot1* ncRNA expression levels. Compared to control cells, *Nup107* and *Nup62* depletion significantly decreased *Kcnq1ot1* ncRNA levels in both ES (0.34 and 0.42 of controls) and TS (0.40 and 0.49 of controls) cells (Fig. 1A). By comparison, no significant difference was observed between control and *Nup98/96*- or *Nup153*-depleted ES and TS cells (1.03 and 1.09; 0.75 and 0.86, respectively). To independently determine the effects of nucleoporin depletion on maternal and paternal *Kcnq1ot1* ncRNA absolute transcript abundance, we used a highly sensitive, precision method of droplet digital PCR with FAM and HEX strain-specific probes (Sachani et al., 2018) (Fig. 1B). Compared to control cells, *Nup107, Nup62* and *Nup153* depletion resulted in significantly reduced paternal *Kcnq1ot1* transcripts by 2,326-2,635 copies in ES cells, and 3,001–4,340 copies in TS cells (Fig. 1B). The low number of maternal *Kcnq1ot1* transcripts did not change in *Nup107*- and *Nup62*-*depleted* ES and TS cells. However, *Nup153* depletion significantly increased maternal *Kcnq1ot1* transcripts by 2,226 copies in ES cells and 3,883 copies in TS cells. By comparison, absolute transcript abundance of maternal and paternal *Kcnq1ot1* ncRNA copies upon *Nup98/96* depletion was indistinguishable from control ES and TS cells. These data are consistent with total *Kcnq1ot1* ncRNA expression levels observed in *Nup*-depleted cells. Overall, these results were similar to those reported in XEN cells, except for *Nup98/96* depletion, which produced significantly increased paternal *Kcnq1ot1* ncRNA copies (Sachani et al., 2018).

**Figure 1:**
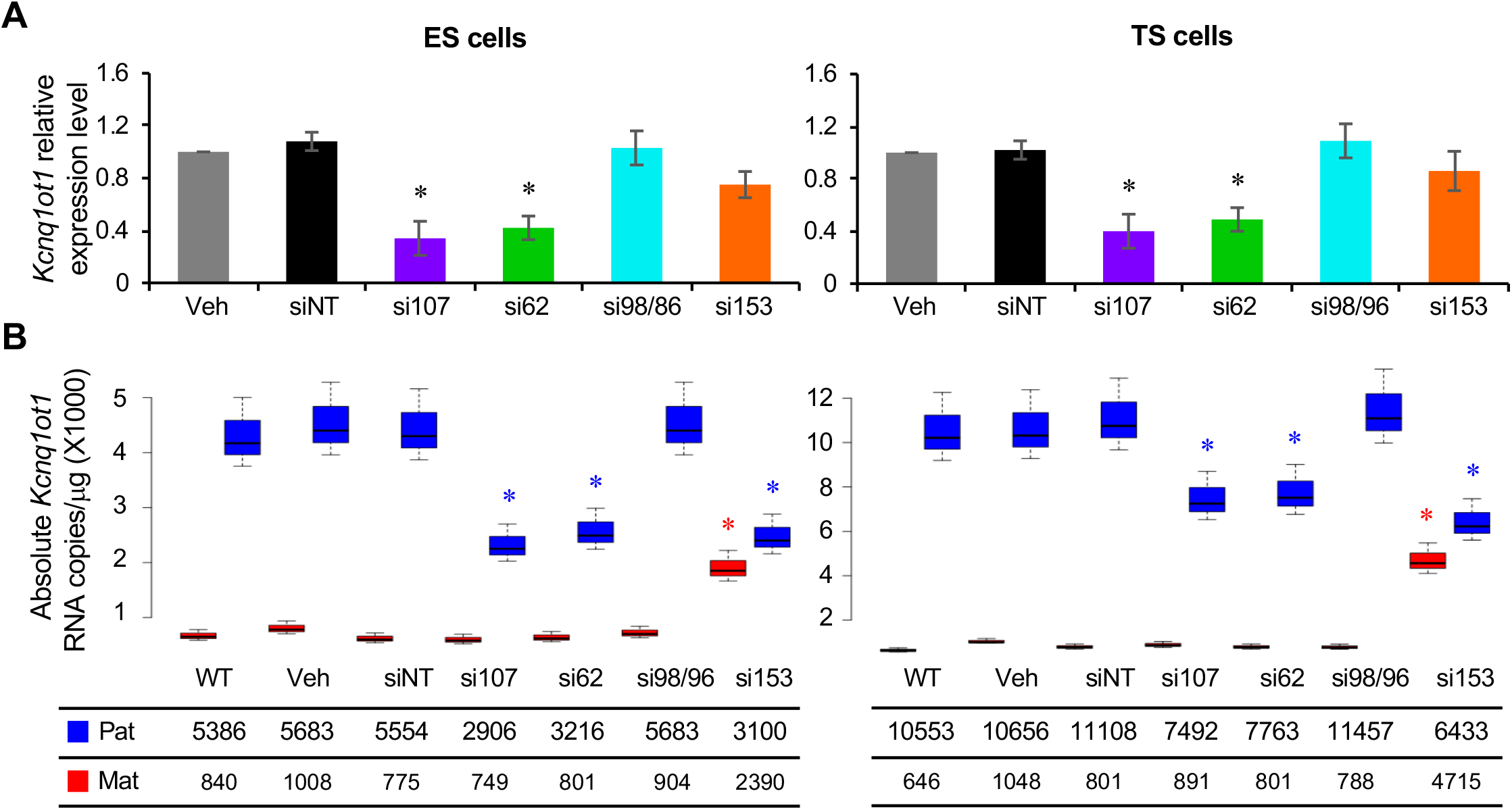
Nucleoporin depletion disrupted *Kcnq1ot1* ncRNA expression in ES and TS cells. (A) Real-time *Kcnq1ot*1 ncRNA expression levels in control and *Nup*-depleted ES and TS cells (normalized to *Gapdh;* n=3 biological samples with 3 technical replicates per sample). (B) Absolute allelic *Kcnq1ot1* transcript abundance in control and *Nup*-depleted ES and TS cells (n=3 biological samples). Paternal, blue; maternal, red; *, significance p < G.G1 compared to siNT control; error bars, s.e.m.; WT, wildtype; Veh, vehicle; siNT, non-targeting siRNA; si1G7, *Nup107* siRNA; si62, *Nup62* siRNA; si98/96, *Nup9S/96* siRNA; si153, *Nup153* siRNA.

### NUP107, NUP62 and NUP153 regulate *Kcnq1ot1* ncRNA volume in ES and TS cells

Within the nucleus, the *Kcnq1ot1* ncRNA overlaps with the three-dimensional structure of the paternal *Kcnq1ot1* domain, which can be calculated as the RNA signal volume (Fedoriw et al., 2012; Pandey et al., 2008; Redrup et al., 2009; Terranova et al., 2008; Sachani et al., 2018). To determine whether *Kcnq1ot1* ncRNA volume was altered in ES and TS cells, *Kcnq1ot1* ncRNA volume was calculated following *Kcnq1ot1* RNA/DNA FISH on G1-synchronized control and *Nup*-depleted ES and TS cells (Fig. 2A). With respect to the paternal *Kcnq1ot1* ncRNA volume, ES and TS controls had 8–10% and 10–12% of cells with low (<0.7 μm^3^); 69–73% and 70–79% of cells with medium (0.7–1.4 μm^3^); 13–16% and 8–16% of cells with high (1.4–2.1 μm^3^); and 5–7% and 2–4% of cells with very high (1.4–2.1 μm^3^) volumes, respectively (Fig. 2B, C). By comparison, upon *Nup107, Nup62* and *Nup153* depletion, a significant increase was observed in ES and TS cells with low paternal *Kcnq1ot1* ncRNA volumes *(Nup107*, 59% 60%; *Nup62*, 65% and 56%; *Nup153*, 26% and 88%, respectively), with a corresponding decrease in cells with medium to very high signal volume. Furthermore, 29% of ES cells and 58% of TS cells gained maternal *Kcnq1ot1* ncRNA signal, of which 68% and 79%, respectively, had low *Kcnq1ot1* ncRNA volumes (Fig 2B, C). *Nup98/96* depletion did not significantly alter the *Kcnq1ot1* ncRNA volume in ES or TS cells. Saliently, alterations in transcript abundance and volume were not a result of altered stability of the *Kcnq1ot1* ncRNA transcript upon *Nup*-depletion in ES and TS cells (Fig. S4A, B). Overall, these results were similar to those for XEN cells, except for *Nup98/96* depletion, which produced a significant increase in paternal *Kcnq1ot1* ncRNA volumes (Sachani et al., 2018).

**Figure 2:**
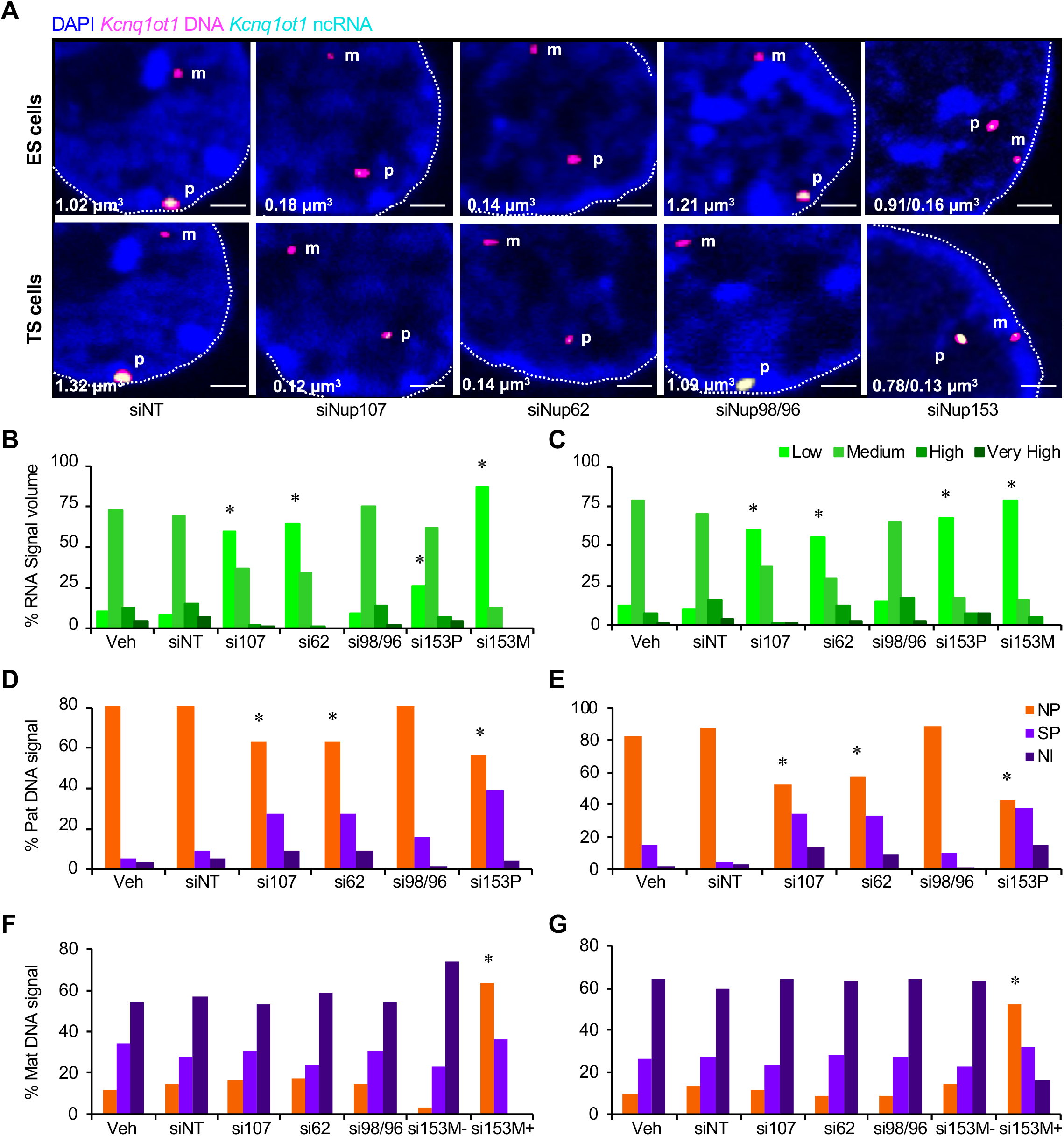
Nucleoporin depletion altered *Kcnq1ot1* ncRNA volume and positioning of the *Kcnq1ot1* domain at the nuclear periphery in ES and TS cells. (A) Representative confocal images of G1-synchronized control and *Nup*-depleted ES and TS nuclei; *Kcnq1ot1* DNA (red fluorescence converted to magenta); *Kcnq1ot1* ncRNA (green fluorescence converted to cyan); merge (white); DAPI staining (blue); n=4 biological samples; cell number=12G. RNA FISH signal intensity is measured in μm^3^; white dashed lines, nuclear rim. (B, C) Percent of cells with *Kcnq1ot1* ncRNA signal volume that is low (G-G.7 μm^3^), medium (G.7–1.4 μm^3^), high (1.4–2.1 μm^3^) or very high (>2.1 μm^3^) in ES and TS cells. In these cells, RNA signals were restricted to the paternal allele, except for *Nup153*-depleted cells where signal was detected from paternal (si153 P) and maternal (si153 M) alleles. (D-G) Distance of the paternal and maternal *Kcnq1ot1* domain from the nuclear periphery in control and *Nup*-depleted ES and TS cells. The maternal *Kcnq1ot1* domain was positioned randomly in nuclei. Cells lacking RNA FISH signal with detectable DNA FISH signals were included, while those without DNA signals were excluded. Nuclear periphery (NP), G-G.5 μm; subnuclear periphery (SP), G.5–1.5 μm; nuclear interior (NI), 1.5–4 μm; scale bar, 1 μm; *, significance p < G.G5 compared to vehicle control; error bars, s.e.m.; WT, wildtype; Veh, vehicle; siNT, non-targeting siRNA; si 1G7, *Nup107* siRNA; si62, *Nup62* siRNA; si98/96, *Nup9S/96* siRNA; si153, *Nup153* siRNA; si153 P, *Nup153*-depleted cells with a *Kcnq1ot1* ncRNA signal; si153 M−, *Nup153*− depleted cells lacking a maternal *Kcnq1ot1* ncRNA signal; si153 M+, *Nup153*-depleted cells possessing a *Kcnq1ot1* ncRNA signal from the maternal allele.

### NUP107, NUP62 and NUP153 regulate *Kcnq1ot1* domain positioning in ES and TS cells

We previously showed that the *Kcnq1ot1* ncRNA-coated domain was positioned at the nuclear periphery in XEN cells (Sachani et al., 2018). To test whether nuclear periphery positioning of the *Kcnq1ot1* ncRNA-coated domain was altered upon *Nup*-depletion in ES and TS cells, we calculated the distance of the *Kcnq1ot1* domain centroid from the nuclear periphery in G1-synchronized cells (Wu and Yao, 2013; Zullo et al., 2012). The maternal (0.1–0.8 μm^3^) and paternal (>0.8–1.3 μ.m^3^) *Kcnq1ot1* domains were clearly differentiated by domain volume using a centrally located DNA probe in ES and TS cells (Fig. S5A, B), similar to XEN cells (Sachani et al., 2018). In ES and TS controls, the paternal *Kcnq1ot1* domain was positioned at the nuclear periphery (0–0.5 μm from rim) in 86–92% and 83–88% of cells, at the sub-nuclear periphery (0.5–1.5 μm from rim) in 5–9% and 4–15% of cells, and at the nuclear interior (1.5–4 μm from rim) in 3–5% and 2–3% of cells, respectively. By comparison, *Nup107, Nup62* and *Nup153* depletion resulted in decreased positioning of the paternal *Kcnq1ot1* domain at the nuclear periphery in ES (63%, 63% and 57%, respectively) and TS (52%, 58%, and 43%, respectively) cells (Fig. 2D, E). In addition, the maternal *Kcnq1ot1* domain gained nuclear peripheral positioning in *Kcnq1ot1* ncRNA-positive, *Nup153*-*depleted* ES and TS cells (M+, 64% and 52%, respectively) compared to control, Nup98-depleted, and maternal *Kcnq1ot1* ncRNA-negative, *Nup153*-*depleted* ES and TS cells (control, 12–13% and 10–13%; *Nup98*, 15–17% and 9–12%; and M-Nup153, 3% and 14%, respectively), where domain positioning was random (Fig. 2F, G). Correlation between *Kcnq1ot1* ncRNA volume and *Kcnq1ot1* domain positioning showed that most ES and TS control cells with medium paternal *Kcnq1ot1* ncRNA volumes had domains positioned at the nuclear periphery (Figs. S6 and S7). In contrast, *Nup107*-, *Nup62*- and *Nup153*-*depleted* ES and TS cells possessed paternal domains with lower *Kcnq1ot1* ncRNA volumes that were shifted away from the nuclear periphery. Moreover, the maternal domain in *Nup153*-depleted cells gained low *Kcnq1ot1* ncRNA signal volume with corresponding nuclear periphery positioning. *Nup98/96*-depleted cells displayed similar volumes and positioning as control cells. These results indicate that NUP107, NUP62, and NUP153 targeted the *Kcnq1ot1* domain to the nuclear periphery in ES and TS cells, similar to XEN cells (Sachani et al., 2018).

### NUP107, NUP62 and NUP153 bind to the *Kcnq1ot1* ICR in ES and TS cells

To investigate nucleoporin interactions at the *Kcnq1ot1* ICR in ES and TS cells, quantitative chromatin immunoprecipitation (ChIP) was performed using antibodies for mAb414 [ChIP-grade NUP107 and NUP62 antibodies, not available; interacted primarily with NUP62, NUP107 and NUP160 in ES and TS cells (Fig. S2)], NUP98 and NUP153 across 21 sites at the *Kcnq1ot1* ICR and H3K4me1 enhancer element region (Figs. S8A, 3A). In control ES and TS cells, mAb414 (NUP107/62) and NUP153 were significantly enriched at the *Kcnq1ot1* ICR (IC3 and IC4), which flanked the *Kcnq1ot1* ncRNA promoter (Figs. S8B, S8C, 3B, and 3D). mAb414 was also enriched at the enhancer element region (E1 and E2) in ES and TS cells (S8B, S8C, 3B, and 3D), while no enrichment was observed for NUP98 (Figs. S8B, S8C). Prior to parental-specific investigation, we verified equal allelic detection for input chromatin at the *Kcnq1ot1* ICR binding sites and a control site (Fig. S9). In ES and TS control cells, NUP107/NUP62 and NUP153 primarily occupied the paternal allele at these sites (Fig. 3C, 3E). This enrichment was significantly reduced at the paternal *Kcnq1ot1* ICR and paternal enhancer element sites in *Nup107/Nup62* double-depleted cells compared to the control (Fig. 3C), as well as at the paternal *Kcnq1ot1* ICR sites upon *Nup153* depletion (Fig. 3E). Overall, NUP107/NUP62 and NUP153 interactions were conserved at the paternal *Kcnq1ot1* ICR and paternal enhancer element sites in ES, TS and XEN cells, although in XEN cells NUP153 occupancy was also found at the maternal *Kcnq1ot1* ICR (Sachani et al., 2018).

**Figure 3:**
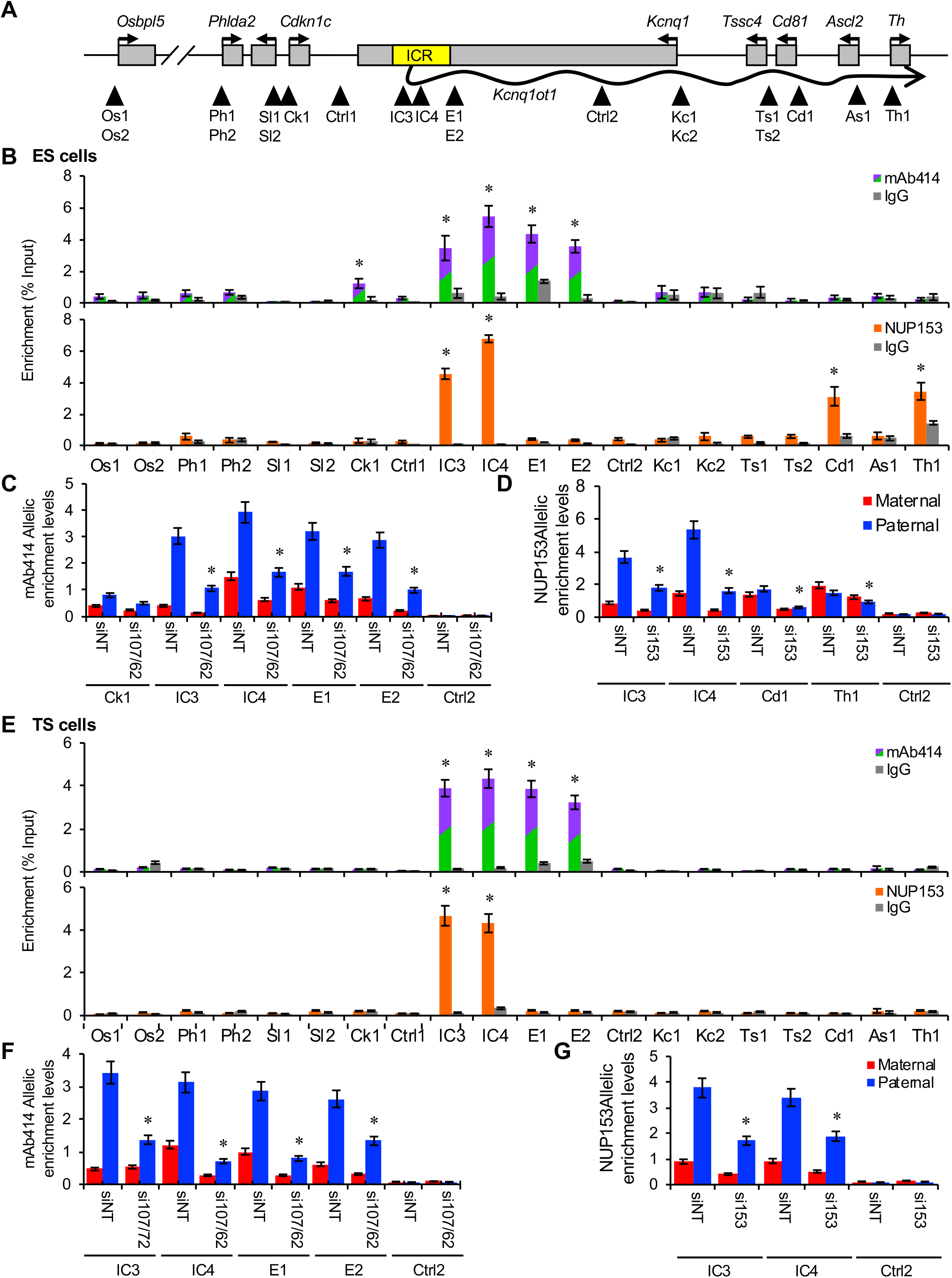
NUP107/62 and NUP15S interaction with the *Kcnq1ot1* domain in control ES and TS cells. (A) The *Kcnq1ot1* domain with regions of analysis (arrowheads); Os1, Os2, *Osbpl5* promoter; Ph1, Ph2, *Phlda2* promoter; Sl1, Sl2, *Slc22alS* promoter; Ck1, *Cdkn1c* promoter; IC3, IC4, *Kcnq1ot1* ICR; E1, E2, enhancer element; Kc1, Kc2, *Kcnq1* promoter; Ts1, Ts2, *Tssc4* promoter; Cd1, *CdS1* promoter; As1, *Ascl2* promoter; Th1, *Th* promoter; Ctrl1, Ctrl2: control negative sites. (B) Quantitative ChIP analysis at regions across the domain using mAb414 (top) and NUP153 (bottom) antibodies in wild type ES (n=3 biological samples with 4 technical replicates per sample). (C, D) Total enrichment and quantitative allelic analysis using mAb414 (C) and NUP153 (D) antibodies in siNT and *Nup153*-*depleted* ES cells. (E) Quantitative ChIP analysis at regions across the domain using mAb414 (top) and NUP153 (bottom) antibodies in wild type TS cells (n=3 biological samples with 4 technical replicates per sample). (F, G) Total enrichment and quantitative allelic analysis using mAb414 (F) and NUP153 (G) antibodies in siNT and *Nup153*-depleted;TS cells. Allelic proportions are shown as percent of total enrichment levels (n=3 biological samples with 3 technical replicates per replicate). Error bars, s.e.m.; * indicates significance p < 0.05 compared to IgG or siNT control.

### NUP107, NUP62 and NUP153 bind to select imprinted gene promoters in ES and TS cells

To investigate whether nucleoporin interactions occurred at sites outside the ICR within the *Kcnq1ot1* imprinted domain in ES and TS cells, ChIP was performed using antibodies for mAb414 and NUP153 at promoters of imprinted genes in the domain (Fig. 3A). Prior to investigation, equal allelic detection of input chromatin was first verified at imprinted gene promoters in the *Kcnq1ot1* imprinted domain (Fig. S10). In control ES cells, we observed significant NUP107/NUP62 (mAb414) enrichment at the *Cdkn1c* promoter (Ck1), and NUP153 enrichment at the *Cd81* (Cd1) and *Th* (Th1) promoters (Fig. 3B). While paternal NUP107/NUP62 enrichment at the *Cdkn1c* promoter was not significantly altered in *Nup107/Nup62*-*depleted* ES cells, NUP153 enrichment was significantly reduced at the maternal and paternal *Cd81* and *Th* promoters in *Nup153*-depleted ES cells, compared to control cells (Fig. 3C). By comparison in TS cells, no further sites of enrichment were observed (Fig. 3D). The results diverged from those of XEN cells, where NUP107/62 occupancy occurred primarily at the paternal *Osbpl5* promoter, while NUP153 binding resided at the paternal *Kcnq1* and *Cd81* promoters (Sachani et al., 2018).

### Nucleoporins regulate paternal allele silencing in ES and TS cells

We next investigated the effects of *Nup*-depletion on paternal allelic silencing in ES and TS cells using droplet digital PCR technology to determine absolute mRNA abundance from each parental allele using FAM and HEX strain-specific probes. Compared to control ES cells, we observed a significant reactivation of the paternal *Cdkn1c* allele upon *Nup107, Nup62* and *Nup153* depletion by 351–710 copies (Figs. 4A and S11). *Nup153* depletion also resulted in significant reactivation of the paternal *Cd81* allele by 1,811,806 copies compared to control ES cells. *Nup98/96* depletion produce a significant increase in paternal *Kcnq1* allele by 42 copies compared to controls. By comparison in TS cells, *Nup107, Nup62* and *Nup153* depletion resulted in reactivation of the paternal *Slc22a18, Cdkn1c* and paternal *Kcnq1* allele by 164–341, 12,778–18,687, and 59–64 copies, respectively, compared to controls (Figs. 4B and S12). *Nup153* depletion also reactivated the paternal *Osbpl5* by 532 copies (Figs. 4B and S12). On the maternal allele in *Nup153*-depleted ES cells, reactivation of the maternal *Kcnq1ot1* ncRNA (Fig. 1B) corresponded to silencing of normally-expressed maternal alleles of several upstream protein-coding genes, including *Osbpl5, Phlda2, Slc22a18* and *Kcnq1* with reduced maternal copies of 44, 2418, 653 and 62, respectively, as well as at the maternal *Osbpl5* allele by 536 copies in TS cells, compared to controls (Figs. 4B and S12). Overall, our data indicate that reactivation of paternal alleles was not domain-wide, similar to results observed in XEN cells (Sachani et al., 2018). Furthermore, divergent subset of genes were paternally reactivated upon *Nup*-depletion in ES, TS as well as XEN cells (Sachani et al., 2018). In addition, silencing of specific maternal expressed alleles occurred in *Nup153*-depleted ES and TS cells, but not in XEN cells (Sachani et al., 2018).

**Figure 4:**
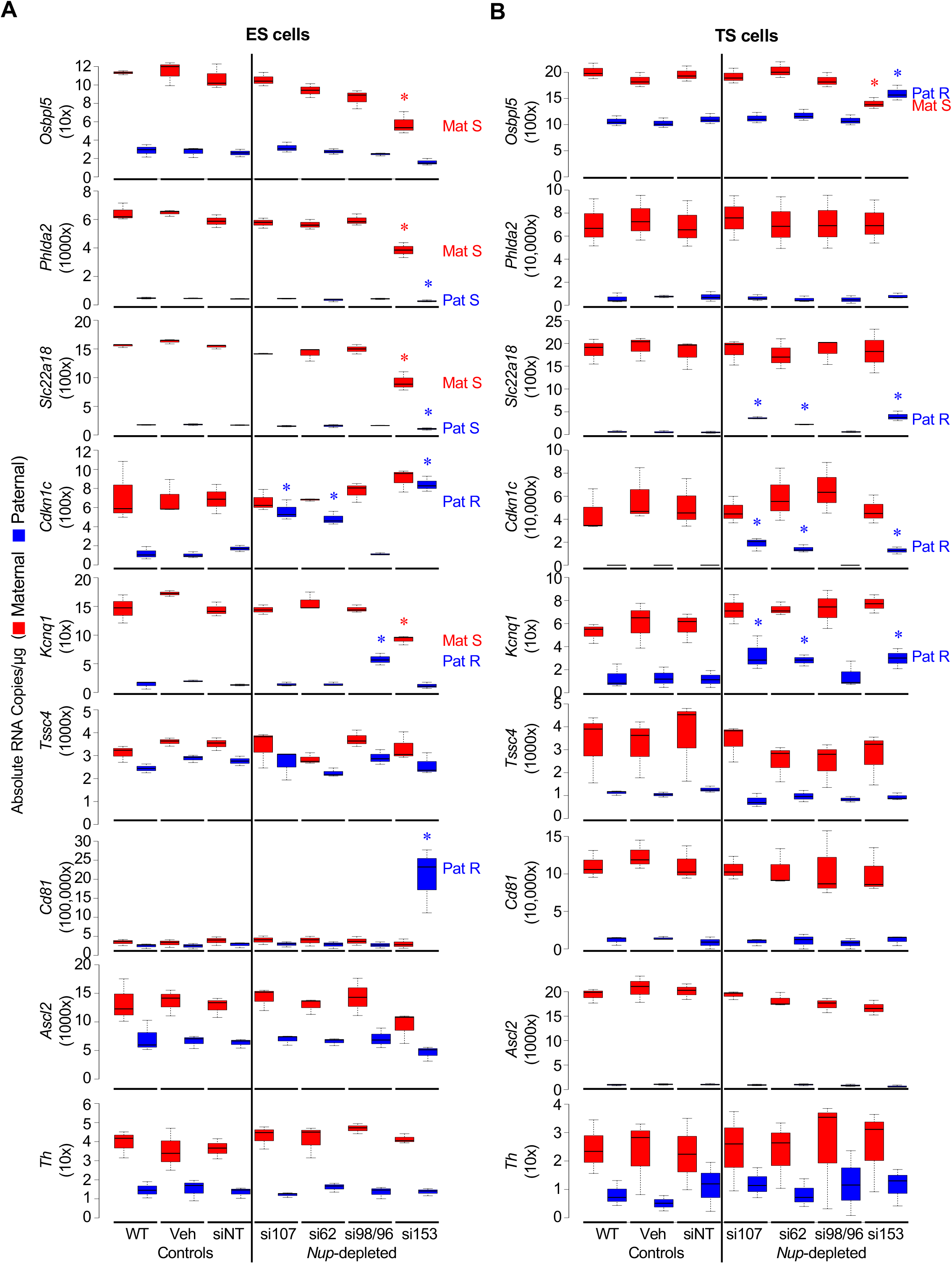
Nucleoporin depletion reactivated subsets of paternal alleles of imprinted genes within the *Kcnq1ot1* domain in ES and TS cells. Absolute allelic transcript abundance was determined in control and *Nup*-depleted (A) ES and (B) TS cells (n=3 biological samples). Center lines, medians; whiskers, 1.5 times the interquartile range from 25th and 75th percentiles; box limits, 25th and 75th percentiles as determined by R software; blue bars, paternal allele; red bars, maternal allele, Mat R, reactivated maternal; Mat S, silenced maternal; Pat R, reactivated paternal; Pat S, silenced allele; *, significance p < 0.01 compared to the siNT control; error bars, s.e.m.; Veh, vehicle; siNT, non-targeting siRNA; si107, *Nup107* siRNA; si62, *Nup62* siRNA; si98/96, *Nup9S/96* siRNA; si153, *Nup153* siRNA.

### Nucleoporin depletion did not impair nuclear-cytoplasmic transport

Since nucleoporins control nuclear-cytoplasmic import and export, alterations in *Kcnq1ot1* imprinted domain regulation upon nucleoporin depletion could be explained by impaired nuclear-cytoplasmic transport. Firstly, impaired transport function could result from altered levels in nucleoporins within the nuclear pore complex upon depletion of one component. Examination of nucleoporins from different structural components of the nuclear pore complex did not generate any change in nucleoporin levels in *Nup*-depleted ES and TS cells compared to controls (Fig. S13A, S13B). This suggests that the assembly of the nuclear pore complex was not affected upon *Nup*-depletion. Furthermore, investigation of exogenous and endogenous NLS containing protein nuclear import (Fig S14A, S14B), or accumulation of polyA-mRNA within nuclei reflective of aberrant nuclear export (Fig. S14C) revealed no change in nuclear-cytoplasmic transport. Finally, cell proliferation was also unaltered, indicating normal transport function (Fig. S15). These results were similar to those observed in XEN cells (Sachani et al., 2018).

### NUP107, NUP62 and NUP153 depletion did not alter imprinted DNA methylation at the *Kcnq1ot1* ICR

Another explanation for *Kcnq1ot1* imprinted domain regulation was a change in DNA methylation status at the *Kcnq1ot1* ICR. For example, alteration at the paternal *Kcnq1ot1* domain in *Nup107, Nup62* and *Nup153*-*depleted* cells could be due to a gain in DNA methylation at the paternal *Kcnq1ot1* ICR, whereas changes in maternal *Kcnq1ot1* imprinted domain regulation upon *Nup153* depletion may be caused by loss of DNA methylation at the maternal *Kcnq1ot1* ICR. However, we observed that the DNA methylation status of the *Kcnq1ot1* ICR was unaltered in *Nup*-depleted ES and TS cells (Fig. S16) compared to controls, which was similar to observations reported for XEN cells (Sachani et al., 2018).

### NUP107, NUP62 and NUP153 depletion alter chromatin state at the *Kcnq1ot1* ICR

While DNA methylation was unchanged, alterations in chromatin modifications at the *Kcnq1ot1* ICR could contribute to altered *Kcnq1ot1* imprinted domain regulation. ChIP assays were performed on control and *Nup*-depleted ES and TS cells using histone 3 lysine 4 trimethylation (H3K4me3) as well as RNA Polymerase II (RNAPII) antibodies as marks for active transcription, and H3K9me2 and H3K27me3 antibodies as marks for repressed chromatin. In addition, ChIP was performed with antibodies for histone methyltransferases KMT2A, EHMT and EZH2, which confer these histone modifications. First, experimental controls were performed. Active and repressive modifications were validated at the ES cell-expressed *Pou5f1*, and the TS cell-expressed *Cdx2* gene promoters (Lim *et. al*. 2008) (Fig. S17); and total KMT2A, EHMT and EZH2 protein levels were verified as unchanged between control and *Nup*-depleted ES and TS cells (Fig. S18A, S18B). Next, quantitative-allelic ChIP was performed at the *Kcnq1ot1* ICR to assess active and repressive modification, as well as modifier enrichment. Compared to the paternal *Kcnq1ot1* ICR in control ES and TS cells, *Nup107*-*, Nup62*-, and *Nup153*-*depleted* cells had significantly decreased paternal H3K4me3 and KMT2A (Fig. 5A, 5B) as well as RNAPII (Fig. S19A, S19B) enrichment. While there was an increase in paternal allelic EHMT and EZH2 occupancy in *Nup107*- and *Nup62*-*depleted* ES and TS cells, there was limited change in H3K9me2 or H3K27me3 enrichment at the paternal *Kcnq1ot1* ICR (Figs. 5A, 5B). In *Nup153*-*depleted* ES and TS cells, the maternal *Kcnq1ot1* ICR had significantly increased H3K4me3, KMT2A and RNAPII enrichment, compared to control cells (Figs. 5A, 5B and S19A, 19B). This was concomitant with significantly decreased H3K9me2 and H3K27me3 enrichment, in conjunction with reduced EHMT and EZH2 occupancy at the maternal *Kcnq1ot1* ICR. No changes in histone modification was observed at the *Kcnq1ot1* ICR upon *Nup98/96* depletion. These results were similar to those observed in XEN cells (Sachani et al., 2018)

**Figure 5:**
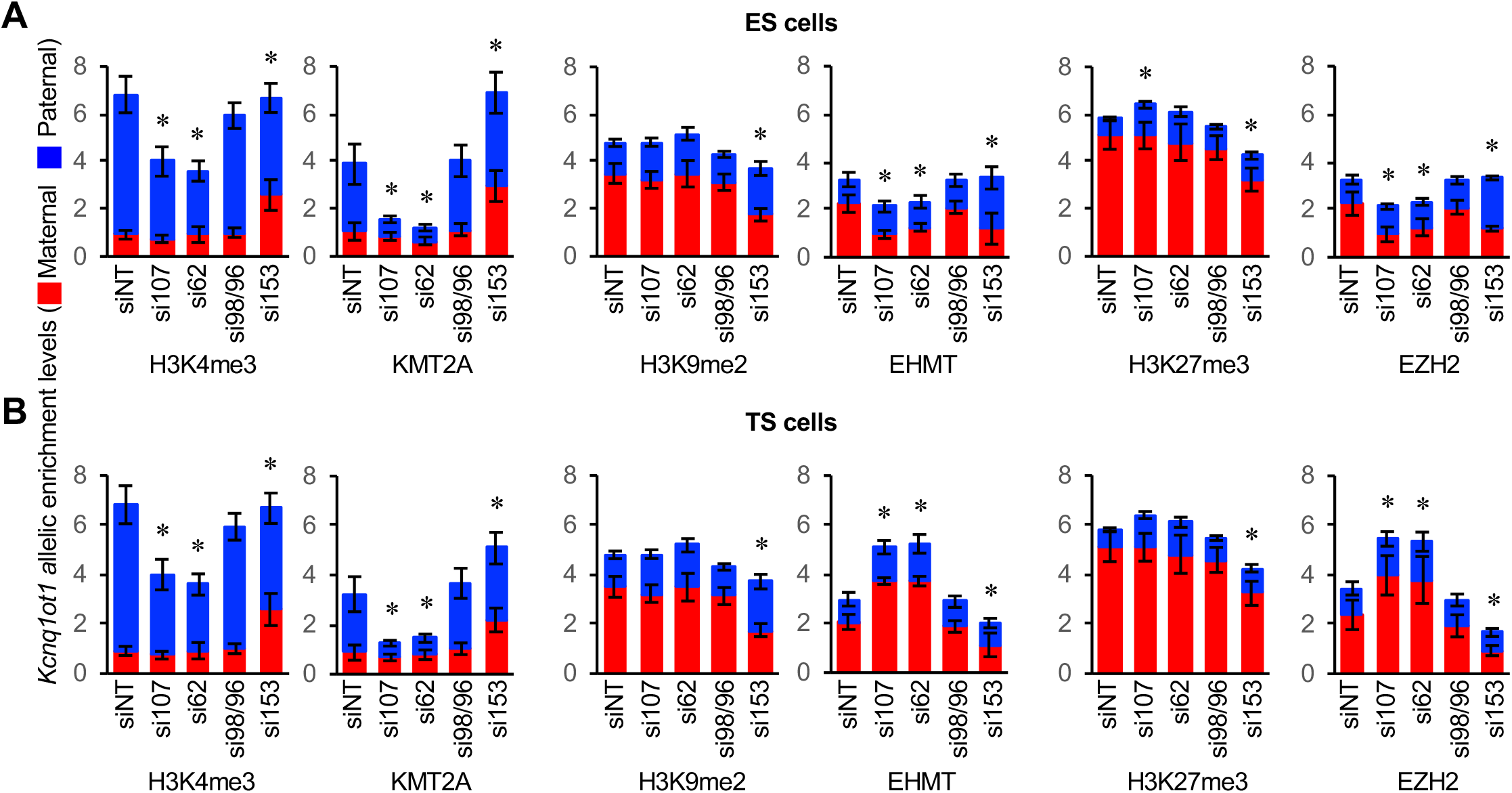
Nucleoporin depletion disrupted histone modifications and histone methyltransferase enrichment at the *Kcnq1ot1* ICR in ES and TS cells. H3K4me3, KMT2A, H3K9me2, EHMT, H3K27me3 and EZH2 ChIP at the paternal and maternal *Kcnq1ot1* ICR (IC3 site) in control and *Nup*-depleted (A) ES and (B) TS cells (n=3 biological samples with 3 technical replicates per sample). Allelic proportions are represented as a percent of the total enrichment level. *, significance p < 0.05 compared to siNT control; error bars, s.e.m.; WT, wildtype; Veh, vehicle; siNT, non-targeting siRNA; si107, *Nup107* siRNA; si62, *Nup62* siRNA; si98/96, *Nup98/96* siRNA, si153, *Nup153* siRNA.

### Nucleoporins maintain chromatin state at imprinted gene promoters

We extended quantitative-allelic ChIP assays for histone modifications, their respective histone methyltransferases, as well as RNAPII, to promoter regions of imprinted genes within the *Kcnq1ot1* domain. This was to address whether altered expression of imprinted genes in the *Kcnq1ot1* domain was coincident with changes in chromatin state upon nucleoporin depletion. In ES cells, compared to controls, the paternal *Cdkn1c* promoter demonstrated the broadest changes to chromatin state, with a significant increase in H3K4me3 and KMT2A enrichment (Fig. 6), as well as increased RNAPII occupancy (Fig. S20A), upon *Nup107, Nup62* and *Nup153* depletion. A corresponding decrease was observed in paternal H3K9me2, EHMT, H3K27me3 and EZH2 enrichment. Similar chromatin changes were observed at the paternal *Cd81* promoter upon *Nup153* depletion and at the paternal *Kcnq1* promoter upon *Nup98/96* depletion (Fig. 6 and S20A). In TS cells, compared to controls, we found significantly increased H3K4me3, KMT2A and RNAPII enrichment (Fig. 7 and S20B), along with significantly decreased H3K9me2, EHMT, H3K27me3 and EZH2 at the paternal *Slc22a18, Cdkn1c* and *Kcnq1* promoters upon *Nup107, Nup62* and *Nup153* depletion, as well as at the *Osbpl5* promoter upon *Nup153* depletion (Fig. 7). Thus, for both ES and TS cells, changes in chromatin state upon nucleoporin depletion were consistent with the paternal allelic reactivation that was observed for these genes (Fig. 4), indicating that nucleoporins regulate histone modifications at selected imprinted gene promoters in ES and TS cells, similar to XEN cells (Sachani et al., 2018).

**Figure 6:**
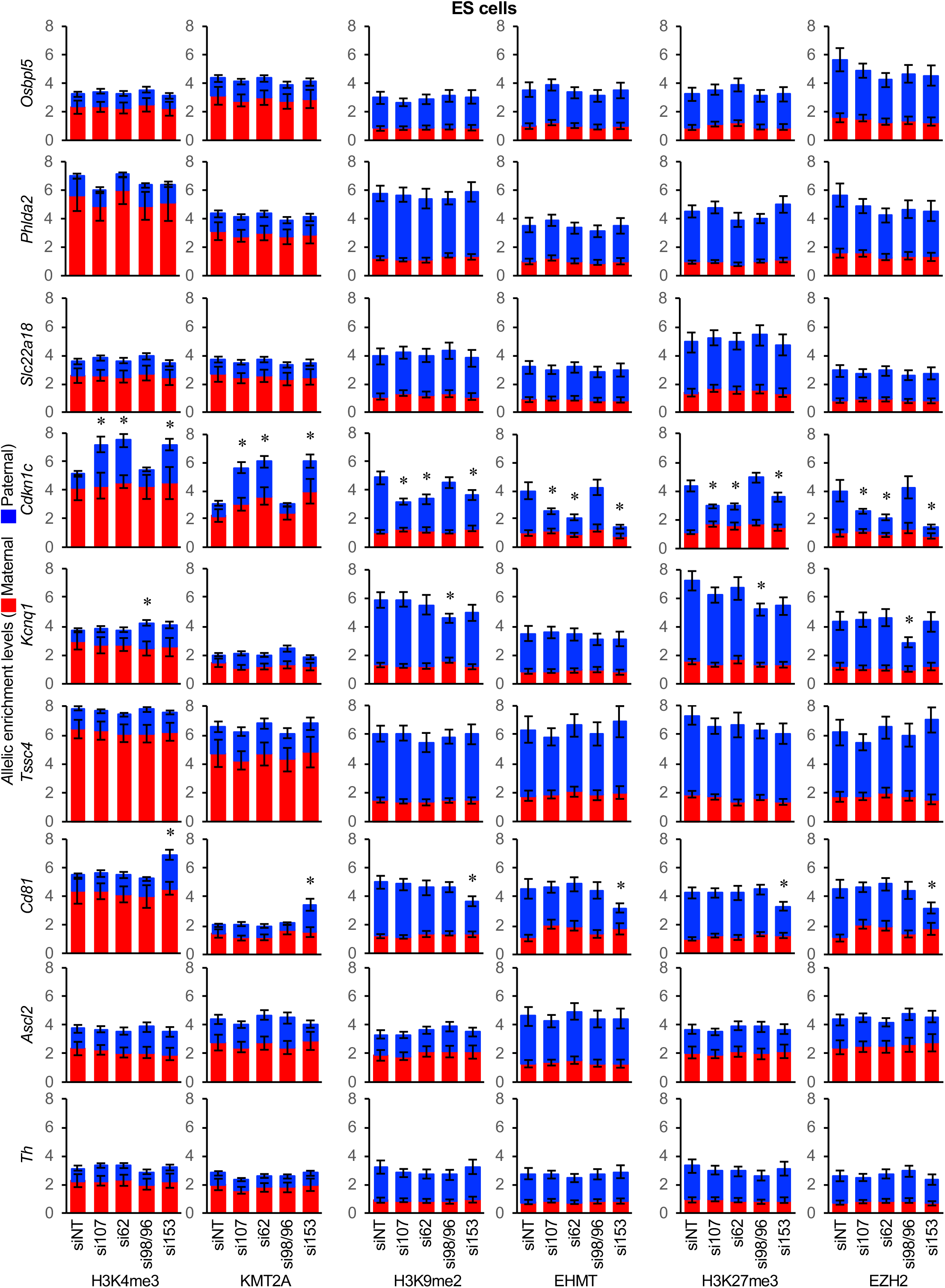
Nucleoporin depletion disrupted histone modifications and histone methyltransferase enrichment at reactivated imprinted gene promoters in ES cells. H3K4me3, KMT2A, H3K9me2, EHMT, H3K27me3 and EZH2 ChIP at the paternal and maternal imprinted gene promoters in control and *Nup*-depleted ES cells (n=3 biological samples with 3 technical replicates per sample). Allelic proportions are shown as a percent of total enrichment level. *, significance p < 0.05 compared to siNT control; error bars, s.e.m.; WT, wildtype; Veh, vehicle; siNT, non-targeting siRNA; si107, *Nup107* siRNA; si62, *Nup62* siRNA; si98/96, *Nup98/96* siRNA, si153, *Nup153* siRNA.

**Figure 7:**
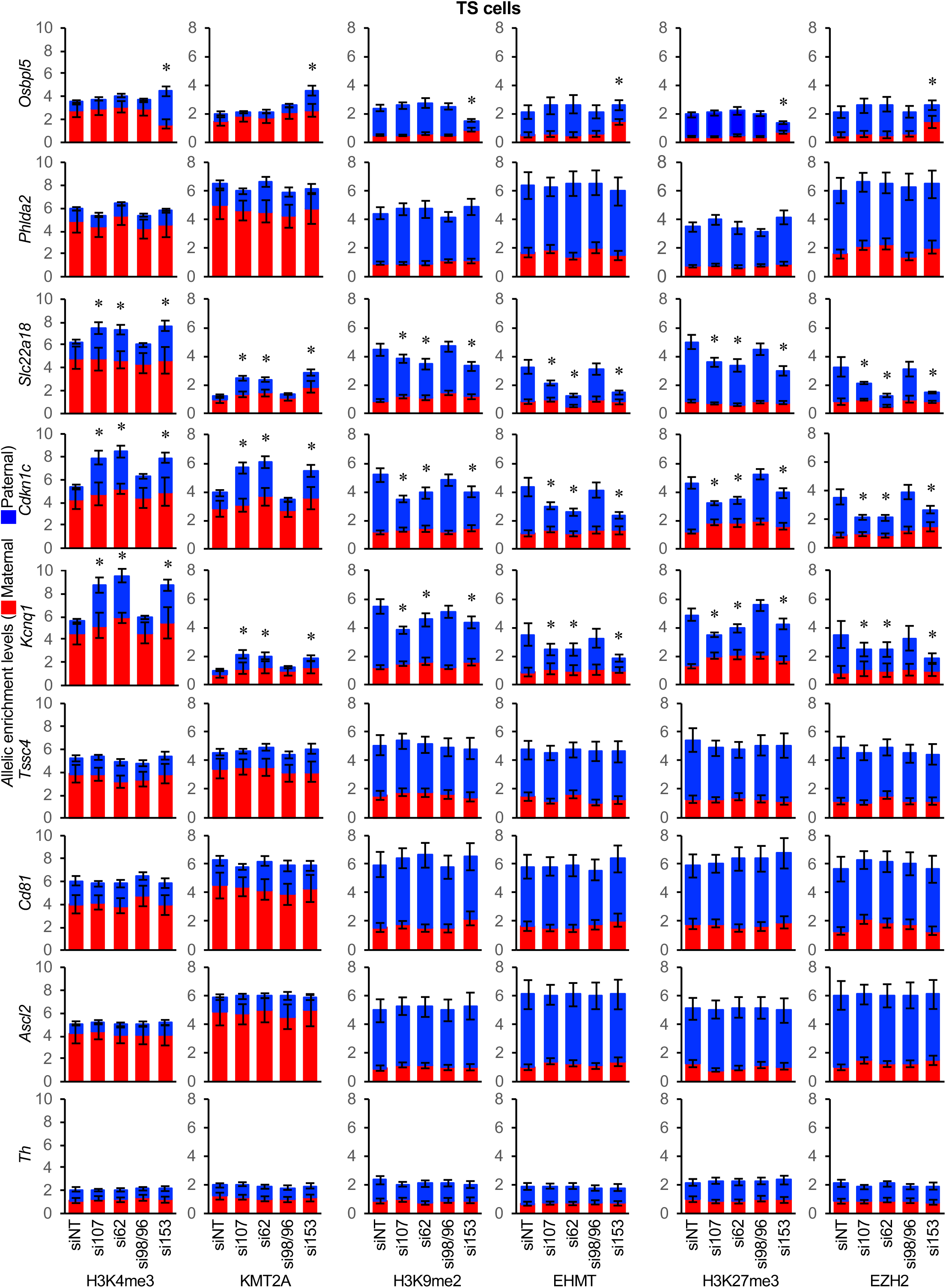
Nucleoporin depletion perturbed histone modifications and modifier enrichment at reactivated imprinted gene promoters in TS cells. H3K4me3, KMT2A, H3K9me2, EHMT, H3K27me3 and EZH2 ChIP at the paternal and maternal imprinted gene promoters in control and *Nup*-depleted TS cells (n=3 biological samples with 3 technical replicates per sample). Allelic proportions are shown as a percent of total enrichment level. *, significance p < 0.05 compared to siNT control; error bars, s.e.m.; WT, wildtype; Veh, vehicle; siNT, non-targeting siRNA; si107, *Nup107* siRNA; si62, *Nup62* siRNA; si98/96, *Nup98/96* siRNA, si153, *Nup153* siRNA.

### NUP107, NUP62 and NUP153 promote CTCF and cohesin complex interactions at the paternal *Kcnq1ot1* ICR in ES cells

In addition to histone modification, changes in CTCF and/or cohesin complex interactions at the *Kcnq1ot1* imprinted domain could contribute to altered regulation of this domain upon *Nup*-*depletion*. To determine whether CTCF and/or cohesin proteins interact with the *Kcnq1ot1* ICR, ChIP was performed using CTCF, SMC1A and SMC3 antibodies in ES and TS cells. As a control for the antibodies, we validated CTCF, SMC1A and SMC3 binding at the maternal *H19* and the paternal *Peg3* ICRs, and an absence of binding at the *H19* exon 5 and *Peg3* exon 2 negative control sites (Verona et al., 2008) in ES cells (Fig. S21). In TS cells, CTCF, SMC1A and SMC3 were enriched preferentially at the paternal *Peg3* ICR, but not at the *H19* ICR. We also verified that *Nup*-depletion did not alter CTCF, SMC1A and SMC3 total protein levels compared to controls (Fig. S22). Next, CTCF interactions were investigated at the *Kcnq1ot1* ICR and enhancer element sites (Fig. S23A). Similar to previously published data (Fitzpatrick et al., 2007; Hark et al., 2000), CTCF occupied the two CTCF-binding sites (IC3, IC4) within the *Kcnq1ot1* ICR in ES cells (Figure S23B). SMC1A and SMC3 enrichment was also observed at the CTCF-binding sites (IC3, IC4) in ES cells (Fig. S23B). Surprisingly, no enrichment of CTCF, SMC1A or SMC3 was found at the *Kcnq1ot1* ICR sites in TS cells (Fig. S23C). Thus, the disruptive effects of *Nup* depletion on CTCF and cohesin complex binding to the *Kcnq1ot1* ICR were only determined for ES cells. Compared to ES cell controls, CTCF, SMC1A and/or SMC3 enrichment was significantly reduced at the paternal *Kcnq1ot1* ICR sites (IC3, IC4) upon *Nup107, Nup62* and/or *Nup153* depletion (Fig. 8). These results indicate that NUP107, NUP62 and/or NUP153 play a role in CTCF and cohesin complex interaction on the paternal *Kcnq1ot1* ICR in ES cells. Overall, these results diverge from XEN cells where only cohesin complex proteins, SMC1A and SMC3, but not CTCF occupied the paternal *Kcnq1ot1* ICR (Sachani et al., 2018).

**Figure 8:**
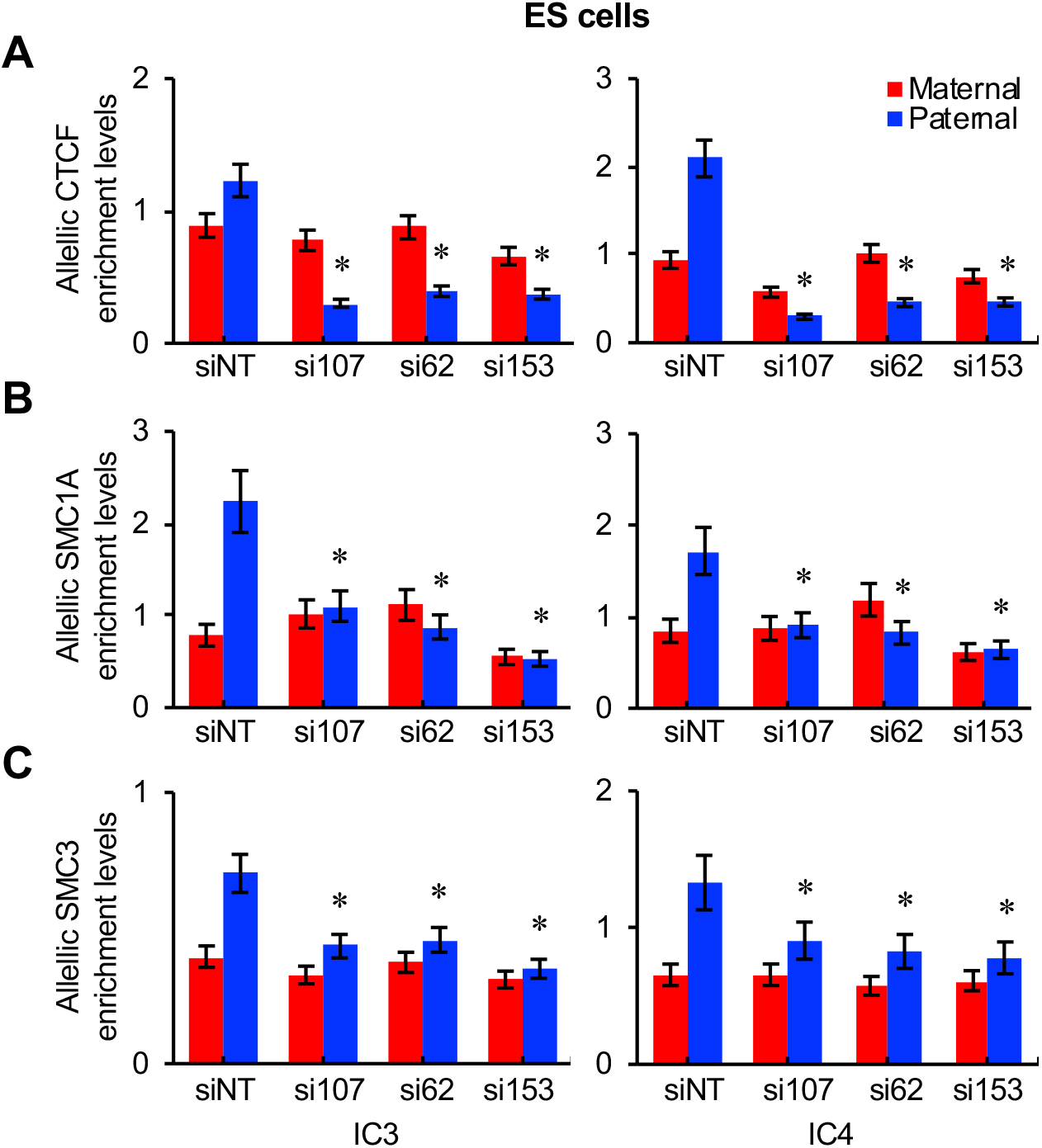
Reduction in CTCF, SMC1A and SMC3 occupancy at the paternal *Kcnq1ot1* ICR upon nucleoporin depletion in ES cells. Quantitative ChIP analysis for (A) CTCF, (B) SMC1A and (C) SMC3 at positive sites of mAb414 and NUP153 enrichment in control and *Nup107, Nup62* and Nup153-depleted ES cells (n=3 biological samples with 3 technical replicates per sample). Allelic proportions are shown as a percent of total enrichment level. Error bars, s.e.m.; *, significance p < 0.05 compared to siNT control.

## DISCUSSION

### Convergent nucleoporin-mediated mechanism with divergent regulation of *Kcnq1ot1* imprinted domain

In a recent study, we discovered a nucleoporin-mediated mechanism that governs *Kcnq1ot1* imprinted domain regulation in XEN cells (Sachani et al., 2018). Here, we investigated whether this mechanism was conserved in ES and TS cells. Our findings demonstrated that many features of NUP107, NUP62 and NUP153 regulation at the *Kcnq1ot1* imprinted domain in XEN cells were conserved in ES and TS cells. Firstly, NUP107/NUP62 and NUP153 bound the paternal *Kcnq1ot1* ICR, while NUP107/NUP62 also interacted with the paternal enhancer element downstream of the ICR in all three lineages, potentially as off-pore function (Fig. S24). Secondly, NUP107, NUP62 and NUP153 depletion reduced paternal *Kcnq1ot1* ncRNA expression and its volume at the paternal *Kcnq1ot1* domain, and shifted the paternal *Kcnq1ot1* domain away from the nuclear periphery in all three cell lineages (Sachani et al., 2018). Thirdly, the maternal *Kcnq1ot1* ncRNA was reactivated, and the maternal *Kcnq1ot1* domain gained nuclear periphery positioning in *Nup153*-*depleted* ES, TS cells as well as XEN cells (Sachani et al., 2018). Finally, in all three cell lineages, DNA methylation of the *Kcnq1ot1* ICR was unaltered. Instead, changes in *Kcnq1ot1* ncRNA expression were concomitant with altered histone modifications and histone methyltransferase occupancy (Sachani et al., 2018). Notably, for all three cell lineages, no change in nuclear-cytoplasmic transport was observed.

Divergence in NUP107, NUP62 and NUP153 regulation at the *Kcnq1ot1* imprinted domain between the three cell lineages also emerged from our findings. Firstly, while NUP107/NUP62 and NUP153 bound the *Kcnq1ot1* ICR, other binding sites within the paternal *Kcnq1ot1* imprinted domain were gene-and lineage-specific. In XEN cells, NUP107/NUP62 occupied the paternal *Osbpl5* promoter, while NUP153 bound to the paternal *Kcnq1* and *CdS1* promoters (Fig. S24). In ES cells, NUP107/NUP62 occupancy was observed at the paternal *Cdkn1c* promoter, whereas NUP153 bound to the maternal and paternal *CdS1* and *Th* promoters. In TS cells, NUP107/NUP62 and NUP153 were not enriched at any additional imprinted gene promoters. Nucleoporin binding at these promoters is likely in an off-pore capacity (Sachani et al., 2G18). Secondly, with respect to expression of imprinted protein-coding genes in the *Kcnq1ot1* imprinted domain, divergent sets of paternal alleles were reactivated in ES, TS as well as XEN cells upon *Nup*-depletion. In XEN cells, *Nup107, Nup62* and *Nup153* depletion led to paternal reactivation of the core group of genes, *Slc22a18, Cdkn1c*, and *Kcnq1*. Furthermore, gene-specific reactivation was observed for the paternal *Osbpl5* and *Phlda2* alleles upon *Nup107* depletion, the paternal *Osbpl5* allele upon *Nup62* depletion, and the paternal *CdS1* allele upon *Nup153* depletion (Sachani et al., 2G18). In ES cells, the paternal *Cdkn1c* was reactivated upon *Nup107* and *Nup62* depletion, while the paternal *Cdkn1c* and *CdS1* alleles were reactivated upon *Nup153* depletion. In TS cells, a core group of paternal alleles for *Slc22alS, Cdkn1c*, and *Kcnq1* were reactivated upon *Nup107, Nup62* and *Nup153* depletion, while paternal *Osbpl5* reactivation also occurred in *Nup153*-depleted TS cells. Thirdly, with respect to the maternal *Kcnq1ot1* imprinted domain, while the maternal *Kcnq1ot1* ncRNA allele was reactivated in *Nup153*- depleted XEN, ES and TS cells, alterations in the expression of maternal allele of imprinted protein-coding genes was divergent. While no changes in maternal expression of imprinted protein-coding genes occurred in XEN cells (Sachani et al., 2G18), there was a decrease in the maternally expressed *Osbpl5, Phlda2, Slc22alS* and *Kcnq1* alleles in ES cells as well as the maternal *Osbpl5* allele in TS cells, compared to controls. Finally, investigation of CTCF and cohesin complex occupancy at the nucleoporin binding sites within the paternal *Kcnq1ot1* ICR revealed that only cohesin bound these sites in XEN cells (Sachani et al., 2G18), while both CTCF and cohesin were enriched at these sites in ES cells (Fig. S24). In contrast, CTCF and cohesin occupancy was absent at the nucleoporin binding sites within *Kcnq1ot1* ICR in TS cells. All together, these results indicate a convergent nucleoporin-mediated mechanism with divergent regulation of *Kcnq1ot1* imprinted domain in ES, TS and XEN stem cell lineages of the early preimplantation embryo.

### Role of NUP107, NUP62 and NUP153 in *Kcnq1ot1* imprinted domain structure

A comparison of nucleoporin-mediated regulation of the *Kcnq1ot1* imprinted domain in the three cell lineages points to the complexity of maintaining imprinting across a large, one-megabase, imprinted domain during early development. To gain a further understanding of the *Kcnq1ot1* imprinted domain, we mapped chromatin topologically associated domain boundaries from ES cell Hi-C data for the *Kcnq1ot1* domain (Dixon et al., 2012; MacDonald et al., 2015). The *Kcnq1ot1* domain mapped into three TADs in ES cells (Fig. S24). TAD1 harbors the *Osbpl5* gene plus 5 non-imprinted genes, TAD2 covers a region from *Phlda2* to the *Kcnq1*, while the TAD3 extends from *Trpm5* to *Th*. Based on the stability of TADs through development (Hug et al., 2017; Pope et al., 2014), we present the following models for NUP107, NUP62 and NUP153-mediated regulation of the paternal *Kcnq1ot1* imprinted domain in XEN, ES and TS cells. In TAD2 of XEN cells, nuclear pore-situated NUP107, NUP62 and NUP153 interact with cohesin and KMT2A at the *Kcnq1ot1* ncRNA promoter, generating boundaries that demarcate the active paternal *Kcnq1ot1* ICR from repressive chromatin at the paternal *Slc22a18* and *Cdkn1c* alleles on one flank and the paternal *Kcnq1* allele on the other flank, including through nucleoporin-dependent EHMT and EZH2 occupancy (Fig. S24; Sachani et al., 2018). The paternal *Phlda2* allele likely has additional repressive mechanisms. NUP107/NUP62 occupancy at the paternal enhancer element may also generate an additional insulator/boundary that isolates the enhancer element from the repressed paternal *Kcnq1* allele. Consistent with this, a previous study reported that a paternal 200 kb-intrachromosomal loop between the enhancer element and *Kcnq1* promoter in fibroblast cells was involved in maintaining paternal *Kcnq1* silencing (Korostowski et al., 2011; Zhang et al., 2014). In TAD1, NUP107/NUP62 binding with EHMT and EZH2 at the paternal *Osbpl5* promoter, and in TAD3, NUP153 binding together with EHMT and EZH2 at the paternal *Cd81* promoter could insulate repressive chromatin at *Osbpl5* and *Cd81* from active chromatin at neighbouring non-imprinted genes. In ES cells, cohesin and CTCF bind the same nucleoporin binding sites in the paternal *Kcnq1ot1* ICR of TAD2, likely producing the same chromatin boundaries that promote *Kcnq1ot1* ncRNA transcription (Fig. S24). NUP107/NUP62 occupancy at the paternal enhancer element may isolate the enhancer element from the repressed paternal *Kcnq1* allele. Furthermore, in ES cells, NUP107/NUP62 enrichment together with EHMT and EZH2 at the paternal *Cdkn1c* promoter region likely separates paternal *Cdkn1c* silencing from the other upstream paternal alleles. In support of this, mapping of CTCF- and cohesin-dependent HiChIP data from ES cells revealed a chromatin interaction from a region between the *Slc22a18* and *Cdkn1c* genes to a region between *Kcnq1* and *Trpm5* (Mumbach et al., 2016), potentially insulating *Cdkn1c* from *Slc22a18* and *Phlda2*. If this is the case, additional repressive mechanisms would be required for paternal *Slc22a18* and *Phlda2* silencing. In TAD3, NUP153 occupancy at the paternal *Cd81* and *Th* promoters may mark repressive chromatin from active chromatin. In TS cells, neither CTCF nor the cohesin complex assembled at the paternal *Kcnq1ot1* ICR, suggesting that other proteins, such as YY1, mediate chromatin interactions within TAD2 (Fig. S24). In TAD3, additional repressive mechanisms are likely required for the paternal *Cd81* and *Th* silencing. To test these models, further investigation is required to determine higher-order, chromatin structure at the *Kcnq1ot1* imprinted domain in ES, TS and XEN cells.

### Tissue-specific regulation between embryonic and extraembryonic lineages

Previous studies of mid-gestation conceptuses have identified differential regulation of genes in the *Kcnq1ot1* imprinted domain. The inner core genes *(Slc22a18, Cdkn1c* and *Kcnq1)* which exhibit imprinted expression in embryonic and extraembryonic tissues were classified as ubiquitously imprinted, whereas the outer genes, which possessed imprinted expression in the placenta but not the embryo, were classified as placenta-specific imprinted genes (Lewis et al., 2004; Lewis et al., 2006; Umlauf et al., 2004). However, it was unclear when differential *Kcnq1ot1* imprinted domain regulation is first established and how this leads to tissue-specific differential imprinted expression in mid-gestation conceptuses. Three possible scenarios can be envisaged. Firstly, during preimplantation development, all imprinted genes (inner and outer) in the domain are paternally silenced in the three lineages, and then during postimplantation development, paternal alleles of outer genes are reactivated in embryonic tissues (i.e. loss of imprinted expression) (Lewis et al., 2006; Umlauf et al., 2004). Secondly, during preimplantation development, only inner imprinted genes are paternally silenced in the three lineages, and then during postimplantation development outer genes gain paternal allelic silencing in extraembryonic tissues (i.e. gain of imprinted expression). Thirdly, lineage-specific paternal silencing of imprinted genes is established during preimplantation development with further modifications during postimplantation development. Data from this study primarily support the latter scenario. With respect to imprinted expression of outer genes in the *Kcnq1ot1* imprinted domain in control cells, paternal allelic silencing was seen for *Osbpl5* in XEN and ES but not TS cells; *Cd81* in XEN and TS but not ES cells; *Ascl2* in TS but not XEN and ES cells; and *Th* in XEN but not ES and TS cells (Fig. S24). Furthermore, *Nup107, Nup62* and *Nup153* depletion did not result in domain-wide loss of paternal allelic silencing in ES, TS or XEN cells, as would have been expected in the first scenario. While all inner genes in the *Kcnq1ot1* imprinted domain underwent paternal silencing in XEN, ES and TS cells, *Nup107*, *Nup62* and *Nup153* depletion showed lineage-specific, paternal allelic reactivation; paternal *Slc22a18, Cdkn1c* and *Kcnq1* alleles in TS and XEN cells, and only paternal *Cdkn1c* in ES cells. Overall, our data provide the strongest support for the third scenario where lineage-specific paternal silencing of imprinted genes has been established during preimplantation and is further augmented in a lineage-specific manner during postimplantation development. Future studies involving transdifferentiation of ES into TS or XEN cells could provide additional insight into lineage-specific *Kcnq1ot1* imprinted domain regulation.

While our work has established lineage-specific regulation of the *Kcnq1ot1* imprinted domain by NUP107, NUP62 and NUP153 in ES, TS and XEN cells, we are still left with the question of when differential *Kcnq1ot1* imprinted domain regulation is first established. To resolve this important question, analyses will be required at the blastomere-level, since morula and blastocyst stage embryos possess more than one cell lineage. Furthermore, future studies are will also be required to investigate the role of NUP107, NUP62 and NUP153 in imprint establishment during gametogenesis.

## MATERIALS AND METHODS

### Stem cell culture and transfection

C57BL/6J female X *Mus musculus castaneus* male ES and TS were produced as described (Golding et al., 2011). ES cells were maintained in DMEM (Sigma, D5671) supplemented with 50 μg/mL penicillin/streptomycin (Sigma, P4333), 1 mM sodium pyruvate (Invitrogen, 11360070), 100 μM β-mercaptoethanol (Sigma, M3148), 1X LIF (Millipore, ESG1106), 2 mM L-glutamine (Sigma, G7513-100), 1X MEM non-essential amino acids (Invitrogen 11140-050) and 15% hyclone ES cell grade fetal bovine serum (FBS) (Fisher Scientific, SH30070.03E). TS cells were maintained in RPMI (Sigma, R0883) supplemented with 50 μg/mL penicillin/streptomycin, 1 mM sodium pyruvate, 100 μM β-mercaptoethanol, 2 mM L-glutamine, 15% hyclone ESC grade FBS, 1 μg/mL heparin (TSCs) (Sigma, H3149), 1X FGF basic (TSCs) (R&D Systems, 233-FB-025) and 1X FGF4 (TSCs) (R&D Systems, 235-F4-025). Prior to experiments, ES and TS cells were cultured on a gelatin-coated (EmbryoMax^®^ 0.1% Gelatin Solution, Millipore, ES-006-B) feeder-free environment to avoid feeder cell contamination. Where specified, cells synchronization in G1 phase was performed by treatment with 2 mM hydroxyurea (Sigma, H8627) for 12 hours, with subsequent siRNA transfection. Transfection was conducted with 10 nmol of siRNAs with 3.6 siPepMute™ siRNA Transfection Reagent (SignaGen, SL100566) in 100 μl of 1X transfection buffer. Two hours before transfection, medium was refreshed. See Table S1 for siRNA sequences. Three ES and TS lines that were obtained from different embryos were used for each experiment with 3–4 technical replicates per line, except for the DNA methylation analyses, where two biological replicates were used.

### RNA isolation, cDNA preparation and PCR amplification

To isolate total RNA, the PureLink^®^ RNA Mini Kit (Invitrogen, 12183018A), RNeasy Plus Mini Kit (Qiagen, 74134), or QuickRNA™ MicroPrep (Zymo Research, R1050) were used as specified in the manufacturers’ instructions. Total RNA was then DNase I (Life Technologies, 18068015) treated as described (Golding et. al., 2011). ProtoScript II Reverse Transcriptase (NEB, M0368X) was used to synthesize cDNA using oligo(dT)23 (Sigma, O4387) and Random Primers (Life Technologies, 48190011), after which any residual RNA was removed from cDNA through RNaseA (Roche) treatment. PCR amplification was performed on C1000 and MJ Research Thermocyclers (BioRad). Table S2 lists primers, annealing temperatures and amplicon sizes.

Quantitative PCR (qPCR) amplification was executed with iQ SYBR Green Supermix (BioRad, 1708880) or SensiFAST™ SYBR^®^ No-ROX Kit (Bioline, BIO-98005) on a MJ Thermocycler Chromo4 Real-time PCR system (BioRad, CFB-3240) or a CFX-96™ Real-time system (BioRad, 185-5096). Primers, annealing temperatures and amplicon sizes are listed in Table S2. For gene expression analysis, data were analyzed using the 2ΔΔCT method (Schmittgen and Livak, 2008).

For droplet digital PCR, RNA extraction was performed with the RNeasy Plus Mini Kit (Qiagen, 74134) as specified by the manufacturers’ instructions. Qubit^®^ 3.0 Fluorometer (Life Technologies, Q33216), Qubit RNA HS Assay Kit (Thermo Fisher Scientific, Q32852), and Qubit RNA BR Assay Kit (Thermo Fisher Scientific, Q10210) were used to determine RNA concentrations. Total RNA was treated with DNase I (Life Technologies, 18068015), and then cDNA synthesis was performed using ProtoScript II Reverse Transcriptase (NEB, M0368X), oligo(dT)23 (Sigma, O4387) and Random Primers (Life Technologies, 48190011). After preparation, cDNA was subjected to RNaseA (Roche, 10109142001) treatment to eliminate any residual RNA. Droplet digital PCR master mixes consisted of 250 nM primers each, 250 nM FAM and HEX probes, Supermix for Probes (no dUTP) (Bio-Rad,1863024), cDNA plus DNAse-RNAse free UltraPure dH20 (Invitrogen, 10977-015) in a 22 μl final reaction volume. Reactions were loaded into a 96-well plate (Bio-Rad, 12001925), and then sealed with a foil heat seal (Bio-Rad, 1814040) on a PX1 PCR Plate Sealer at 180°C for 5 seconds. Droplet generation was performed on the AutoDG (Bio-Rad, 1864100). Droplets were transferred into a new 96-well plate (Bio-Rad, 12001925) and foil heat sealed (Bio-Rad, 1814040). Droplet digital PCR was conducted on a deep well CFX1000 cycler (BioRad,1851197) as described (Sachani et al., 2018). Table S1 lists primer and probe sequences along with optimal annealing temperature. Droplets reading was performed on the QX200 (Bio-Rad, 1864100), which was set to detect absolute levels of FAM/HEX probe fluorophores. Droplets with FAM and HEX containing alleles were selected with the Quantasoft Analysis Pro 1.0.596 (Bio-Rad) 1D and 2D Amplitude tools, eliminating any FAM/HEX double-positive droplets containing both alleles. Copies/μ,g were plotted using BoxPlotR (http://shiny.chemgrid.org/boxplotr/).

### RNA/DNA fluorescence *in situ* hybridization

The Wl1-2505B3 fosmid (32-kb) for the *Kcnq1* intronic region was obtained from BACPAC Resources Center at the Children’s Hospital Oakland Research Institute. Kcnq1ot1RNA/DNA FISH probes were labelled with the BioPrime DNA labeling System (Thermo Fisher Scientific, 18094011) and fluorescein-12-dUTP (Roche, 11373242910) and Biotin-16-dUTP (Roche, 11093070910) for RNA FISH, and with Cy5-UTP (GE Healthcare, 45001239) for DNA FISH, as described (Golding et al., 2011). To conduct RNA FISH, cells were seeded on coverslips (VWR, 48380046) in 12-well plates (Corning, C3512) and then fixed through a dehydration series of 25%, 50%, 75% and 100% ethanol, following which hybridization was carried out with RNA-FISH probes in 100% molecular grade formamide (VWR, 97062-006), as described (Golding et al., 2011). Briefly, coverslips were incubated overnight at 37°C in a formamide sealed hybridization chamber, and then washes were carefully performed in 4X SSC and 2X SSC (Thermo Fisher Scientific, AM9763) at 37°C for 5 minutes 3 times each. Coverslips were first treated for the primary antibody in blocking buffer (20X SSC, 10% Tween-20 (Sigma, P9416), 10% Skim Milk, 37°C, 1 hour in the dark), and then washed 3 times in 4X SSC buffer with shaking (37°C), in blocking buffer with the secondary antibody (37°C in the dark for 1 hour), 3 times in 4X SSC buffer (37°C, 5 minutes each), and then twice in 2X SSC buffer (37°C, 5 minutes each) with continuous agitation. A final wash was performed (1X SSC, 5 minutes). Coverslips were mounted with Vectashield DAPI (Vector labs, H-1000) on glass slides, and then housed in the dark at 4°C for several hours to overnight. For DNA/RNA FISH, we first conducted DNA FISH as described (Korostowski et al., 2011) after which RNA FISH was performed. Briefly, fixed cells were rinsed (1X PBS) and dehydrated (ethanol washes), quickly washed (2X SSC), and then hybridized with the denatured DNA FISH probe overnight (37°C in a parafilm sealed formamide chamber). On the following day, excess DNA probe was washed off (pre-warmed 42°C 4X SSC and 2X SSC buffer, 3 times each), and then RNA hybridization was performed as described above. Z-stack images were acquired on a FluoView FV1000 with an IX81 motorized inverted system (Olympus). For DNA and RNA FISH, Volocity (PerkinElmer) was used to measure signal volume with z-stacks intensity-based thresholding (p,m^3^), as well as the distance of the DNA FISH signal centroid from the nuclear rim to determine domain positioning (p,m) (ImageJ). Table S3 lists antibody and their dilutions.

### Chromatin immunoprecipitation assay

ChIP was performed as described (Sachani et al., 2018). Briefly, cells were collected from control and siRNA treatment groups (~0.5–1.2 million cells). Cells (~40,000 per sample) were then crosslinked in freshly prepared, 1% final concentration formaldehyde for 6 minutes, quenched for 5 minutes in 1.25 M glycine, and then washed in 1X PBS. Pellets were resuspended in cytoplasm extraction buffer (60 mM KCl, 1 mM PMSF, 10 mM HEPES (pH 7.6), 1 mM DTT, 1 mM EDTA, 0.075% NP40), incubated for 5 minutes on ice and then spun at 3000 rpm at 4 °C for 4 minutes. Nuclear pellets were lysed by resuspension in 1% SDS buffer for 10 minutes on ice, and then 1X TE, proteinase inhibitor (Sigma, P8340). Sonication (Covaris M220 or Diagenode, B01060001) was used to shear chromatin into 100–400 bp fragments, which was followed by immunoprecipitation. Frozen, crosslinked chromatin lysates were stored at −80 °C. Protein G Dynabeads (Life Technologies) were washed with ChIP dilution buffer (1% Triton X-100, 0.1% SDS, 2 mM EDTA, 500 mM NaCl, 20 mM Tris pH 8.1) three times, bound with antibody at 4 °C for 1 hour, and then precleared chromatin lysate was incubated for 3 hours to overnight as specified in Table S3. Antibody-bound Dynabeads were washed at 4 °C once in Low Salt Buffer (1% Triton X-100, 0.1% SDS, 2 mM EDTA, 150 mM NaCl, 20 mM Tris pH 8.1) for 5 minutes, and then twice in High Salt Buffer (1% Triton X-100, 0.1% SDS, 2 mM EDTA, 500 mM NaCl, 20 mM Tris pH 8.1) for 5 minutes each. A final wash was performed in 1X TE Wash Buffer (10 mM Tris pH 8.1, 1 mM EDTA) at 4 °C for 5 minutes. DNA was eluted from the Dynabeads through a 65 °C incubation for 1 h with elution buffer (1% SDS,100 mM NaHCO_3_). To purify eluted DNA, RNA was degraded by RNase A (Roche) treatment for 1 hour at 37 °C, while protein was degraded with proteinase K (Sigma) for 2 hours at 55 °C. Following ChIP, DNA was extracted with Chelex beads (Bio-Rad, 1421253), QIAquick PCR Purification Kit (Qiagen, 28104), or ChIP DNA Clean & Concentrator™ (Zymo Research, D5201). ChIP-qPCR amplification was conducted with iQ SYBR Green Supermix (Bio-Rad, 1708880) and SensiFAST™ SYBR^®^ No-ROX Kit (Bioline, BIO-98005) on an MJ Thermocycler Chromo4 Real-time or CFX-96 Real-time (Bio-Rad) PCR systems. Enrichment levels (% input) were calculated 100×(2^^^[ΔCt_Input_ − ΔCt_Input_] − [ΔCt_Input_ − ΔCt_Ab_])/25. Data were presented as total enrichment (percent input) or allelic enrichment (total enrichment times allelic enrichment ratio).

### Statistical analysis

All statistical analysis was performed using GraphPad Prism. A two-tailed Student’s t-test was performed on mean values with treatment samples compared to controls. p-values were adjusted for multiple comparisons. Significance was considered to have a p-value less than 0.05, or for the droplet digital PCR allelic analysis at p-value less than 0.01.

## ACKNOWLEDGMENTS

Support for this work was provided by grants from the Canadian Institutes of Health Research (MOP 111210), Magee-Womens Research Institute and the University of Pittsburgh to MRWM. SSS was supported by Magee-Women’s Research Institute Paul M. Rike Postdoctoral Fellowship. CRW was supported by Ontario Graduate Scholarship. MRWM was a Magee Auxiliary Research Scholar.

## CONTRIBUTIONS

SSS performed experiments and wrote manuscript. WAM designed droplet digital PCR assays and conducted bioinformatics analysis. AMF performed Western analysis and immunoprecipitation. CRW performed bisulfite mutagenesis assay. LZ performed Western blot protein assay. MRWM designed and supervised the project, assisted with data interpretation, as well as manuscript preparation. All authors have read and approved the final manuscript

## SUPPLEMENTAL METHODS

### RNA stability assay

*Kcnq1ot1* half-life studies were performed by addition of 2 μg/mL of actinomycin D (Sigma A9415) to control and nucleoporin-depleted ES and TS cells for 4 hours according to Hazan-Halevy et al. (2010). Cells were collected at 0 hours and every hour time-intervals up to 12 hours. RNA was extracted, and cDNA was synthesized as described in the manuscript. *Kcnq1ot1* ncRNA levels were normalized to time 0 hours for each treatment.

### Nuclear transport

ES and TS cells were were transfected with 3 μg of E47-RFP^NLS^ construct using Lipofectamine2000 (Invitrogen, 11668027), followed by siRNA transfection. As a positive control, ES and TS cells were treated with 10 μM of ivermectin (Sigma, I8898) for 48 hours. To access growth rates, ~25 000 cells were seeded on a 6-well plate (Corning, C3506), and then transfected 12 hours later with siRNAs. For nuclear export, biotin-labeled oligo-dT-50 (Life Technologies, custom primer design) was used for polyA-mRNA FISH as described in the manuscript.

### ES and TS cell growth rate and doubling time

Direct cell counts were performed every 12 hours using a hemacytometer (VWR, 15170–208). Population doubling time (DT) was calculated using the following equation, DT = T * [ln (2) / ln (Xe/Xb)] where X_B_ is the cell number at the beginning of the incubation, X_E_ is the cell number at the end of the incubation time (or collection).

### Western blot analysis and immunoprecipitation assay

Cytoplasm and nuclear protein extracts were isolated using cytoplasm extraction buffer and nuclear extraction buffer using the NE-PER Nuclear and Cytoplasmic Extraction Kit (ThermoFisher Scientific, 78833) followed by western blot analysis as described (Golding et al. 2010; Baldwin 1996). Briefly, protein samples were quantified using the Bradford assay, Nanodrop or Qubit protein concentration determination assay. Typically, 20–30 μg of protein was separated on an 8–12% SDS-PAGE (Bio-Rad, Bulletin 6201) gel or any kD™ Mini-PROTEAN^®^TGX™ Precast Protein Gels (Bio-Rad, 4569033), transferred to PVDF membrane (Bio-Rad, 1620177) or Trans-Blot^®^ Turbo™ Mini PVDF Transfer Packs (Bio-Rad, 1704156), blocked for 1 h in 5% skim milk in 1X TBST (1X TBS (Bio-Rad, 1706435) with 0.1% Tween-20, (Sigma, P9416)), incubated with primary antibody in 5% skim milk as described in Table S3 followed by three washes of 1X TBST for 7 min each. Blots were next incubated with secondary antibody in 5% skim milk followed by detection using chemiluminescence reaction with Clarity Western ECL Substrate (Bio-Rad, 1705060). See Table for antibodies and dilutions.

Protein co-immunoprecipitation was performed using Dynabeads Protein G (Invitrogen, 10004D) as per the manufacturer’s instructions. Briefly, 50 μg of nuclear lysates obtained from ES and TS cells using the NE-PER Nuclear and Cytoplasmic Extraction Kit (ThermoFisher Scientific, 78833) were incubated with the respective antibody, 2 μL RNase A (Roche, 10109142001), 2 μL DNase I (Life Technologies, 18068015), and 4μL protease inhibitor (Sigma, P8340) as described in Table S3 overnight at 4 °C. Samples were then incubated with Dynabeads protein G (Invitrogen) for 1 h at 4 °C. Immunoprecipitates were washed three times with 200μLof 0.1% Tween-20 in 1×PBS, eluted in 1X Laemmli buffer (Bio-Rad, 1610747) and resolved on 10–12% SDS-PAGE followed western blot analysis as described above.

### Bisulfite mutagenesis and sequencing

Control and siRNA-treated ES and TS cells (20% confluent) were seeded on gelatin-coated 6-well dishes. Forty-eight hours after transfection, cells were washed once with 1X PBS (Sigma) followed by a 5-minute incubation with 1X Trypsin-EDTA (Sigma, 59418C) in PBS. Trypsin was inactivated by addition of RPMI medium. Detached cells were collected and pelleted gently at 200 RCF for 5 minutes, washed and re-suspended in 1X PBS. One percent of cells (~10,000 cells) were embedded into a 2:1 3% LMP agarose (Sigma, A9414) and lysis solution [100 mM Tris-HCl, pH 7.5 (Bioshop, TRS002.1), 500 mM LiCl (Sigma, L4408), 10 mM EDTA, pH 8.0 (Sigma, 324506), 1% LiDS (Bioshop, LDS701), and 5 mM DTT (Sigma, 43816), 1 μLof 2mg/ml proteinase K (Sigma, P4850), and 1 μL 10% Igepal (Sigma, I8896)] as described (Denomme et al. 2011). Briefly, the agarose-sample mixture was then placed on ice for 10 minutes to produce an agarose/lysis bead. Samples were then incubated overnight for 20 hours in SDS lysis buffer [450 μLTE pH 7.5, 50 μL 10% SDS, 1 μL proteinase K] at 50°C, following which lysis buffer was removed and 300 μL of mineral oil was added to the top of each ES or TS cell/bead mixture. Samples were either processed immediately for bisulfite mutagenesis or frozen at −20°C for a maximum of 5 days. Bisulfite mutagenesis was performed as described for *Kcnq1ot1* amplification in ES and TS cells (Denomme et al. 2011). Samples were first incubated at 90°C to heat inactivate the proteinase K for 2.5 minutes, then transferred to ice for 10 minutes. To denature DNA, 0.1 M NaOH solution was added at 37°C for 15 minutes. For bisulfite conversion, samples were covered with 300 μL of mineral oil, under which 500 μL of 2.5 M bisulfite solution was added. Samples were incubated for 3.5 hours at 50°C, following which desulfonation was completed in 1 mL of 0.3 M NaOH at 37°C for 15 minutes. Two washes each were performed in TE pH 7.5 and autoclaved water. Negative controls (beads without the embedded cells) were processed with each bisulfite reaction. For first round PCR amplification, the agarose bead with bisulfite converted DNA (10 μL) was added to 15μL of Hot Start Ready-To-Go PCR bead (GE Healthcare) containing 0.2 μM *Kcnq1ot1* external primers, 9.6 ng/mL transfer RNA with a 25 μL mineral oil overlay. For second round PCR, first round PCR product (5 μL) was added to PCR beads containing 0.2 μM *Kcnq1ot1* internal primers. See Supplementary Table 4 for primers. PCR products were ligated into pGEM-Easy vector (Promega) as per the manufacturer’s instructions and sequenced at the BioBasic Sequencing Facility (Markham, Canada). Sequences with less than 90% conversion were excluded. Percent methylation was calculated as the number of methylated CpGs over the total number of CpGs.

---

**Table S1:** siRNA transfection details

| siRNA Target | Type | Supplier | Catalog | Concentration | Transfection Duration |
| --- | --- | --- | --- | --- | --- |
| Nup107-A | SMARTpool ON-TARGETplus | Dharmacon | L-065221-01-0005 | 10 nmol | 48 h |
| Nup62-A | SMARTpool ON-TARGETplus | Dharmacon | L-064100-01-0005 | 10 nmol | 48 h |
| Nup98-A | SMARTpool ON-TARGETplus | Dharmacon | L-060137-01-0005 | 10 nmol | 48 h |
| siNT-A | SMARTpool ON-TARGETplus | Dharmacon | D-001810-01-20 | 10 nmol | 48 h |
| Nup107-B1 | Silencer Select siRNA | Life Technologies | s98083 | 10 nmol | 48 h |
| Nup107-B2 | Silencer Select siRNA | Life Technologies | s98085 | 10 nmol | 48 h |
| Nup62-B1 | Silencer Select siRNA | Life Technologies | s70887 | 10 nmol | 48 h |
| Nup62-B2 | Silencer Select siRNA | Life Technologies | s70885 | 10 nmol | 48 h |
| Nup98-B1 | Silencer Select siRNA | Life Technologies | s114479 | 10 nmol | 48 h |
| Nup98-B2 | Silencer Select siRNA | Life Technologies | s114477 | 10 nmol | 48 h |
| Nup153-A | Silencer Select siRNA | Life Technologies | s104224 | 10 nmol | 48 h |
| Nup153-B | Silencer Select siRNA | Life Technologies | s104225 | 10 nmol | 48 h |
| siNT-B | Silencer Select siRNA | Life Technologies | 4390847 | 10 nmol | 48 h |

**Table S2:**
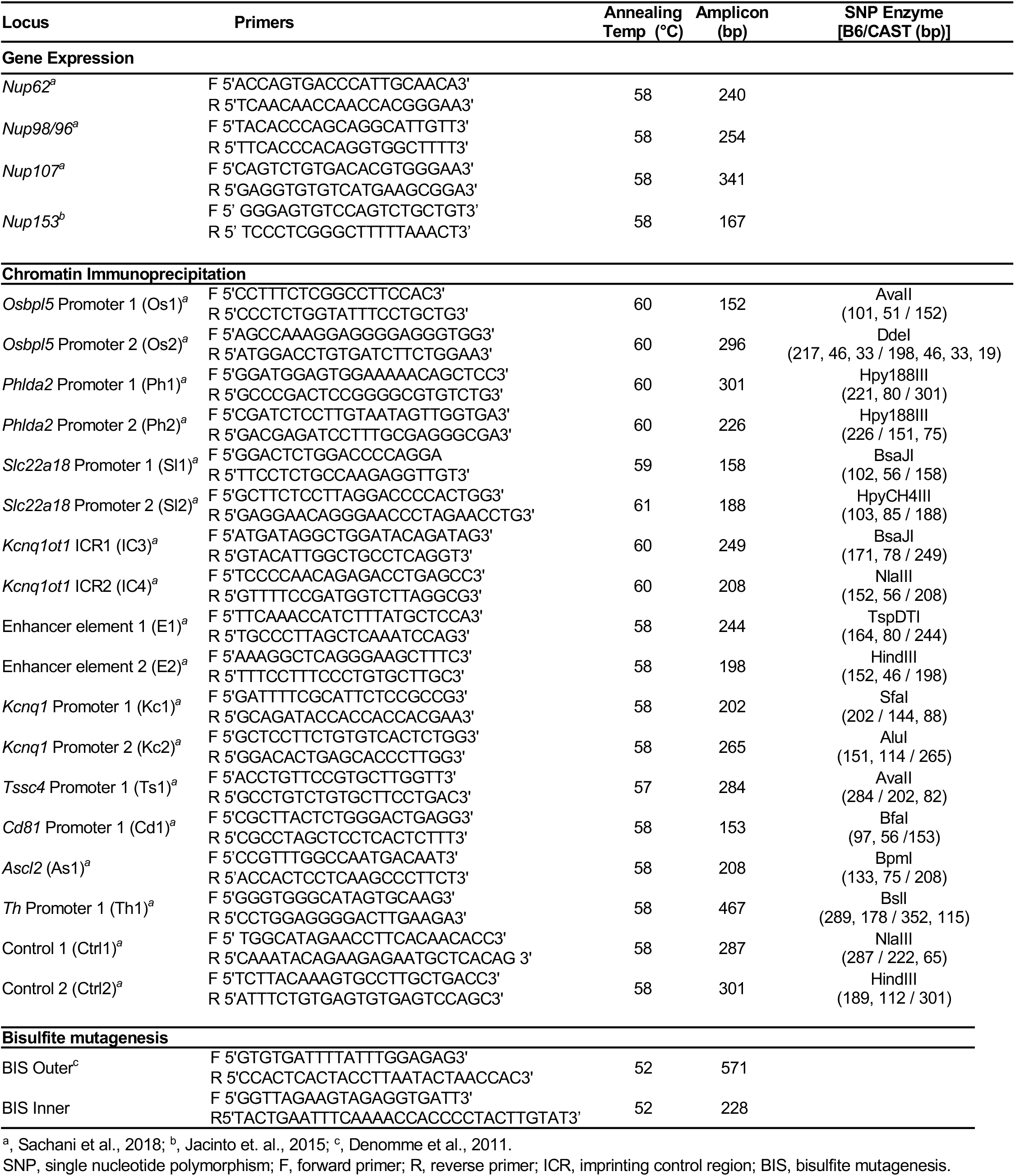
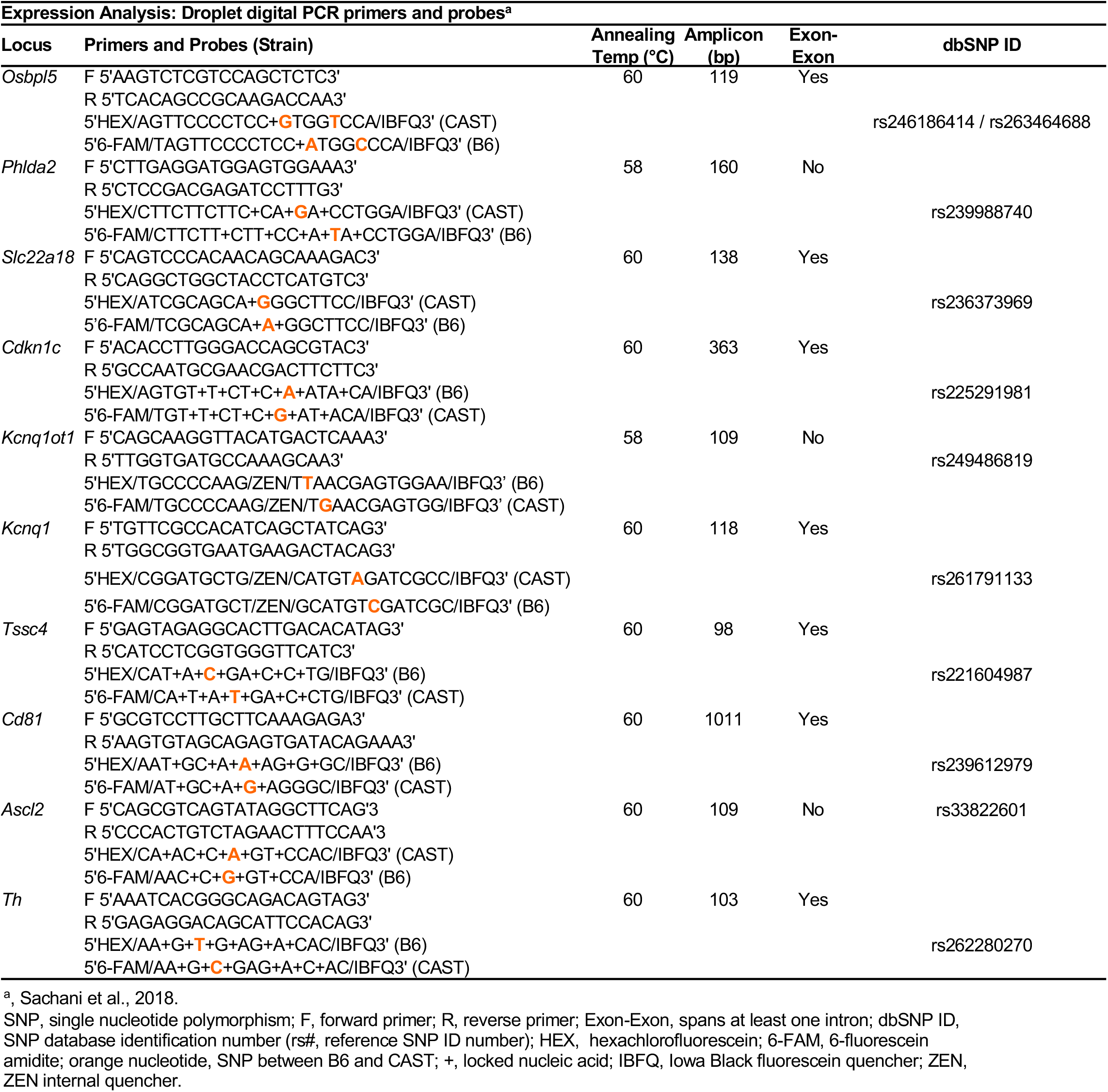
Primers and PCR conditions continued

**Table S3:** Antibodies and Probes

| **Antibody** | **Catalogue Number** | **Supplier** | **Concentration/**<br>**Dilution** | **Duration** | **Temp (°C)** |
| --- | --- | --- | --- | --- | --- |
| **Western** |  |  |  |  |  |
| mAb414 | ab24609 | Abcam | 1:1000 | overnight | 4 |
| NUP98 | ab50610 | Abcam | 1:1000 | overnight | 4 |
| NUP107 | ab73290 | Abcam | 1:1000 | 1 hr | RT |
| NUP153 | sc-101545 | Santa Cruz Biotech | 1: 1000 | 1 hr | RT |
| NUP50 | ab137092 | Abcam | 1:1000 | 1 hr | RT |
| NUP93 | ab168805 | Abcam | 1:1000 | 1 hr | RT |
| NUP160 | ab74147 | Abcam | 1:1000 | 1 hr | RT |
| NUP358 | ab2938 | Abcam | 1:1000 | 1 hr | RT |
| AHCTF | A300-166A | Bethyl Laboratories | 1:1000 | 1 hr | RT |
| TPR | sc-271565 | Santa Cruz Biotech | 1:1000 | 1 hr | RT |
| H3 | ab1791 | Abcam | 1:5000 | 1 hr | RT |
| INCENP | I5283 | Sigma-Aldrich | 1:10,000 | 1 hr | RT |
| GFP | G46-66M | SignalChem | 1:3000 | overnight | 4 |
| SMC1A | A300-055A | Bethyl Laboratories | 1:3000 | overnight | 4 |
| SMC3 | A300-060A | Bethyl Laboratories | 1:3000 | overnight | 4 |
| α-TUBULIN | sc-8035 | Santa Cruz Biotech | 1:7000 | 1 hr | RT |
| Anti-Mouse-HRP Secondary | SC-2314 | Santa Cruz Biotech | 1:5000 | 1 hr | RT |
| Anti-Rat-HRP Secondary | SC-2956 | Santa Cruz Biotech | 1:4000 | 1 hr | RT |
| Anti-Rabbit-HRP Secondary | G33-62G | SignalChem | 1:6000 | 1 hr | RT |
| **Immunohistochemistry/RNA FISH** |  |  |  |  |  |
| LaminB1 (S-20) | sc-30264 | Santa Cruz Biotech | 1:1000 | 1 hr | 37 |
| Anti-Biotin | ab1227 | Abcam | 1:1000 | 1 hr | 37 |
| Anti-Rabbit Alexa Fluor® 488 | A-11034 | Life Technologies | 1:1000 | 1 hr | 37 |
| Anti-Goat Alexa Fluor® 594 | A-11080 | Life Technologies | 1:1000 | 1 hr | 37 |
| **Chromatin Immunoprecipitation / Immunoprecipitation** |  |  |  |  |  |
| H3 | ab1791 | Abcam | 2 µg | overnight | 4 |
| H3K4me3 | 39159 | Active Motif | 3 µg | overnight | 4 |
| H3K9me2 | ab1220 | Abcam | 3 µg | overnight | 4 |
| H3K27me3 | 07-449 | Millipore | 3 µg | overnight | 4 |
| RNAPII | sc-889 | Millipore | 3 µg | 3 hr | 4 |
| mAb414 | ab24609 | Abcam | 4 µg | overnight | 4 |
| NUP98 | ab50610 | Abcam | 4 µg | overnight | 4 |
| NUP153 | sc-101545 | Santa Cruz Biotech | 4 µg | overnight | RT |
| SMC1A | A300-055A | Bethyl Laboratories | 1.5 µg | overnight | 4 |
| SMC3 | A300-060A | Bethyl Laboratories | 1.5 µg | overnight | 4 |
| KMT2A | 39829 | Active Motif | 3 µg | overnight |  |
| EHMT | ab41969 | Abcam | 3 µg | overnight |  |
| EZH2 | 07-689 | Millipore | 3 µg | overnight |  |
| Anti-Mouse IgG | sc-2029 | Santa Cruz Biotech | * | * | * |
| Anti-Rabbit IgG | 2729s | Cell Signalling | * | * | * |
| Anti-Goat IgG | sc-2028 | Santa Cruz Biotech | * | * | * |

\* same as corresponding antibody used

**Table S4:**
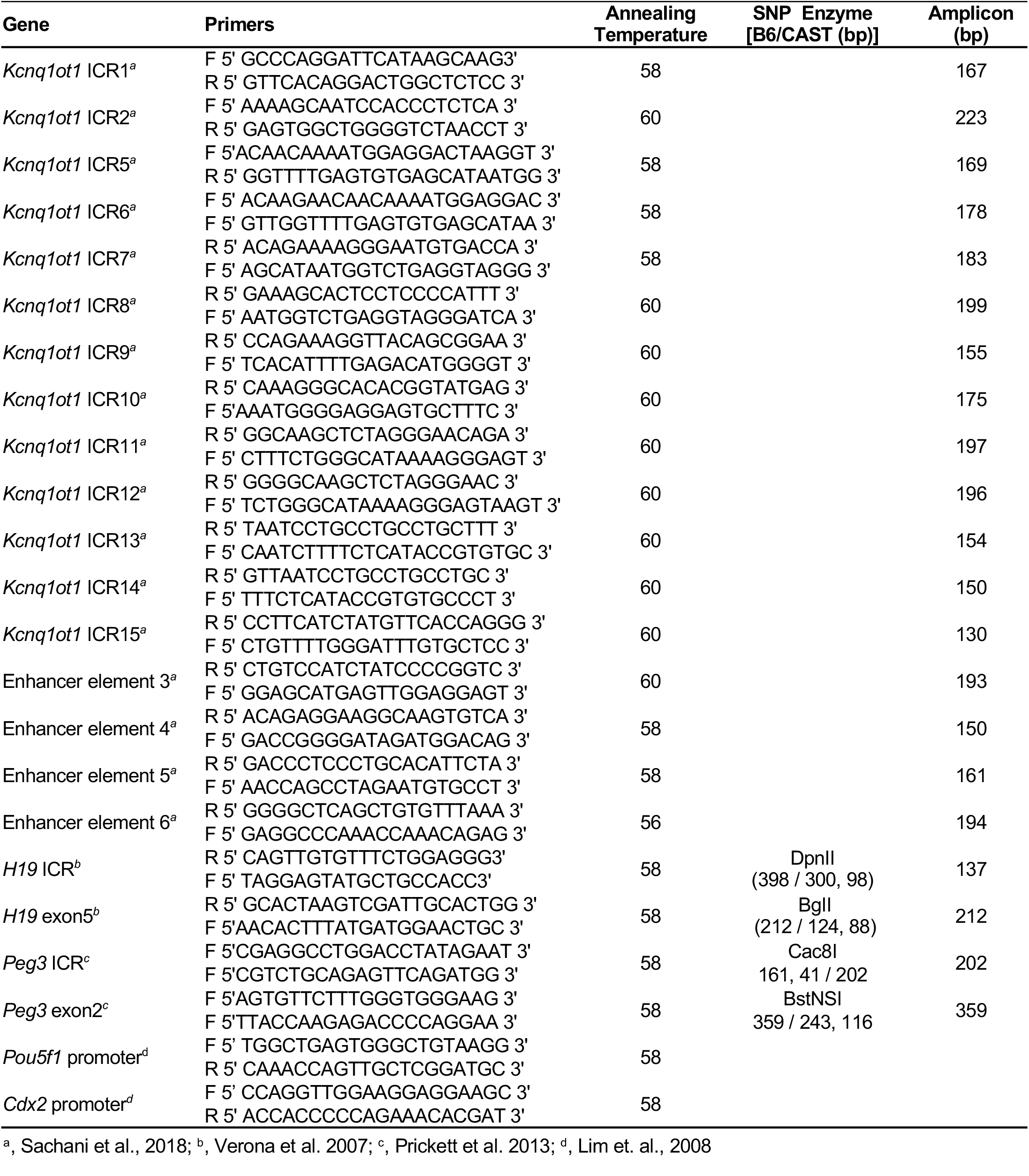
List of ChIP primers for supplementary information

**Figure S1:**
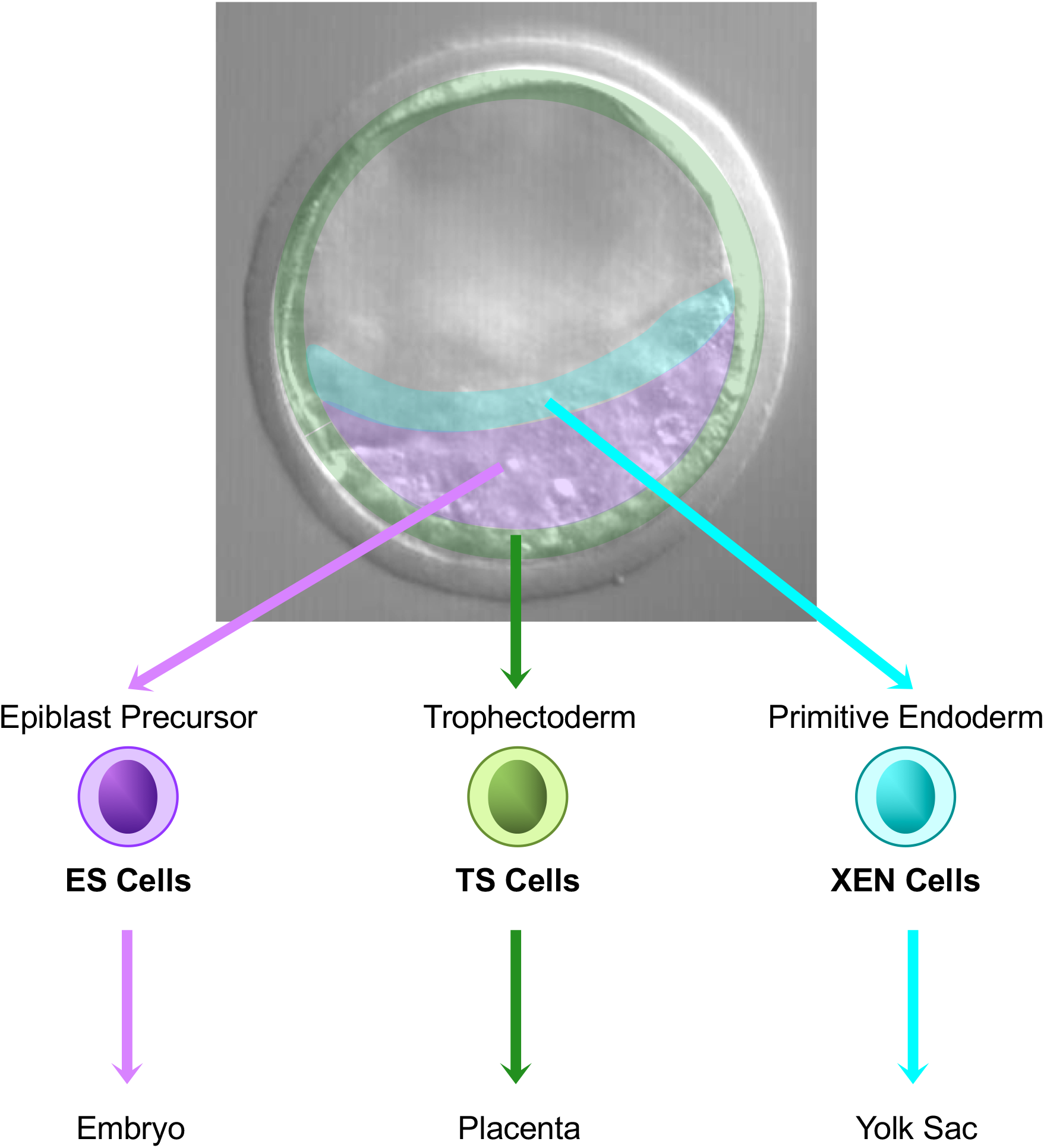
Model for NUP107, NUP62 and NUP153-mediated regulation of the paternal *Kcnq1ot1* imprinted domain in XEN, ES and TS cells. (A) In blastocyst stage embryos, there are three cell lineages, epiblast precursor, trophectoderm and primitive endoderm cells, which give rise to the fetus, placenta and yolk sac, respectively. Stem cells can be generated as embryonic (ES), trophoblast (TS) and extraembryonic endoderm (XEN) stem cells.

**Figure S2.**
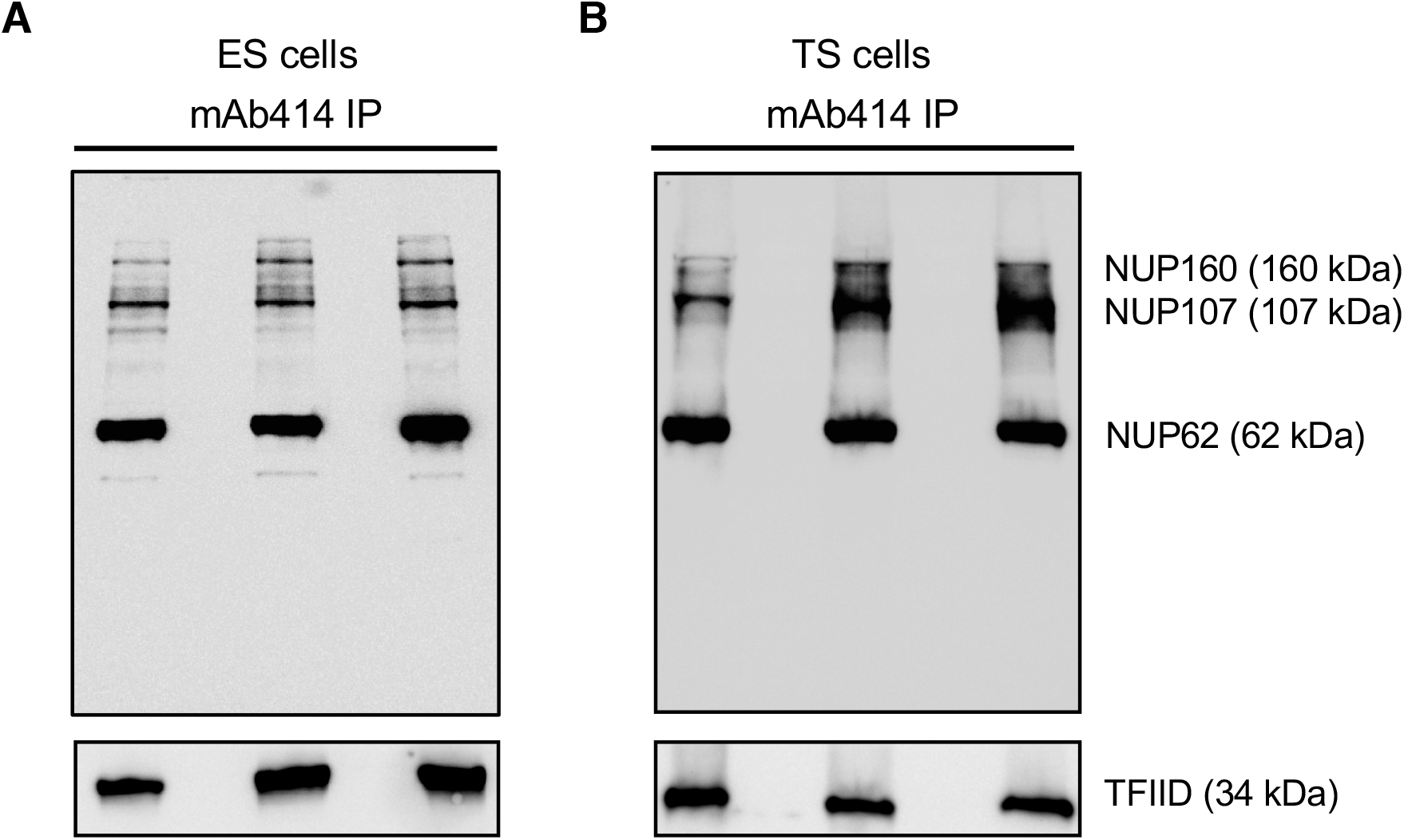
Antibody validation in ES and TS cells. mAb414 IP was performed followed by Western blot analysis using the same mAb414 antibody in (A) ES and (B) TS cells. Representative blots for 3 biological samples are shown. The most prominent nucleoporins detected with the mAb414 antibody in ES and TS cells were NUP62, NUP107 and NUP160.

**Figure S3.**
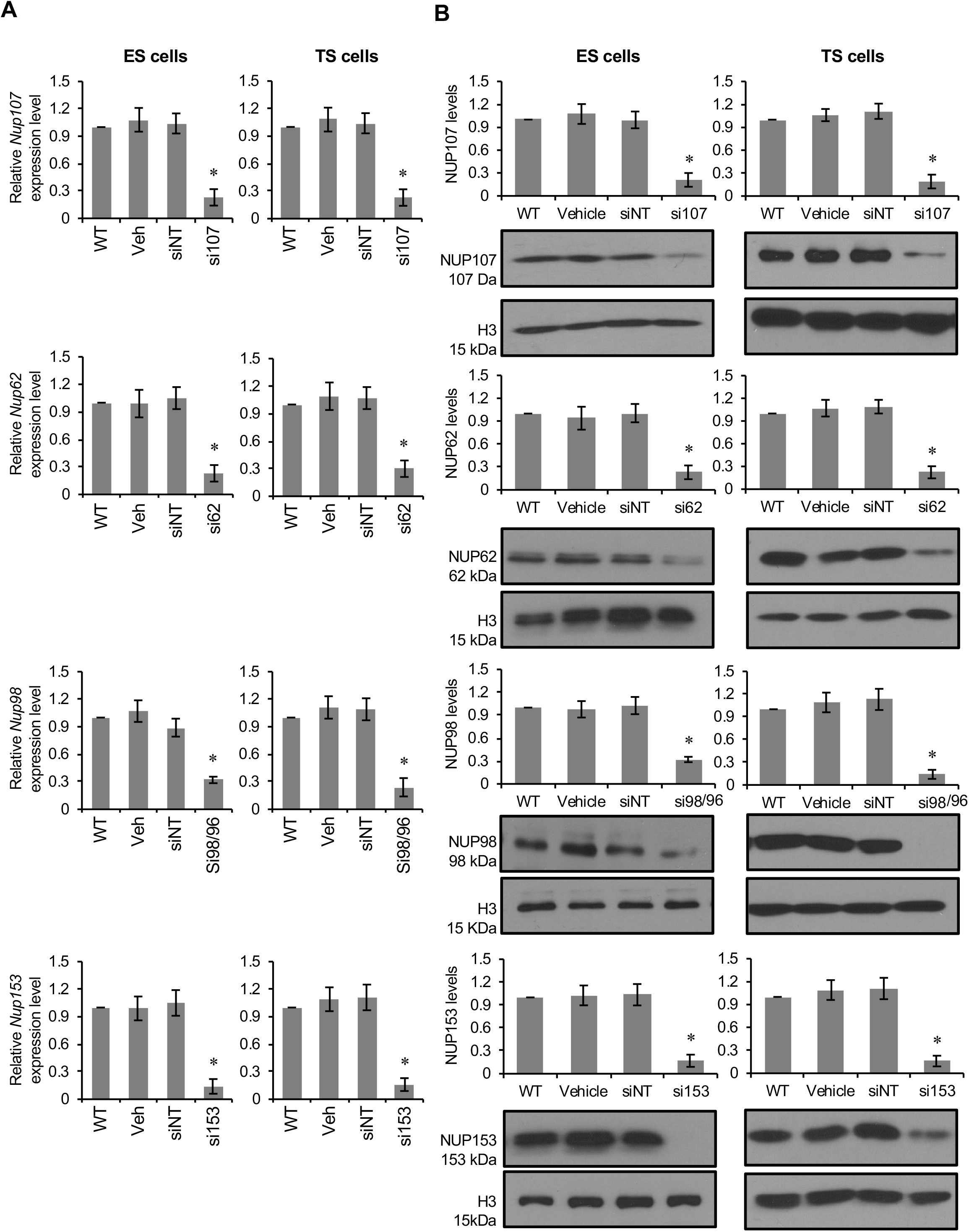
Nucleoporin depletion levels in ES and TS cells. (A) Nucleoporin RNA depletion levels in ES and TS cells. Quantitative real-time PCR analysis for *Nup107, Nup62, Nup98/96* and *Nup153* relative to *Gapdh* expression 48 hours after transfection. Transfections were performed using the two different sets of siRNAs previously validated in XEN cells (n=3 biological samples with 3 technical replicates per sample). The *Nup98* gene is a bicistronic gene that encodes for two separate nucleoporins, NUP98 and NUP96, from one mRNA. Since the siRNAs target the mRNA, which will produce both proteins, the siRNAs have been designated si98/96. All subsequent transfections employed both sets of siRNAs. mRNA abundance in ES and TS cells was depleted to 23% and 23% of controls for *Nup107*, to 23% and 30% of controls for *Nup62*, to 32% and 24% of controls for *Nup98/96*, and to 14% and 16% of controls for *Nup153*, respectively. (B) Nucleoporin protein depletion levels in ES and TS cells. Western blot analysis using NUP107, mAb414, NUP98 and NUP153 antibodies was performed 48 hours after transfection. The NUP98 antibody specifically recognized NUP98; no commercial antibody was available for NUP96. Histone 3 (H3) was used as loading control. Transfections were performed using the two different sets of siRNAs (n=3 biological samples with 1 technical replicate per sample). Protein levels in ES and TS cells were depleted to 21% and 19% of controls for *Nup107*, to 23% and 22% of controls for *Nup62*, to 32% and 14% of controls for *Nup98/96*, and to 17% and 16% of controls for *Nup153*, respectively. Error bars, s.e.m.; *, significance p < 0.05 compared to the WT control; Veh, vehicle; siNT, non-targeting siRNA; si 107, *Nup107* siRNA; si62, *Nup62* siRNA; si98/96, *Nup98/96* siRNA; si153, *Nup 153* siRNA.

**Figure S4.**
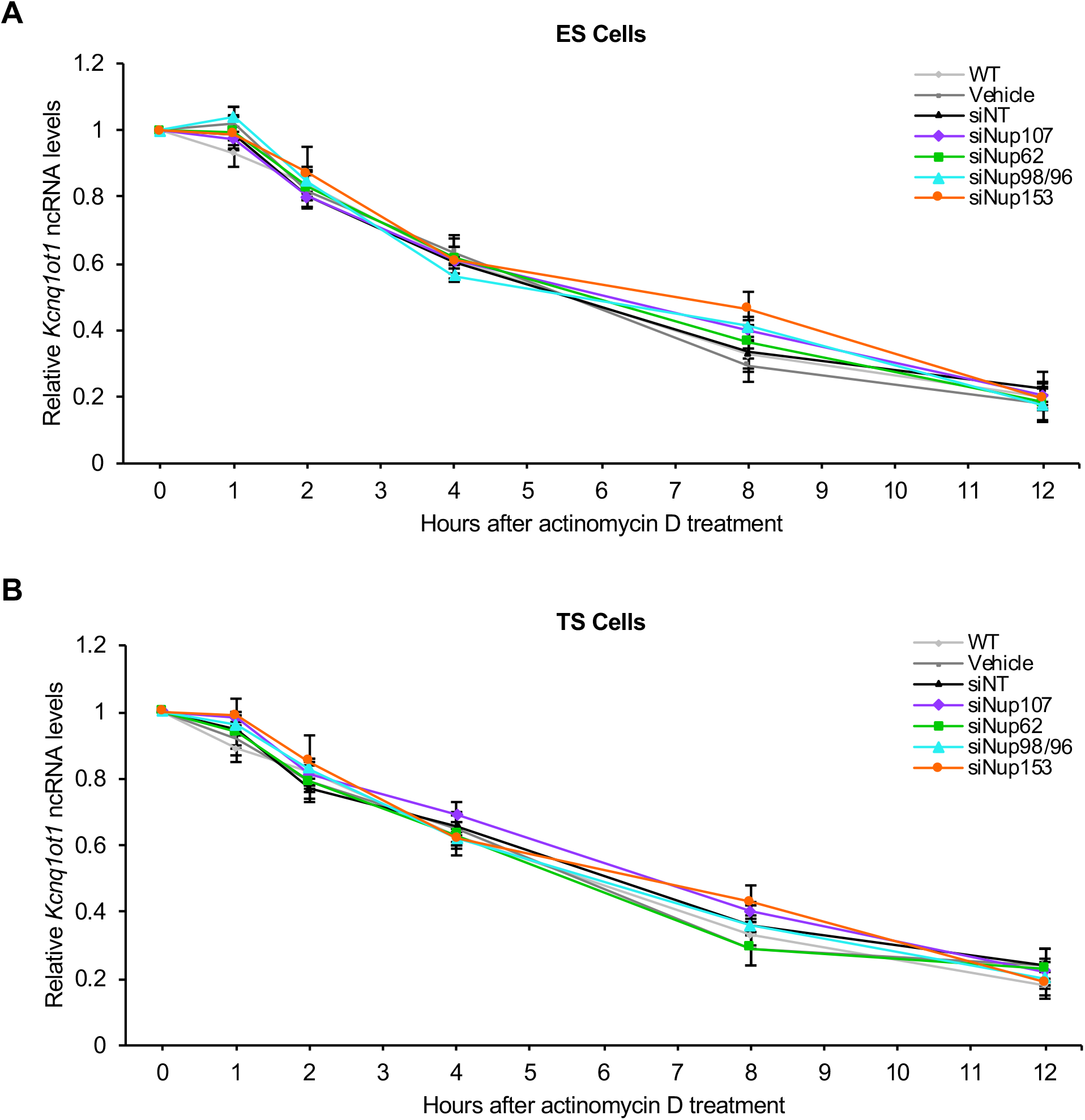
*Kcnq1ot1* ncRNA stability was not altered upon nucleoporin depletion in ES and TS cells. Control and *Nup*-depleted (A) ES and (B) TS cells were treated with actinomycin D for 1 hour, after which cells were collected up to 12 hours after released from treatment. *Kcnq1ot1* expression levels were normalized to 0 hours. No significant changes in *Kcnq1ot1* ncRNA levels were seen at different time intervals after treatment between samples compared to the WT control, indicating that there was no difference in *Kcnq1ot1* ncRNA half-life in control and *Nup*-depleted ES or TS cells (n=3 biological samples with 3 technical replicates per sample). Error bars, s.e.m; *, significance p < 0.05 compared to the WT control; WT, wildtype; Veh, vehicle; siNT, non-targeting siRNA; si107, *Nup107* siRNA; si62, *Nup62* siRNA; si98/96, *Nup98/96* siRNA, si153, *Nup153* siRNA.

**Figure S5.**
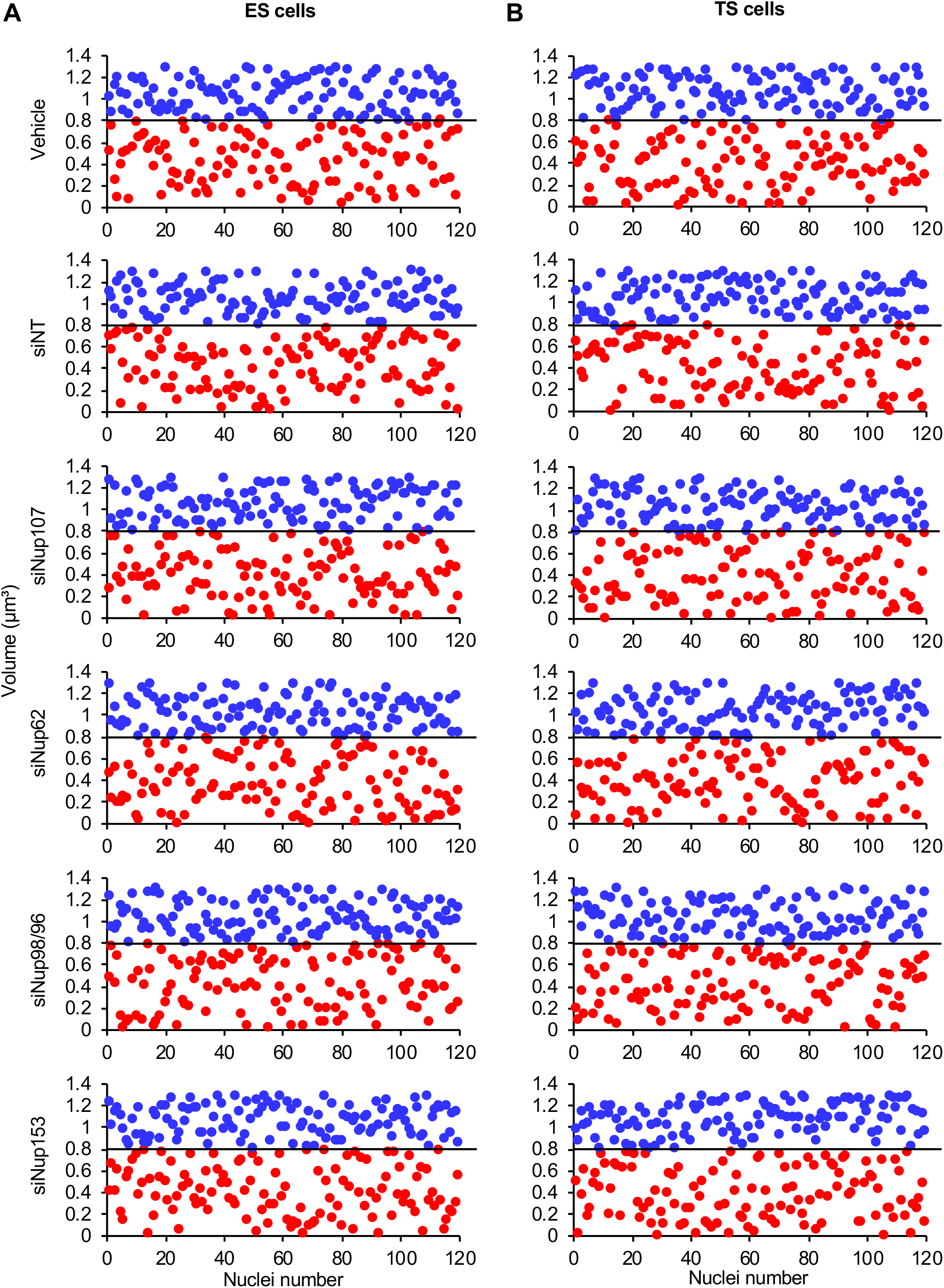
Paternal and maternal *Kcnq1ot1* domain have distinct volumes in ES and TS cells. Paternal (blue, identified by *Kcnq1ot1* ncRNA expression) and maternal (red) *Kcnq1ot1* domain volume were plotted on the Y-axis with the number of G1-synchronized control and *Nup*-*depleted* (A) ES and (B) TS cells plotted on the X-axis. Paternal *Kcnq1ot1* DNA domain volume had a range of >0.8 to 1.3 μm^3^ while maternal *Kcnq1ot1* DNA domain volume ranged from 0.1 to 0.8 μm^3^ (black bar, 0.80 μm^3^; n=4 biological samples; n=120.

**Figure S6.**
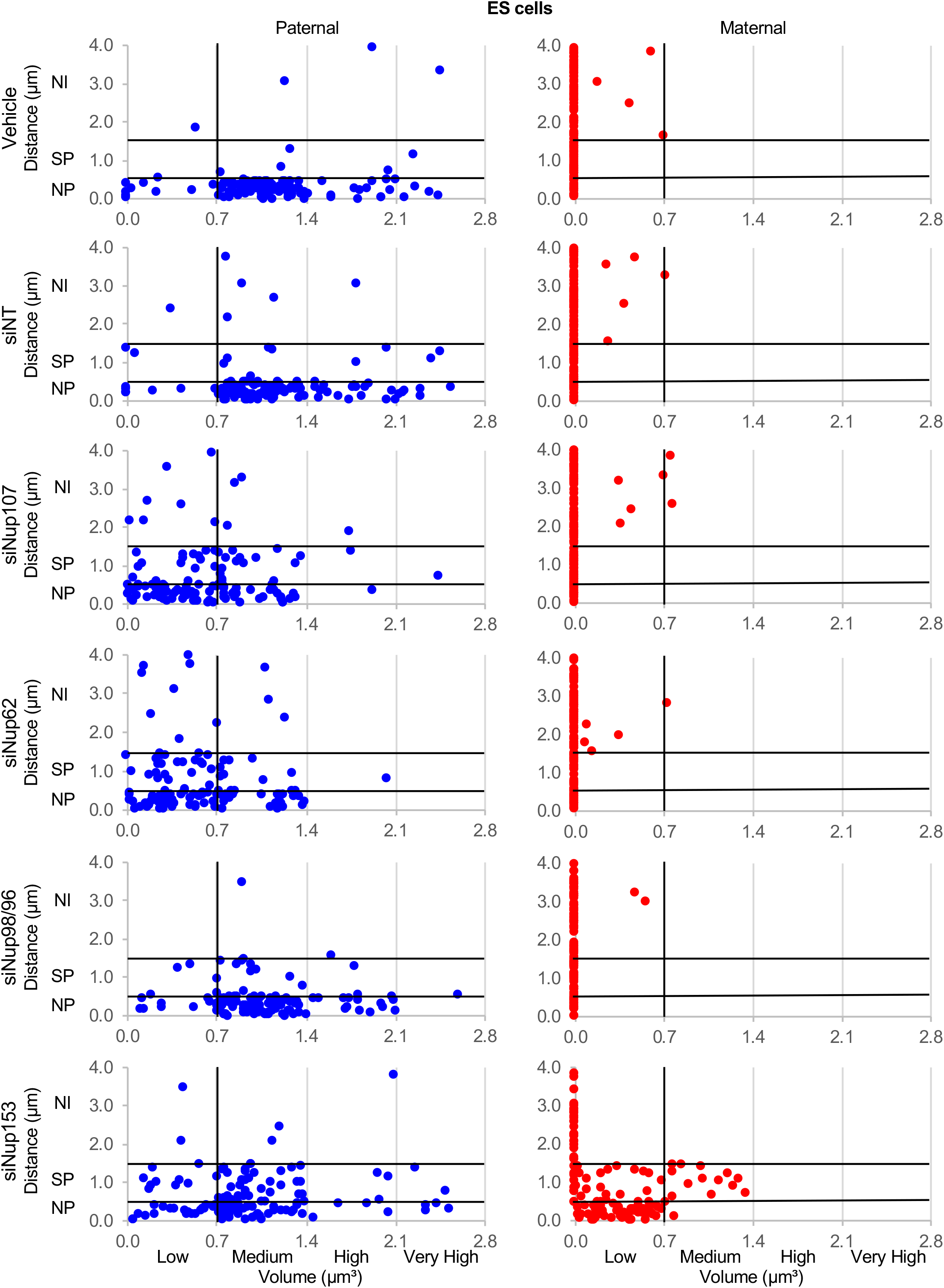
*Kcnq1ot1* ncRNA volume to distance correlation in control and nucleoporin-depleted ES cells. *Kcnq1ot1* ncRNA volume and domain distance from nuclear periphery were plotted on X- and Y-axes, respectively, for G1-synchronized control and *Nup*-*depleted* ES cells. Upon *Nup107, Nup62* and *Nup153* depletion, cells with low paternal *Kcnq1ot1* ncRNA volume (blue) shifted to sub-nuclear peripheral and nuclear interior positions. The maternal *Kcnq1ot1* domain (red) was randomly positioned within the nucleus (expected NP 15%, SP 30%, NI 60%; observed NP 12–17%, 24–34%; NI, 53–59%), except for Nup153-depleted cells. Upon *Nup153* depletion, those cells with maternal *Kcnq1ot1* ncRNA expression (primarily low) had a shift in maternal *Kcnq1ot1* domain positioning toward the nuclear periphery (observed NP 64%, SP 36%, NI, 0%). NP, nuclear periphery; SP, sub-nuclear periphery; NI, nuclear interior; Veh, vehicle; siNT, non-targeting siRNA; si 107, *Nup107* siRNA; si62, *Nup62* siRNA; si98/96, *Nup98/96* siRNA; si153, *Nup153* siRNA; n=109-123.

**Figure S7.**
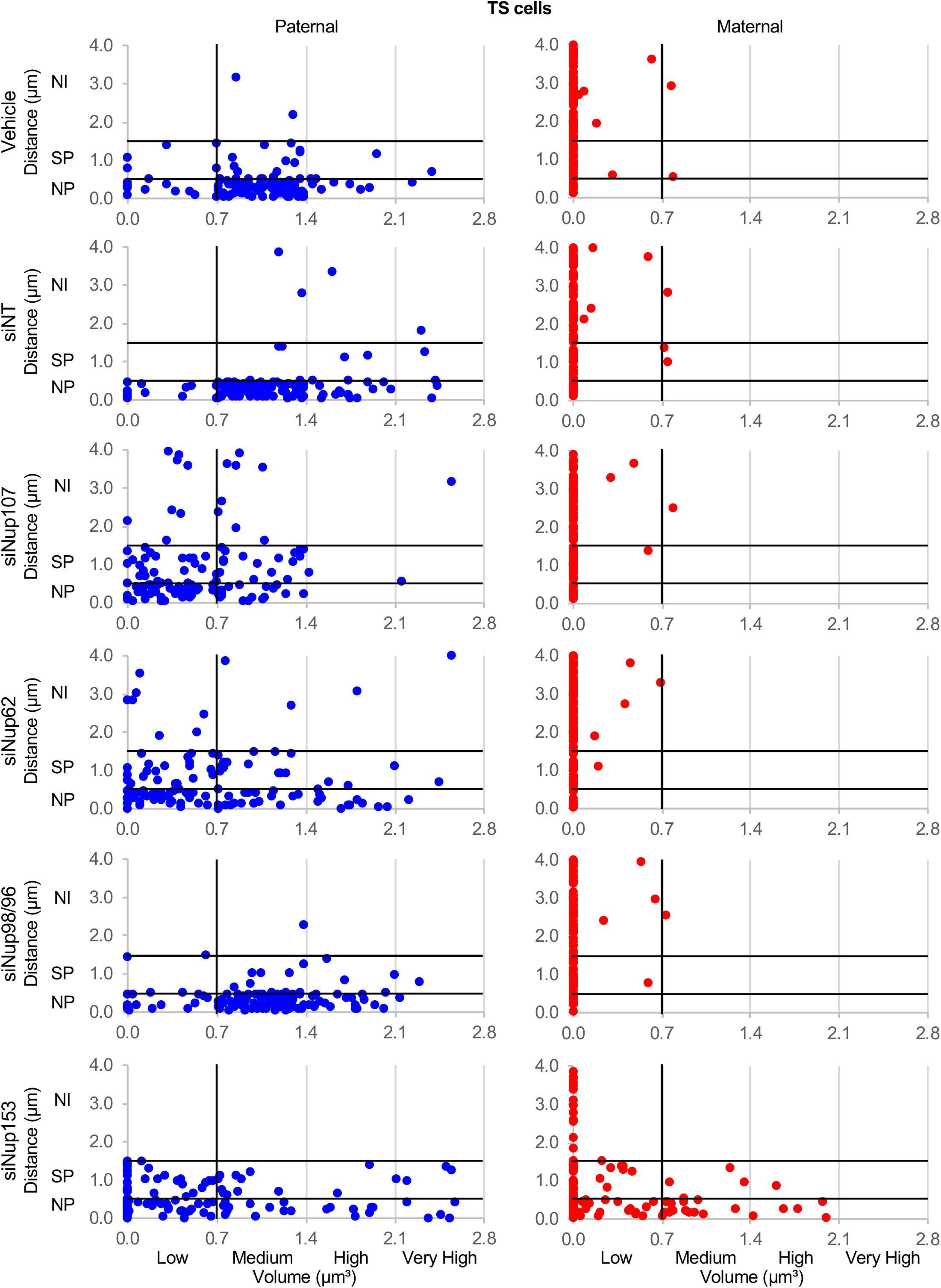
*Kcnq1ot1* ncRNA volume to distance correlation in control and nucleoporin-depleted TS cells. *Kcnq1ot1* ncRNA volume and domain distance from nuclear periphery were plotted on X- and Y-axes, respectively, for G1-synchronized control and *Nup*-*depleted* TS cells. Upon *Nup107, Nup62* and *Nup153* depletion, cells with low paternal *Kcnq1ot1* ncRNA volume (blue) shifted to sub-nuclear peripheral and nuclear interior positions. The maternal *Kcnq1ot1* domain (red) was randomly positioned within the nucleus (expected NP 15%, SP 30%, NI 60%; and observed NP 9–14%, 24–28%; NI, 60–64%), except for Nup153-depleted cells. Upon *Nup153* depletion, those cells with maternal *Kcnq1ot1* ncRNA expression (primarily low) had a shift in maternal *Kcnq1ot1* domain positioning toward the nuclear periphery (observed TS cells NP 52%, 32%, NI, 16%). NP, nuclear periphery; SP, sub-nuclear periphery; NI, nuclear interior; Veh, vehicle; siNT, non-targeting siRNA; si 107, *Nup107* siRNA; si62, *Nup62* siRNA; si98/96, *Nup98/96* siRNA; si 153, *Nup153* siRNA; n=109-123.

**Figure S8.**
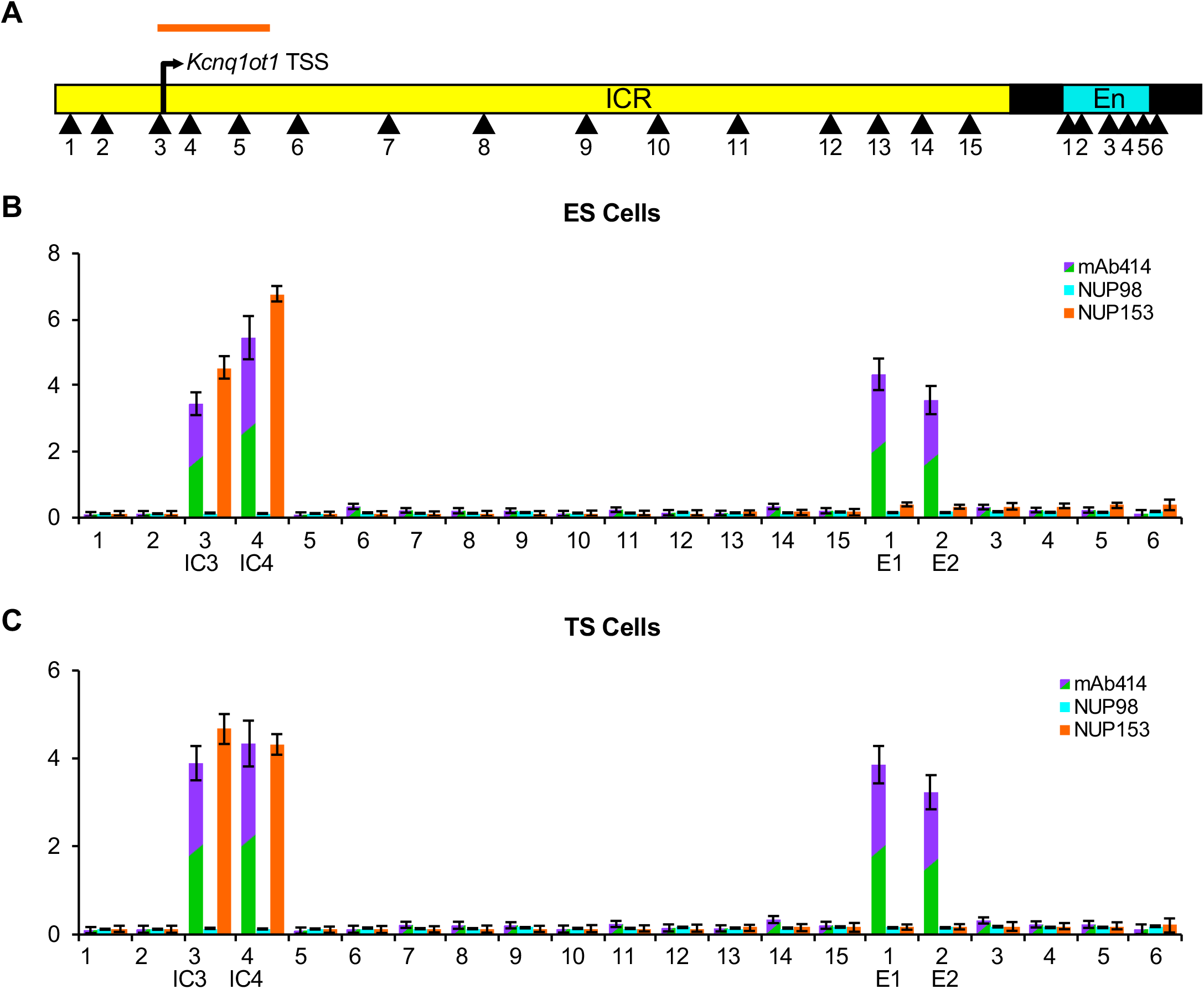
mAb414 and NUP153 enrichment at the *Kcnq1ot1* ICR and enhancer element in ES and TS cells. (A) mAb414 and NUP153 enrichment at 21 sites (arrowheads) examined across the *Kcnq1ot1* ICR and enhancer element. (B, C) Significant enrichment was observed at regions IC3, IC4, E1 and E2 for mAb414, and regions IC3 and IC4 for NUP153 in ES and TS cells. Orange bar represents mapped NUP153 binding site from ES cell DamlD-seq data (Jacinto et. al, 2015). ICR, imprinting control region; En, enhancer element; TSS, transcription start site (n=3 biological samples with 3 technical replicates per sample). Error bars, s.e.m; *, significance p < 0.05 compared to the IgG controls.

**Figure S9.**
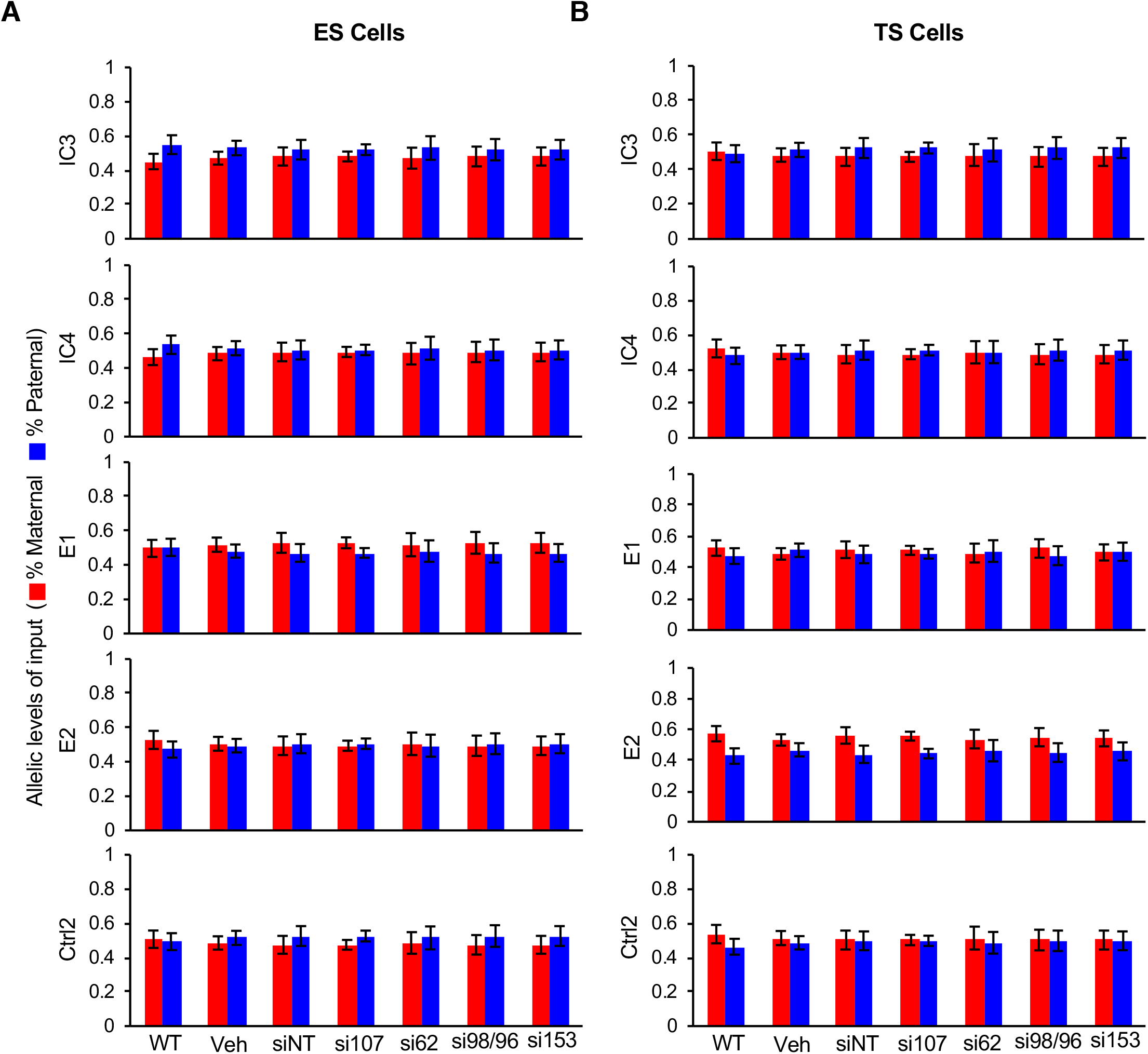
Allelic analysis of ChIP input chromatin from the *Kcnq1ot1* ICR sites in (A) ES and (B) TS cells. Allelic analysis followed by quantification was performed for the maternal and paternal alleles for each primer pair on input chromatin.

**Figure S10.**
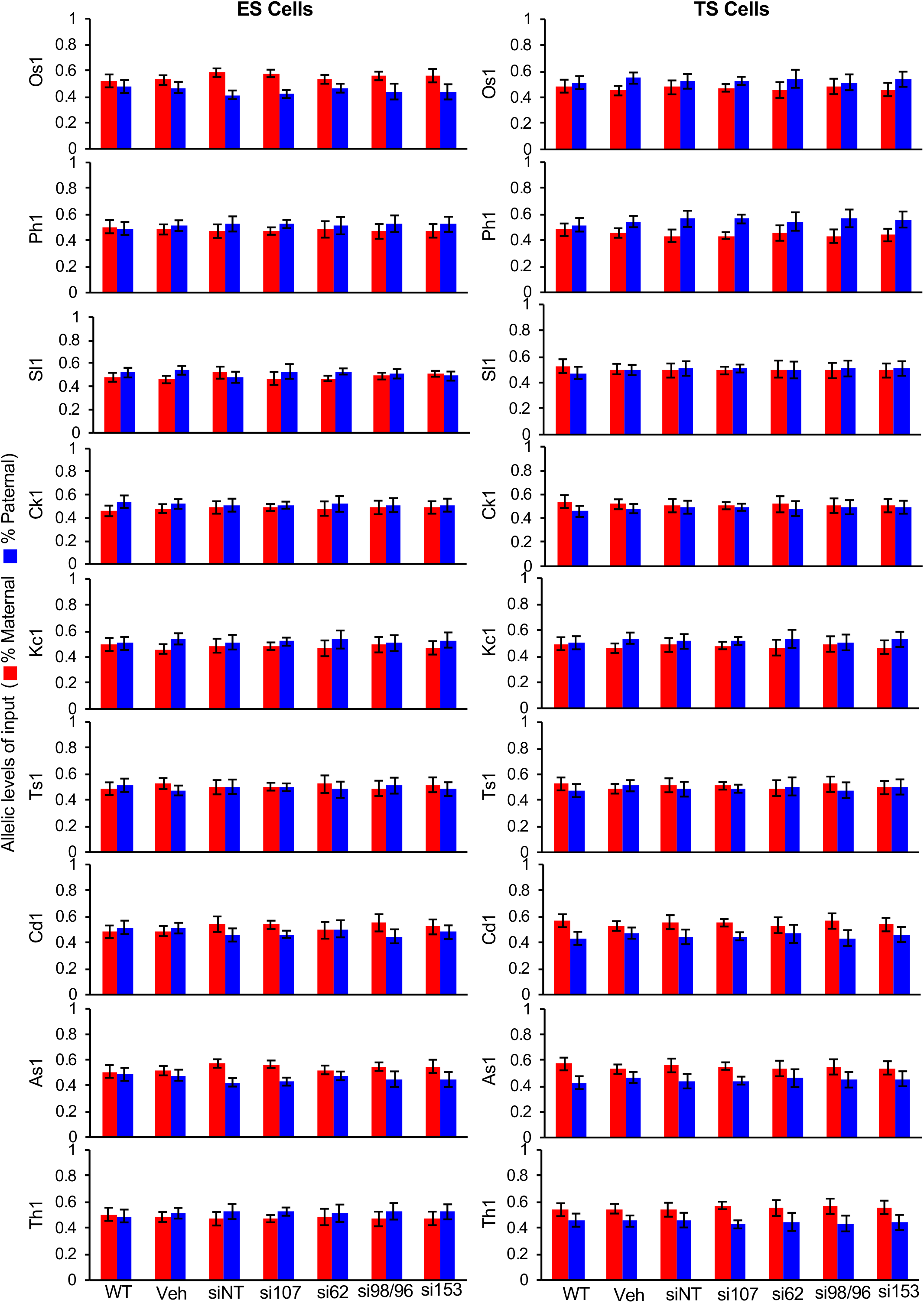
Allelic analysis of ChIP input chromatin for imprinted gene promoters at *Kcnq1ot1* imprinted domain in (A) ES and (B) TS cells. Allelic analysis followed by quantification was performed for the maternal and paternal alleles for each primer pair on input chromatin.

**Figure S11:**
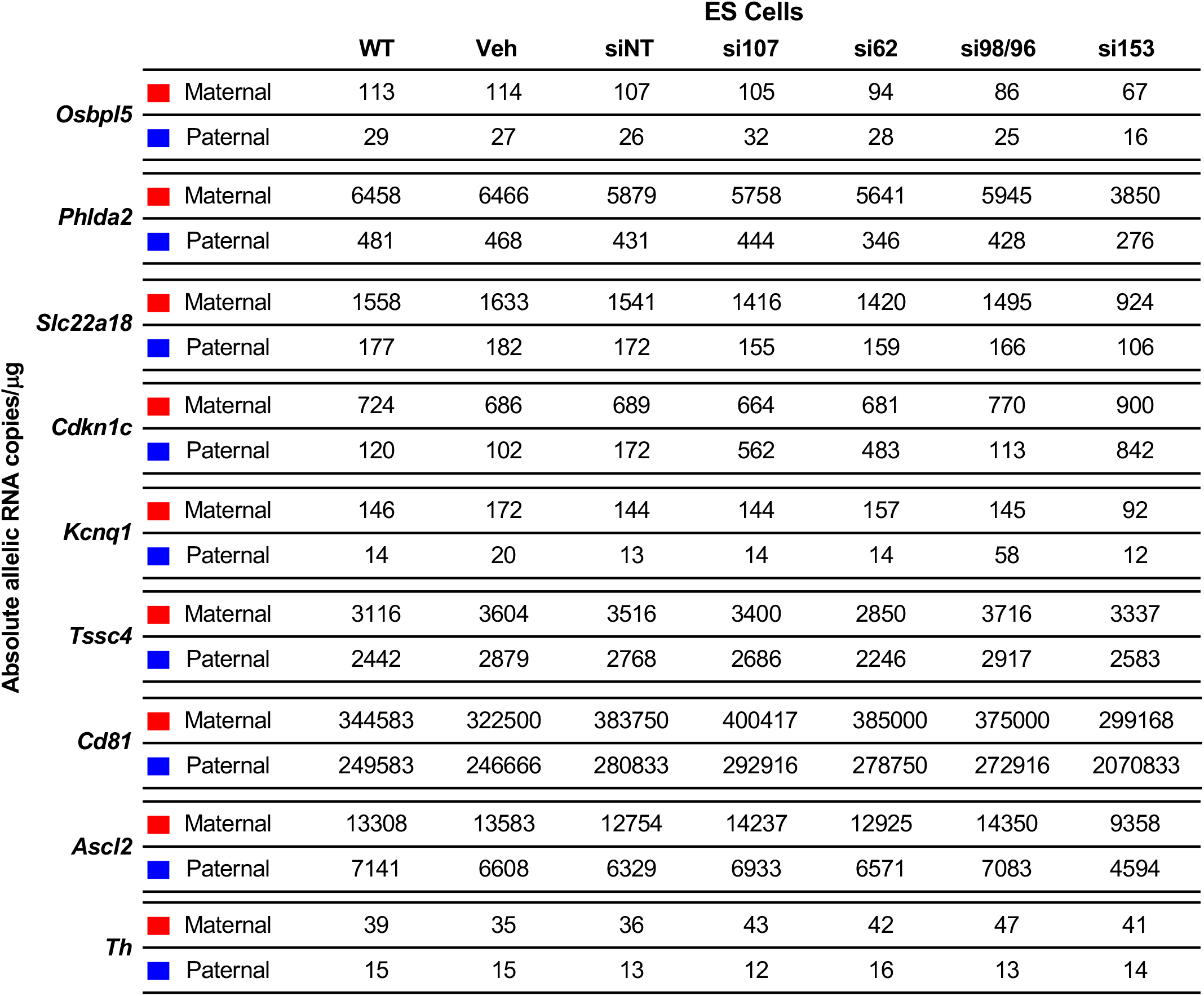
Absolute allelic transcript abundance of imprinted genes in the *Kcnq1* ot1 imprinted domain in ES cells as determined by droplet digital PCR assays. Absolute quantification of B6 and CAST *Kcnq1ot1* RNA copies in 1 of RNA as detected by FAM and HEX probes in control and *Nup*-depleted ES cells (n=3 biological samples) using droplet digital PCR.

**Figure S12:**
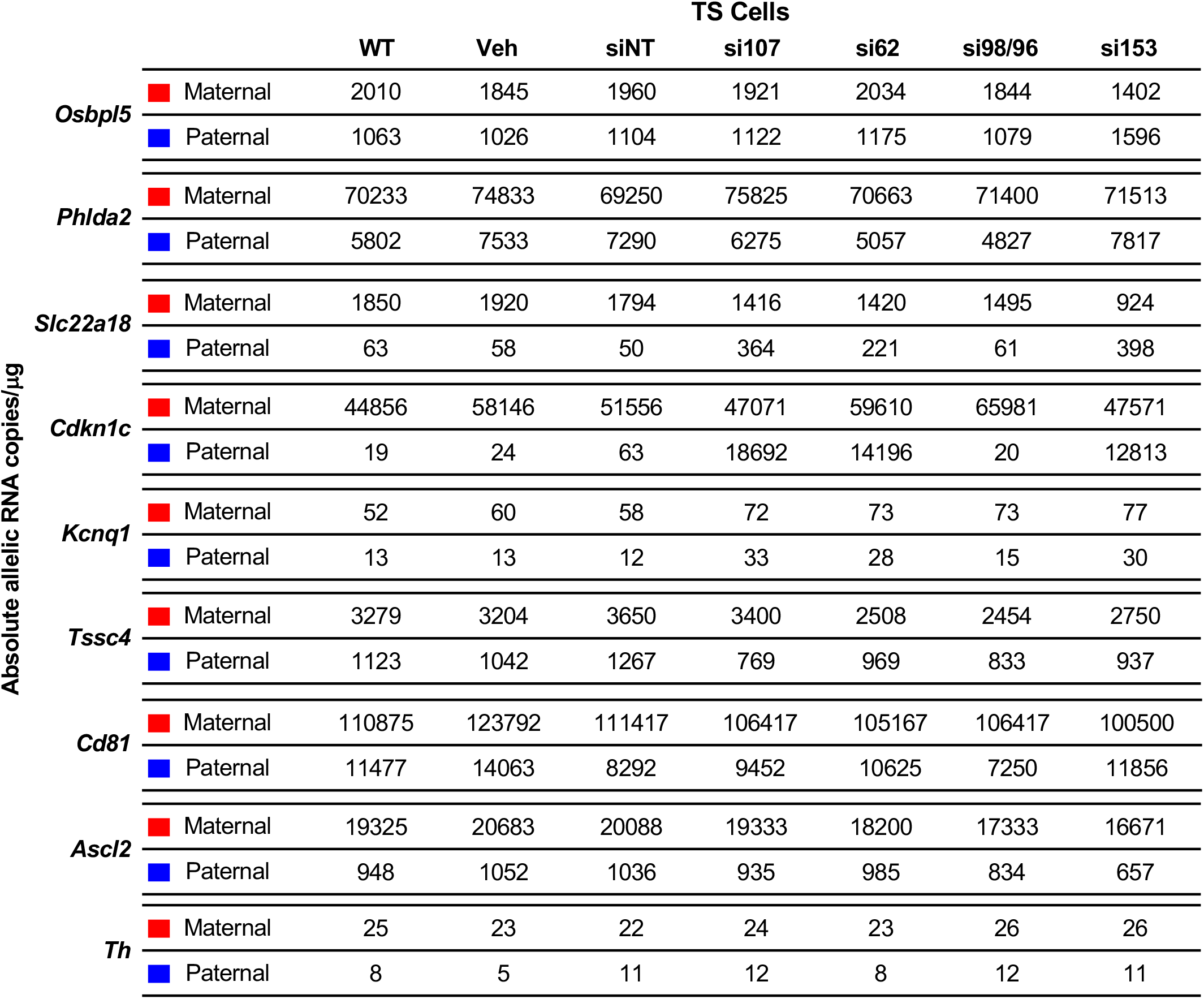
Absolute allelic transcript abundance of imprinted genes in the *Kcnq1ot1* imprinted domain in TS cells as determined by droplet digital PCR assays. Absolute quantification of B6 and CAST *Kcnq1ot1* RNA copies in 1 of RNA as detected by FAM and HEX probes in control and *Nup*-depleted TS cells (n=3 biological samples) using droplet digital PCR.

**Figure S13.**
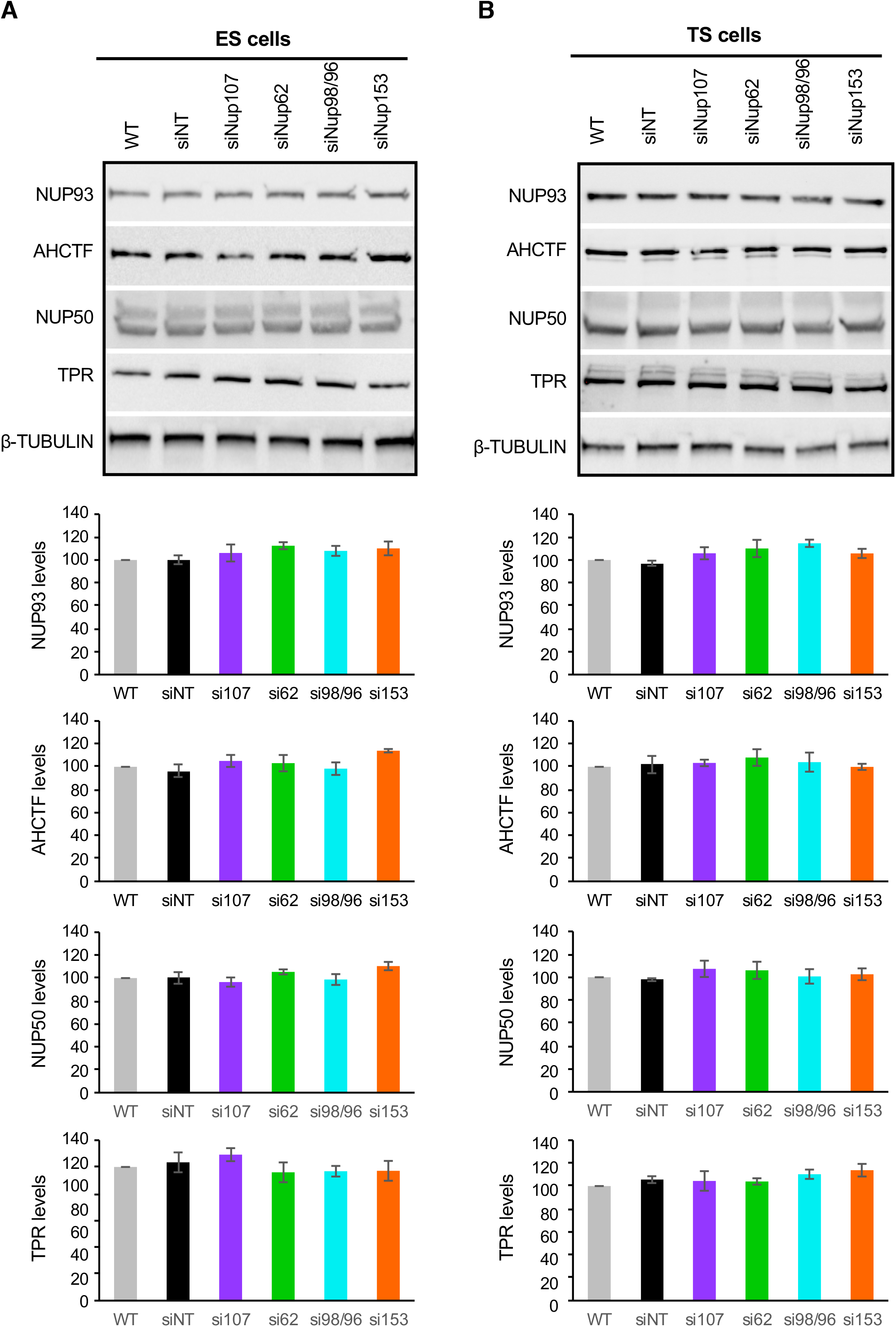
Nucleoporin levels were not disrupted upon *Nup107, Nup62, Nup98/96* or *Nup153* depletion in (A) ES and (B) TS cells. Nucleoporin levels were not altered upon *Nup107, Nup62, Nup98/96* and *Nup153* depletion. Western blot analysis using TPR, AHCTF1 (also known as ELYS), NUP93 and NUP50 antibodies was performed 48 hours after transfection in ES and TS cells. Nucleoporin levels were not significantly different between control and *Nup*-depleted ES and TS nuclear extracts. β–tubulin was used as a loading control. Error bars, s.e.m.; *, significance p < 0.05 compared to the vehicle control; (n=3 biological samples).

**Figure S14.**
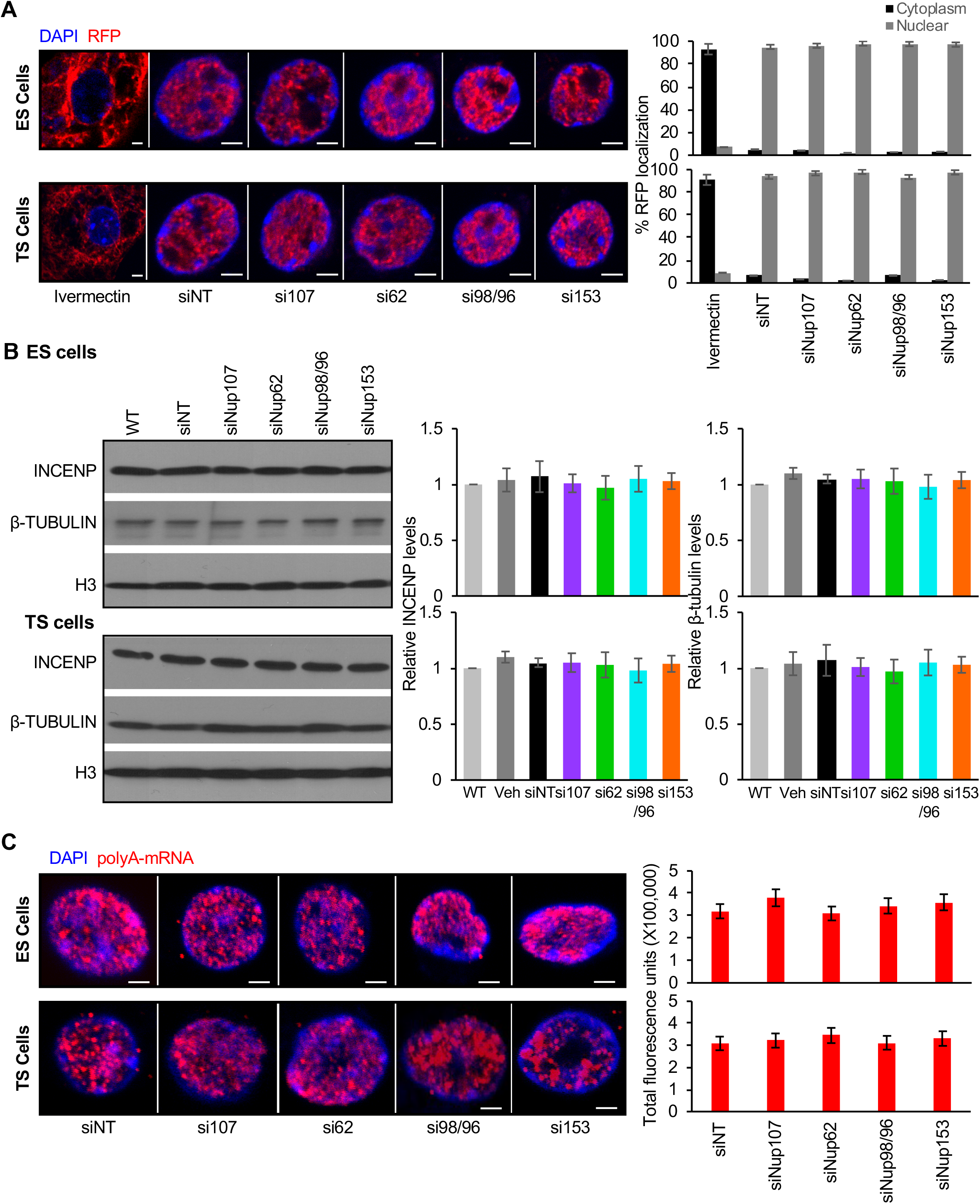
Nuclear transport was not altered upon nucleoporin depletion in ES and TS cells. (A) Endogenous E47-RFP^NLS^ protein transport was not disrupted upon nucleoporin depletion, compared to the ivermectin control where nuclear import was blocked in ES and TS cells. Percent nuclear E47-RFP^NLS^ localization was not significantly different between control and nucleoporin-depleted ES and TS cells, compared to ivermectin control. Scale bar, 1 μm; (n=50). (B) Endogenous protein transport was not disrupted upon nucleoporin depletion. Relative INCENP and β–tubulin protein levels were not significantly different between control and *Nup*-depleted ES and TS nuclear extracts, respectively. Histone 3 (H3) was used as a loading control (n=2-3 biological samples with 3 technical replicates per sample). (C) Nuclear polyA-mRNA retention levels were not changed upon nucleoporin depletion. Compared to controls, nucleoporin-depleted ES and TS cells did not accumulate polyA-mRNA, showing no significant difference in biotin-lablelled oligodT fluorescence levels corrected for background levels. Scale bar, 1 μm; (n=40). Error bars, s.e.m; *, significance p < 0.05 compared to the siNT control; (n=3 biological samples with 3 technical replicates per sample).

**Figure S15.**
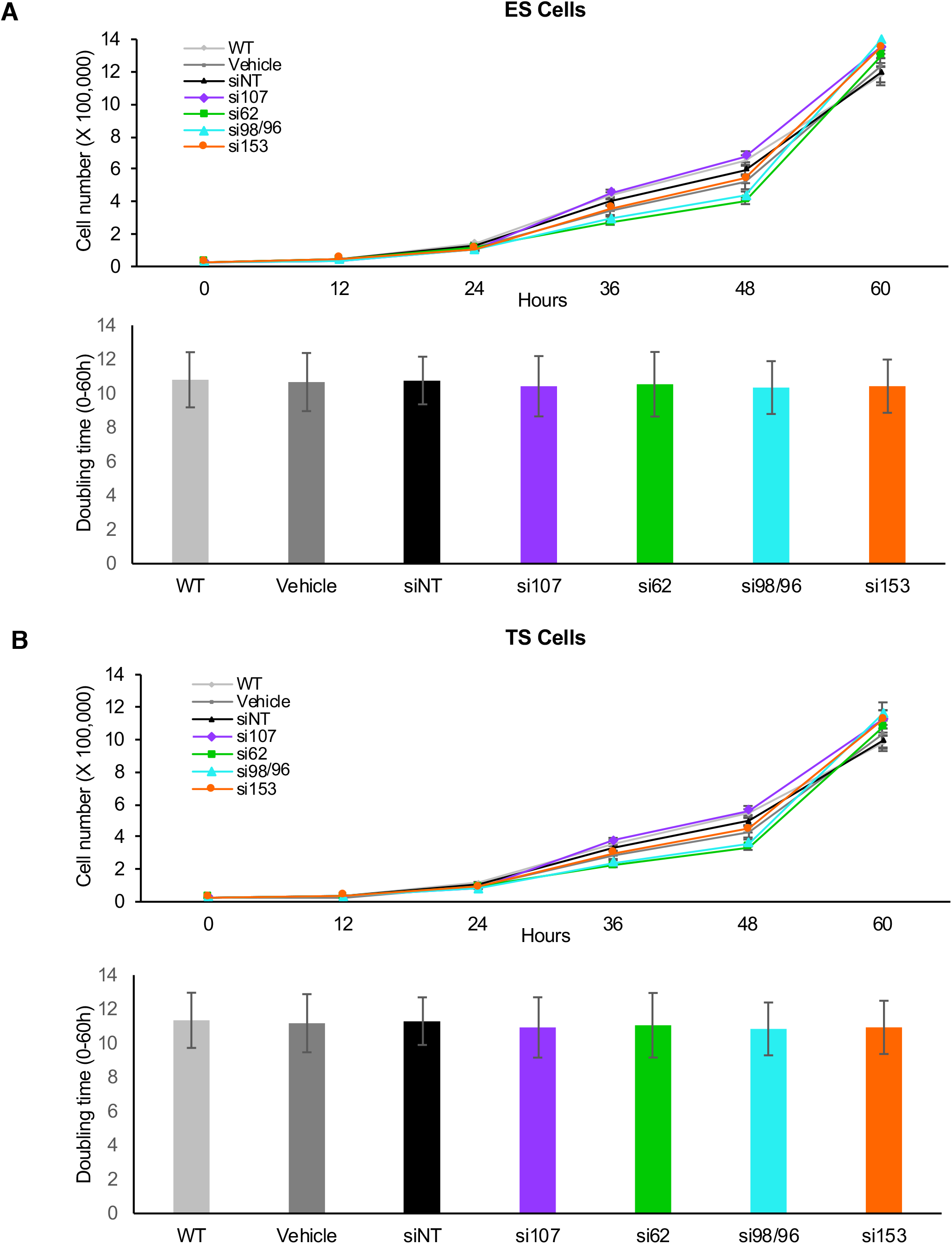
ES and TS cell proliferation was not altered upon nucleoporin depletion. (A) Approximately 25000 cells were seeded and then transfected 12 hours later with siRNAs. Control and *Nup*-depleted ES and TS cells were monitored for 60 hours. Direct cell counts were performed every 12 hours (n=3 biological samples with 3 technical replicates per sample). No significant changes in cell growth rate were observed at different time intervals between samples compared to the WT control. (B) Doubling time (DT) was calculated as specified in the ATCC guidelines using the formula DT = T * [ln (2) / ln (Xe/Xb)], T= time interval; Xe, Cell count at end of time interval; Xb, cell count at start of time interval. No significant change in cell growth rate or doubling time was observed at different time intervals between samples compared to the WT controls. Error bars, s.e.m

**Figure S16.**
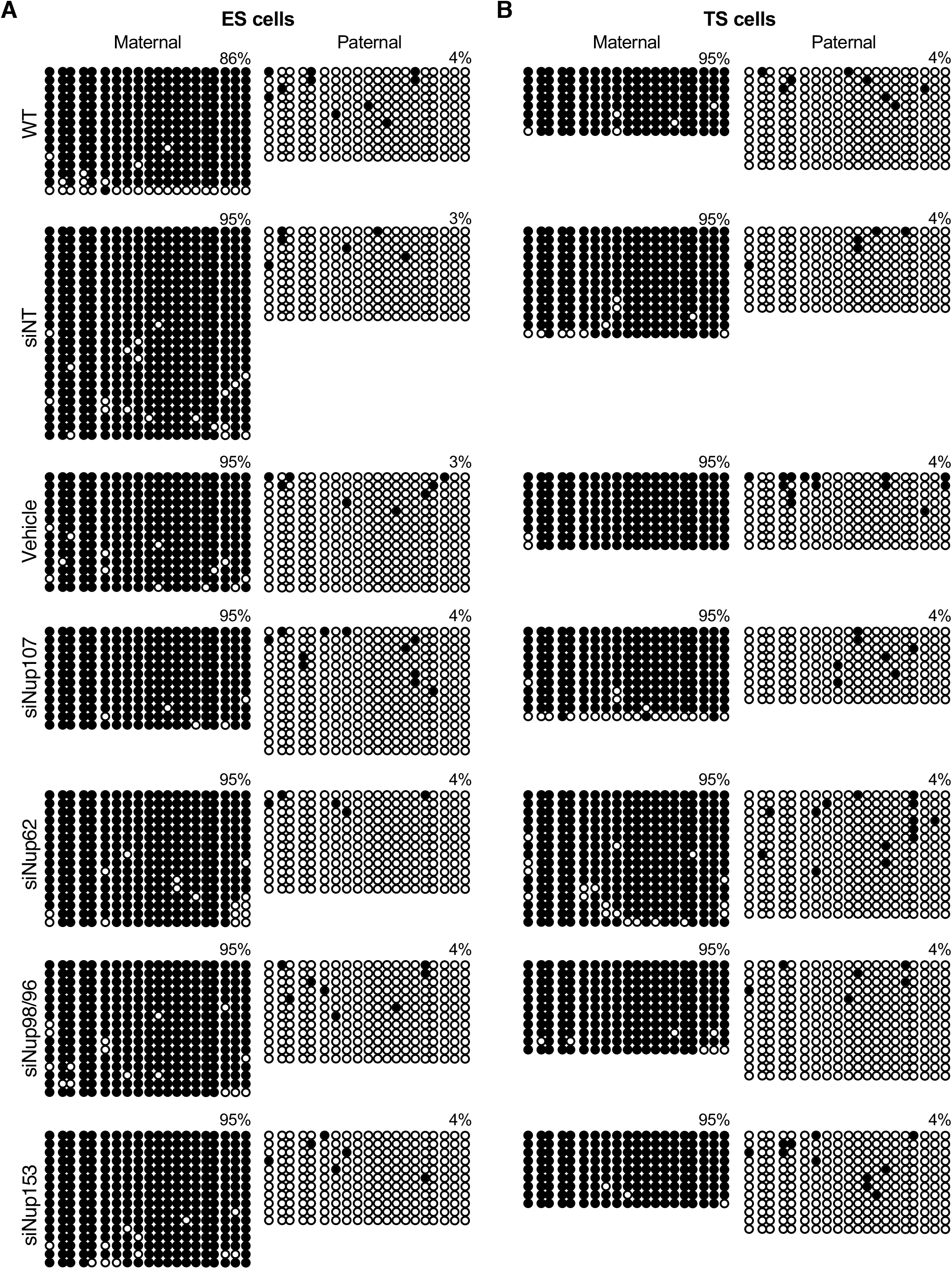
DNA methylation was maintained upon nucleoporin depletion in ES and TS cells. Methylation status of the maternal and paternal *Kcnq1ot1* ICR in control and *Nup*-*depleted* (A) ES and (B) TS cells (one of two biological replicates shown). Black circles, methylated CpGs; white circles, unmethylated CpGs. Each line represents an individual DNA strand. Total methylation percent is represented above each set of DNA strands.

**Figure S17.**
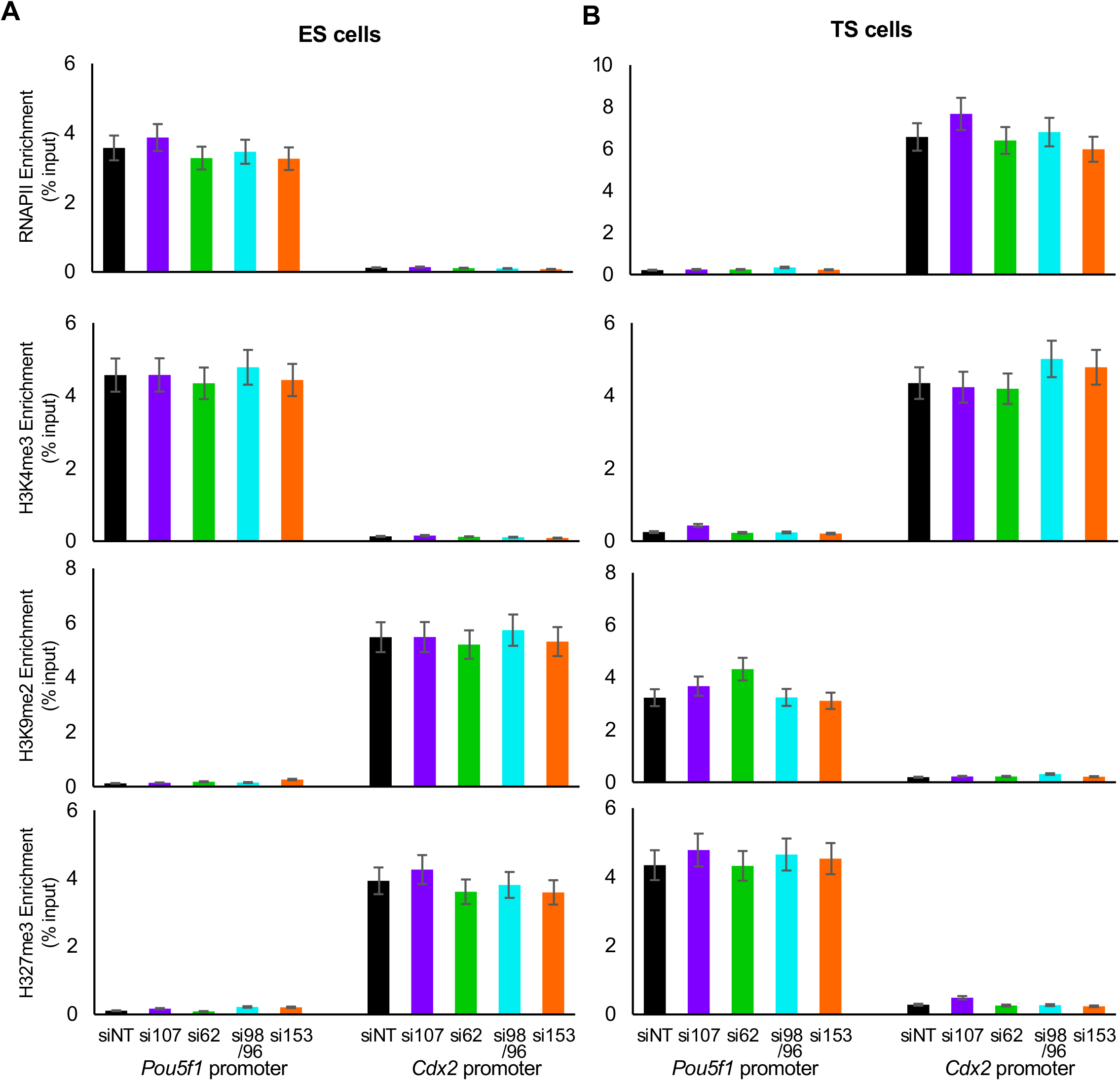
Active and repressive chromatin modifications at the *Pou5f1* and *Cdx2* genes in (A) ES and (B) TS cells, which are unchanged by nucleoporin depletion. From *Nup153* depletion RNA-seq data, Jacinto et al. (2015) observed no change in expression of pluripotency markers in ES cells, including for *Pou5f1 (Oct4)*. Thus, the ES-expressed/TS-repressed marker, *Pou5f1*, as well as ES-repressed/TS-expressed gene, *Cdx2*, were used as positive and negative controls in ES and TS cells, respectively (Lim *et. al*. 2008). ChIP analysis was performed using RNAPII, H3K4me3, H3K9me2 and H3K27me3 antibodies at *Pou5f1* and *Cdx2* promoters. In ES cells, the *Pou5f1* promoter harbored active chromatin modification, while the *Cdx2* promoter was enriched for repressive modifications, as expected. In TS cells, the *Cdx2* promoter harbored active chromatin modification, while the *Pou5f1* promoter was enriched for repressive modifications, as expected. No significant change in enrichment levels were observed upon nucleoporin depletion compared to the siNT control (n=3 biological samples with 3 technical replicates per sample). Error bars, s.e.m; *, significance p < 0.05.

**Figure S18.**
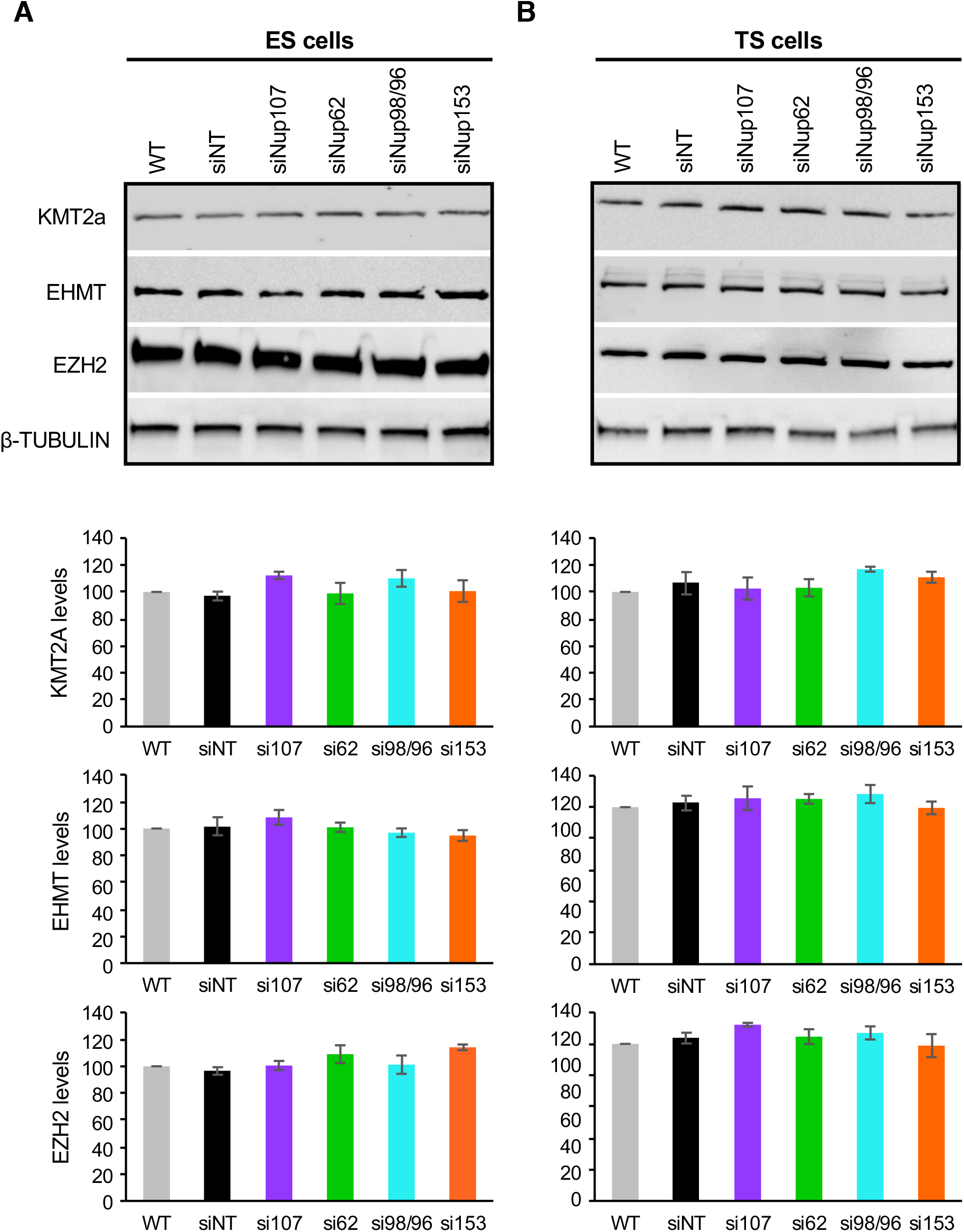
EZH2, KMT2Aand EHMT levels were not changed upon nucleoporin depletion in (A) ES and (B) TS cells. Histone modifier levels were not altered upon *Nup107, Nup62, Nup98/96* or *Nup153* depletion. Western blot analysis using EZH2, KMT2a, and EHMT antibodies was performed 48 hours after transfection. No significant difference was observed between protein levels in control and A/ivp-depleted ES and TS nuclear extracts. β–tubulin was used as a loading control. Error bars, s.e.m.; *, significance p < 0.05 compared to the vehicle control; (n=3 biological samples).

**Figure S19.**
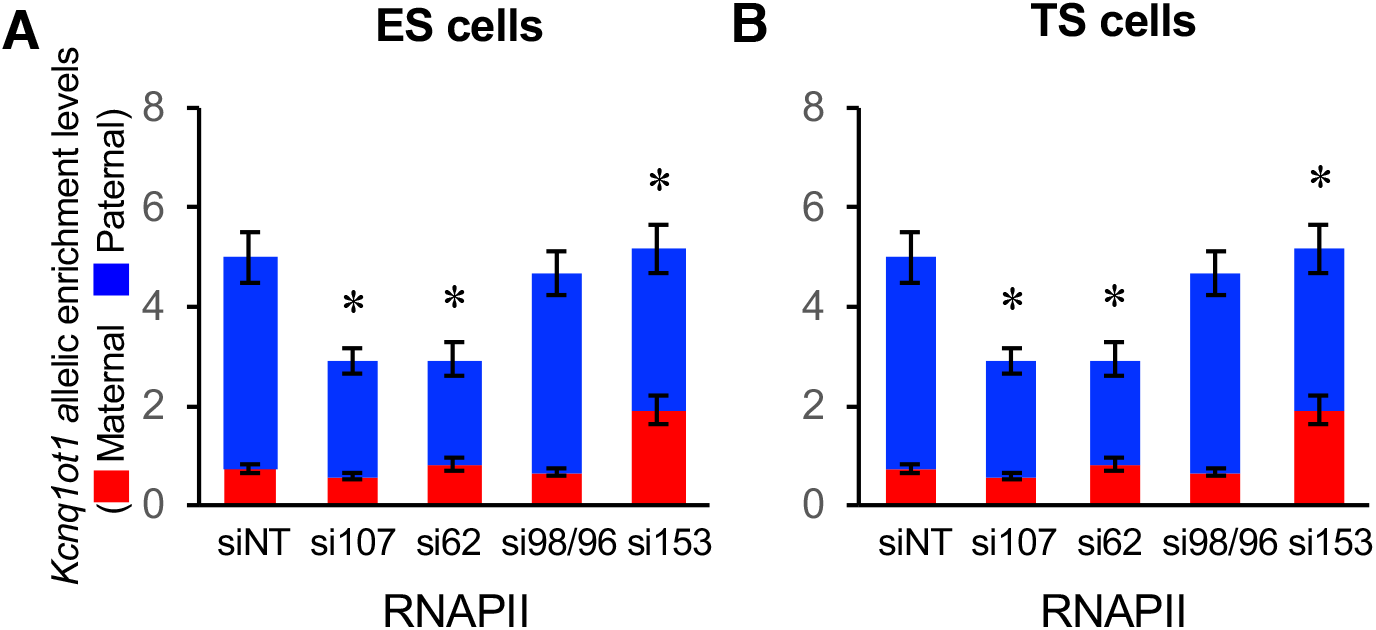
Nucleoporin depletion altered RNAPII enrichment at the *Kcnq1ot1* ICR upon *Nup* depletion in (A) ES and (B) TS cells. RNAPII ChIP at the maternal and paternal *Kcnq1ot1* ICR in control and *Nup*-depleted ES and TS cells (n=3 biological samples with 3 technical replicates per sample). Allelic proportions are represented as a percent of the total enrichment level. WT, wildtype; Veh, vehicle; siNT, non-targeting siRNA; si107, *Nup107* siRNA; si62, *Nup62* siRNA; si98/96, *Nup98/96* siRNA, si153, *Nup153* siRNA. error bars, s.e.m; *, significance p < 0.05 compare to the siNT control.

**Figure S20.**
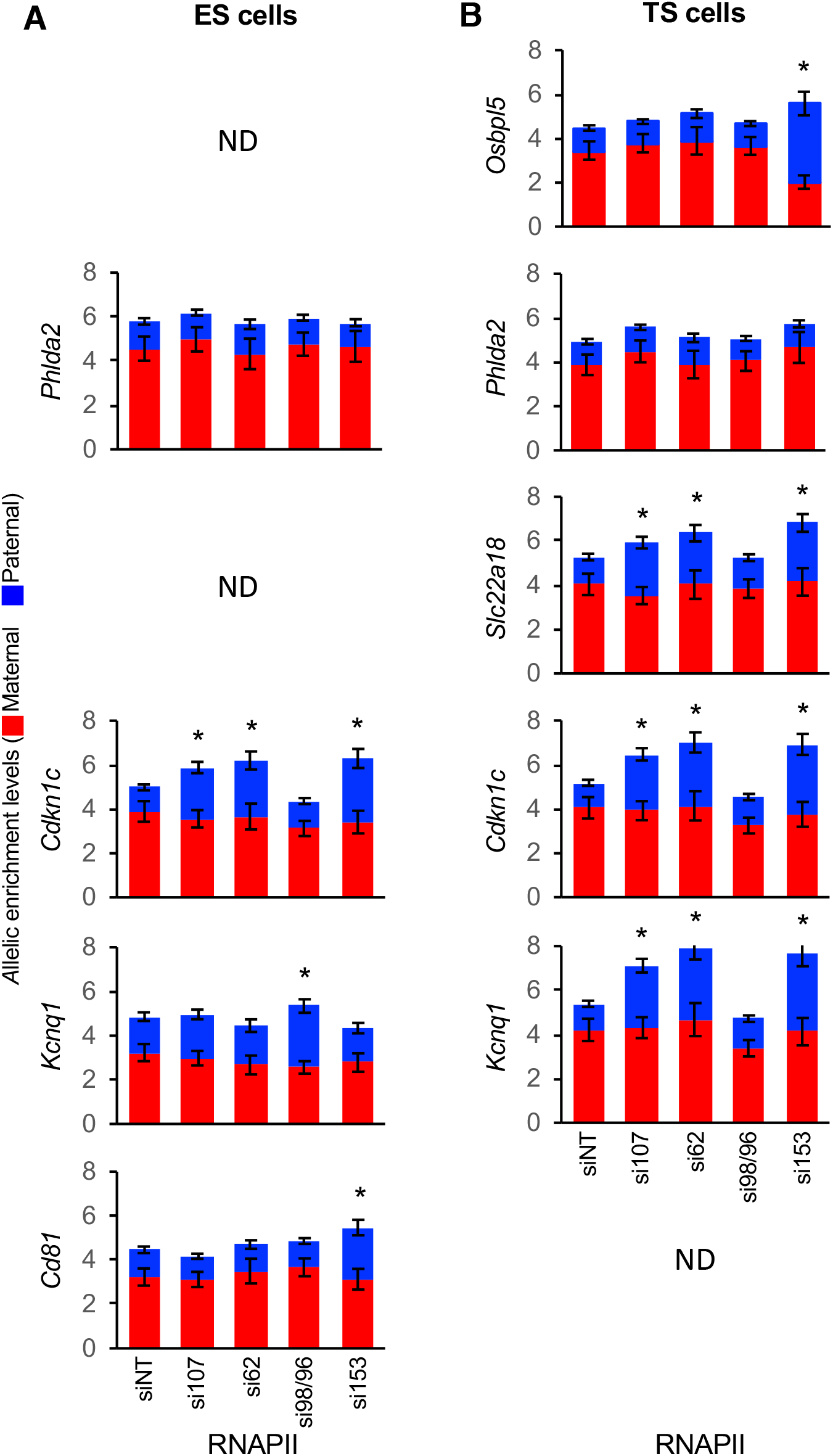
Nucleoporin depletion disrupted RNAPII enrichment at reactivated imprinted gene promoters but not at *Phlda2* in ES and TS cells. RNAPII, ChIP at the maternal and paternal imprinted gene promoters in control and *Nup*-depleted ES and TS cells (n=3 biological samples with 3 technical replicates per sample). Allelic proportions are represented as a percent of the total enrichment level. ND, not determined; WT, wildtype; Veh, vehicle; siNT, non-targeting siRNA; si 107, *Nup107* siRNA; si62, *Nup62* siRNA; si98/96, *Nup98/96* siRNA, si 153, *Nup153* siRNA. error bars, s.e.m; *, significance p < 0.05 compare to the siNT control.

**Figure S21.**
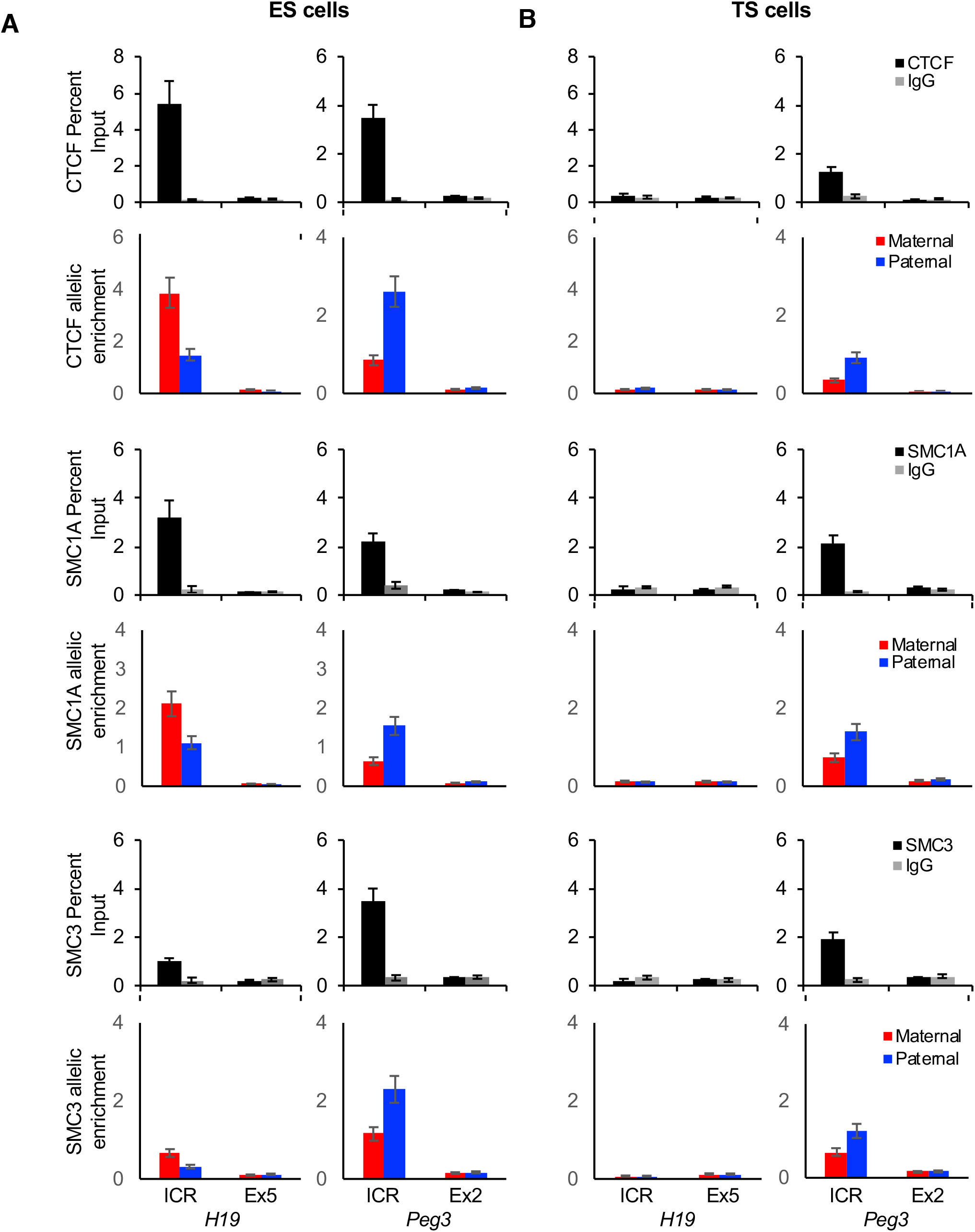
CTCF, SMC1A and SMC3 enrichment at the maternal *H19* ICR and paternal *Peg3* DMR in ES cells and at the *Peg3* DMR in TS cells. CTCF, SMC1A and SMC3 ChIP at the *H19* ICR and *Peg3* DMR in wildtype ES and TS cells (n=3 biological samples with 3 technical replicates per sample). Allelic analysis indicated maternal enrichment for CTCF, SMC1A and SMC3 at *H19* ICR and paternal enrichment for CTCF, SMC1A and SMC3 at the *Peg3* DMR. Error bars, s.e.m.

**Figure S22.**
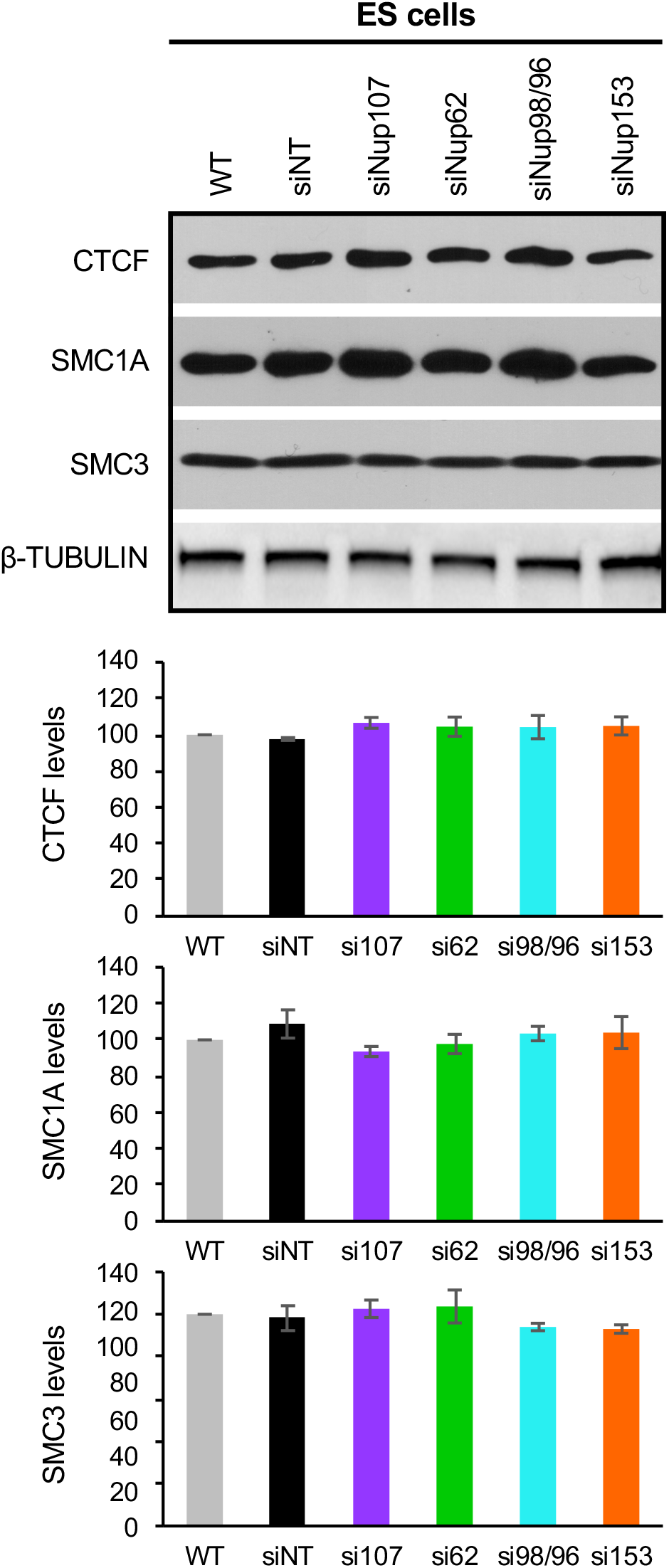
CTCF, SMC1A and SMC3 levels were unchanged upon nucleoporin depletion in ES cells. Western blot analysis using CTCF, SMC1A, and SMC3 antibodies was performed 48 hours after transfection. No significant difference was observed between protein levels in control and *Nup*-depleted ES nuclear extracts. (3-tubulin was used as a loading control. Error bars, s.e.m.; *, significance p < 0.05 compared to the vehicle control; (n=3 biological samples).

**Figure S23.**
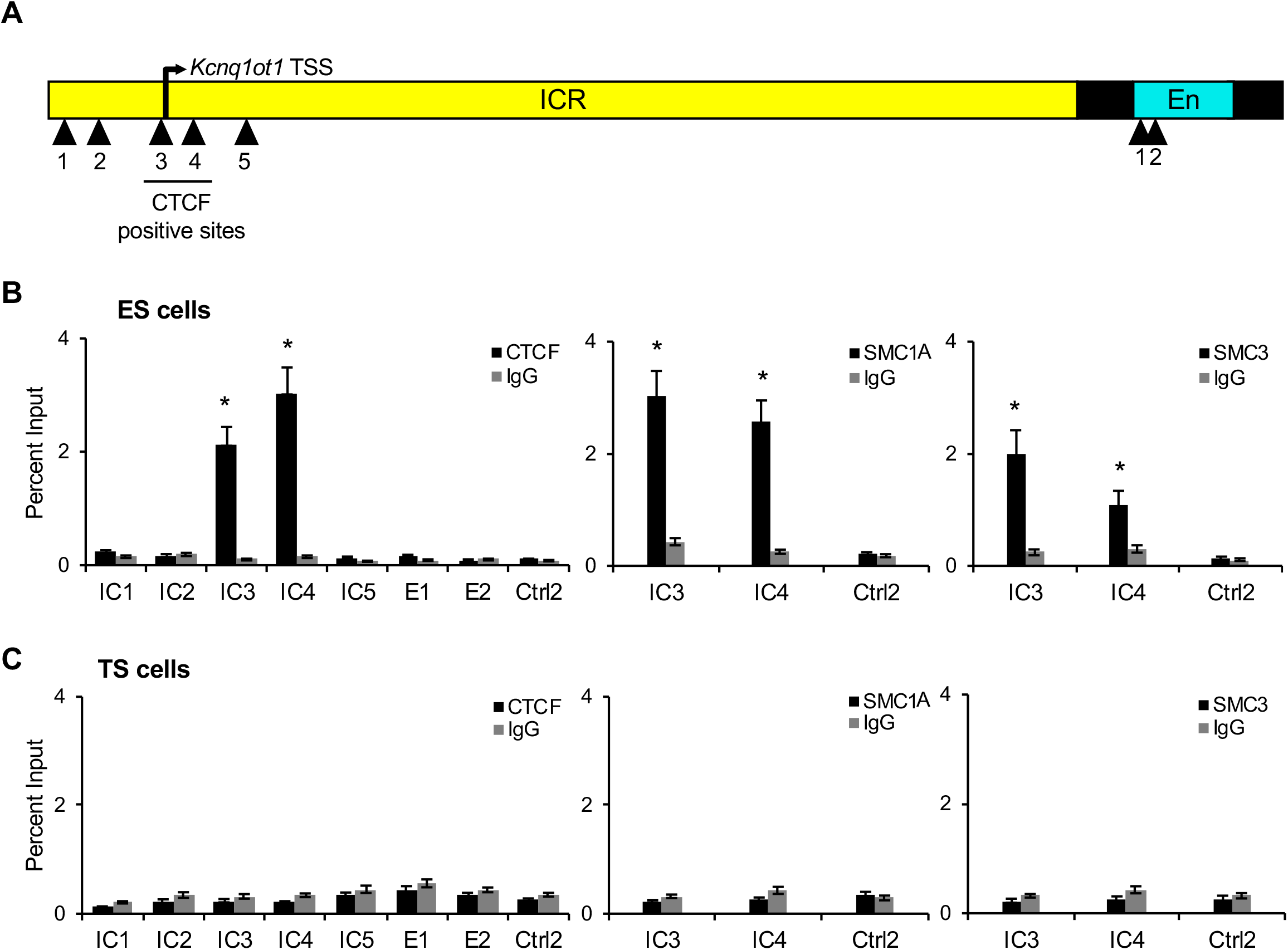
CTCF, SMC1A and SMC3 enrichment at the *Kcnq1ot1* ICR and enhancer element sites in ES and TS cells. (A) Map of the primer test sites for *Kcnq1ot1* ICR and enhancer region. (B). In ES cells CTCF, SMC1A and SMC3 enrichment was seen at the IC3 and IC4 regions. (C). No CTCF, SMC1A and SMC3 enrichment was observed at sites within the *Kcnq1ot1* ICR and enhancer region in TS cells. Error bars, s.e.m; *, significance p < 0.05 compared to IgG control; (n=3 biological samples with 3 technical replicates per sample).

**Figure S24:**
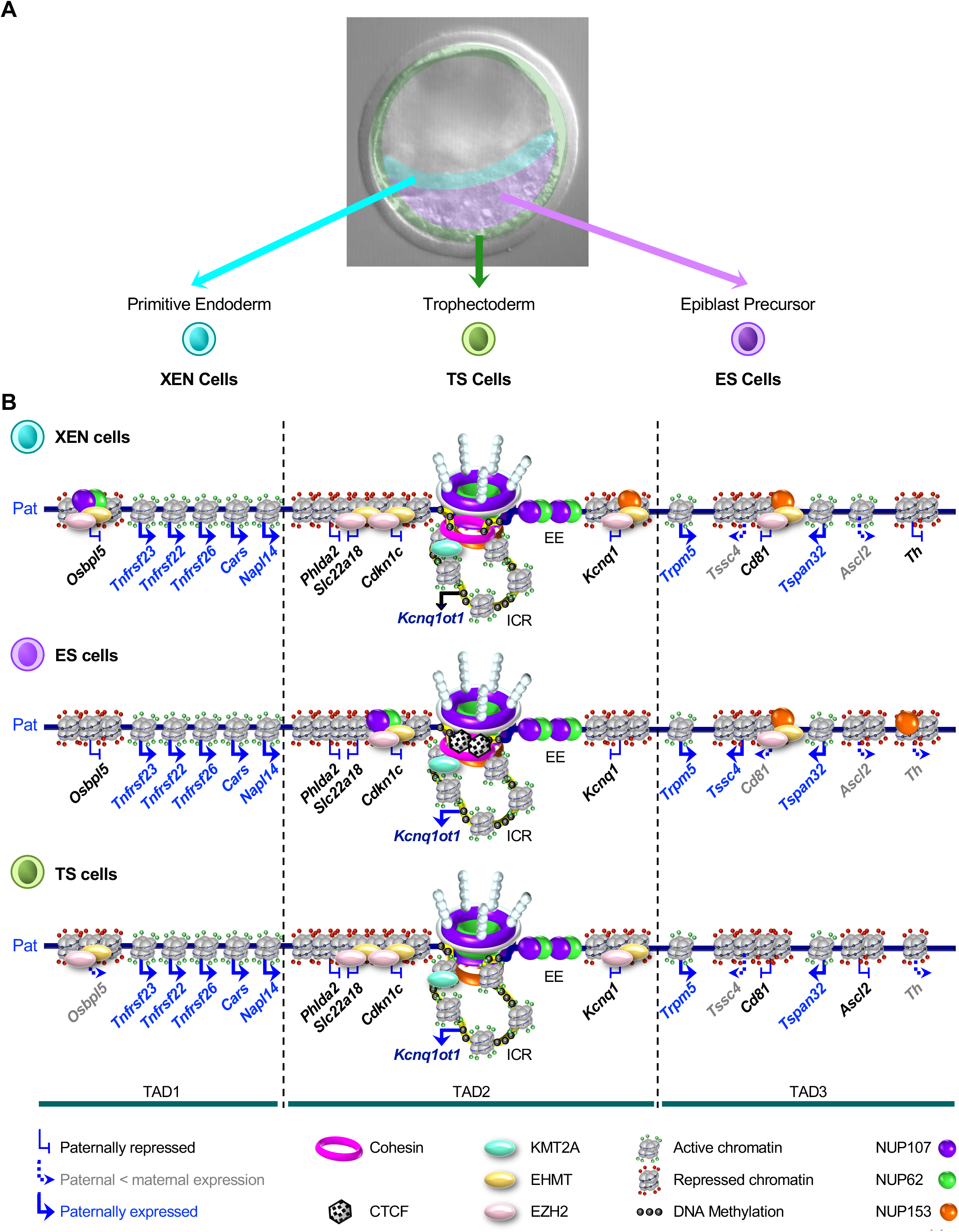
Model for NUP107, NUP62 and NUP153-mediated regulation of the paternal *Kcnq1ot1* imprinted domain in XEN, ES and TS cells. (A) In blastocyst stage embryos, there are three cell lineages, epiblast precursor, trophectoderm and primitive endoderm cells, which can be used to derive embryonic (ES), trophoblast (TS) and extraembryonic endoderm (XEN) stem cells. (B) NUP107, NUP62 and NUP153-mediated regulation of the paternal *Kcnq1ot1* imprinted domain in XEN, ES and TS cells. The *Kcnq1ot1* ncRNA is expressed primarily from the paternal allele (navy text). Genes in black text are paternally silent, while genes in grey text show less expression from the paternal (<35%) than the maternal (>65%) allele. Genes in blue text are expressed biallelically from the paternal and maternal alleles. In TAD2, NUP107, NUP62 and NUP153 within the nuclear pore complex together with cohesin in XEN cells, cohesin and CTCF in ES cells, and neither in TS cells interact with *Kcnq1ot1* ICR on both sides of the *Kcnq1ot1* promoter, generating boundaries that demarcated active chromatin at the paternal *Kcnq1ot1* ICR from flanking repressive chromatin. NUP107/NUP62 occupancy at the paternal enhancer element (EE) in all lineages may also generate an additional insulator/boundary that isolates the enhancer from the repressed paternal *Kcnq1* allele. In ES cells, NUP107/NUP62 are also enriched at the paternal *Cdkn1c* promoter region, likely separating paternal *Cdkn1c* silencing from the other upstream paternal alleles. In TAD1, NUP107/NUP62 binding at the paternal *Osbpl5* promoter in XEN and TS cells could delineate repressive chromatin at *Osbpl5* from active chromatin at neighbouring non-imprinted genes. Similarly, in TAD3 of XEN cells, NUP153 occupancy at the paternal *Cd81* promoter may separate repressive chromatin at *Cd81* from active neighbouring non-imprinted genes. In TAD 3 of ES cells, NUP153 enrichment at the paternal *Cd81* and *Th* promoters may mark repressive from active chromatin. Histone modifiers that show NUP-dependent enrichment at the *Kcnq1ot1* ICR and imprinted gene promoters are indicated.

